# Amino acid homorepeats act as buffers to maintain proteostasis and constrain the compatible sequence space of proteomes

**DOI:** 10.64898/2025.12.19.695406

**Authors:** Yukihiro Murase, Naoki Kitamura, Shotaro Namba, Ayano Satoh, Takashi Makino, Ayako Moriya, Hisao Moriya

## Abstract

Proteins are composed of combinations of 20 amino acids, yet the principles governing their sequence compatibility within living cells remain poorly understood. Here, we evaluated the “neutrality”—an integrated measure of harmful versus beneficial properties—of amino acid homorepeats (PolyX) in *Saccharomyces cerevisiae*. Using the genetic tug-of-war (gTOW) method to quantify maximum expression limits, we found that neutrality is highly amino-acid-specific and intrinsically linked to physicochemical properties. Hydrophobic repeats (I, V, W, F, Y) were universally harmful and aggregation-prone, while hydrophilic repeats (E, S, N, Q) not only exhibited high neutrality but also actively suppressed the proteotoxic stress and aggregation of host proteins. These experimental neutrality values show a striking correlation with the occurrence and length of homorepeats across diverse eukaryotic proteomes. Our findings suggest that PolyX neutrality defines the “compatible sequence space” of proteins, where certain sequences like Poly_10_E act as evolutionary buffers to maintain proteostasis. This study provides a quantitative framework for understanding how physical constraints on protein solubility shape the evolution of proteomes and offers a new metric for rational protein design.

## Introduction

Most proteins produced by extant organisms are composed of combinations of 20 amino acids with diverse physicochemical properties. The possible combinations of amino acid sequences—and thus their physicochemical characteristics—are virtually infinite. Among them, sequences that are compatible with living organisms have been selectively retained through the course of evolution ^1^. This compatibility is determined by a balance between harms and benefits to biological functions, where the harms include various factors that decrease fitness, such as cytotoxicity or excessive consumption of cellular resources ^2^.

One effective way to understand the compatibility of amino acid sequences is to identify patterns commonly present in amino acid sequences (natural sequences) used by extant organisms. With the vast amount of protein sequence data now available, attempts have been made to extract features of natural sequences using artificial intelligence and to apply them to three-dimensional structure prediction and de novo protein design ^3–6^. Although this approach has achieved success in extracting, predicting, and designing functional sequences, it does not directly reveal the compatibility of sequences that do not exist in nature, nor the characteristics of sequences that are not compatible due to harmful effects. “Nullpeptides,” which are peptides absent from the proteome of a given organism, may represent sequences that have been eliminated during evolution ^7,8^. However, such inferences still depend on the limited sequence space of natural proteins, and thus have inherent limitations in capturing the full landscape of non-compatible sequences.

Another approach to investigating sequence compatibility is the synthetic exploration of compatible combinations among amino acid sequences. By constructing a comprehensive sequence library and performing functional screening, it is possible to obtain functional sequences that do not exist in nature ^9–11^. Furthermore, the use of Deep Mutational Scanning (DMS) enables systematic and quantitative evaluation of sequences within the library, allowing the identification of both the diversity and the characteristics of compatible sequences ^12,13^. The resulting data can also serve as training datasets for machine learning, and with sufficient data density, highly accurate predictive models can be developed ^14,15^. Neme *et al.* demonstrated through DMS in *Escherichia coli* that, while many random sequences are harmful, a considerable number of beneficial sequences also exist ^16,17^. However, this exploration still covers only a limited portion of the vast sequence space, and the specific features and physiological effects of harmful or beneficial sequences remain largely unknown.

Therefore, in this study, we aimed to extract harmful and beneficial features from extremely simple model sequences. Of particular interest are continuous stretches of identical amino acids, known as amino acid homorepeats ^18–24^ or PolyX. PolyX sequences are also referred to as homopolymeric amino acid stretches/repeats ^25–29^, amino acid runs ^30^, or homopeptide repeats ^31^, and they are present in a wide range of natural proteins. PolyX motifs exhibit distinct physiological functions and have been exploited in various biotechnological applications ^19,25,32^. For example, PolyP (∼10 residues) segments of formin act as nucleation sites for actin filament formation ^33^, and the arginine-rich region of protamine containing PolyR tracts (∼6 residues) binds strongly to DNA, inducing chromatin condensation ^34^. Translation of the poly(A) tail resulting from termination failure produces PolyK, which serves as a signal for protein degradation ^35,36^. The metal-chelating property of PolyH (∼6 residues) is utilized as a purification tag ^37^, while the membrane-penetrating ability of PolyR (∼8 residues) is exploited for molecular delivery ^38^. Conversely, PolyX sequences can also exert harmful effects and are associated with diseases. Abnormal expansion of PolyQ beyond ∼40 residues leads to aggregate formation and neurodegenerative disorders ^39^. Positively charged PolyR and PolyK (∼10 residues) interact with the negatively charged ribosomal tunnel, causing translational stalling ^40^, and PolyW (∼10 residues) induces ribosome stalling that prevents CAT-tail addition ^41^. Negatively charged PolyD and PolyE (∼10 residues) destabilize ribosomes and induce translational arrest ^42,43^.In natural protein sequences, both the frequency and the length of PolyX motifs show amino acid–specific biases ^18,29,31,44^, which are thought to reflect a balance between harmful and beneficial effects. PolyX sequences are considered evolutionary hotspots, as they can readily arise through mutations ^29,45–47^. When present in natural proteins, they may confer benefits, whereas their absence could indicate harms that led to their elimination. Although the cytotoxicity of certain PolyX sequences has been examined in mammalian cells ^26,27,48^, general trends that transcend species and expression conditions remain unexplored.

In this study, we systematically evaluated the harmful and beneficial effects of peptide sequences composed of ten identical amino acids (Poly_10_X) by individually overexpressing them in yeast. A length of ten residues was chosen as it is sufficiently long to manifest amino-acid-specific physicochemical effects, such as charge accumulation or hydrophobic interactions, while remaining short enough to minimize sequence context and secondary-structure complexity. By fixing sequence length and repeating a single residue, this approach allows direct comparison of how intrinsic physicochemical properties—such as charge and hydrophobicity—affect cellular tolerance. Moreover, comparison with PolyX occurrence in natural proteomes enables us to infer which sequence features are favored or disfavored by evolutionary selection, thereby testing whether their harmful or beneficial effects contribute to determining their abundance in extant proteomes.

## Results

### The harmful effects of each Poly_10_X are generally conserved across species

In this study, we began by investigating the expression limits of enhanced green fluorescent protein (EGFP) fused at its C-terminus with 20 different Poly_10_X peptides (Figure 1A, S1, and S2). The fusion was made at the C-terminus rather than the N-terminus to minimize the influence of N-terminal sequences, which are known to strongly affect translation efficiency ^49^. For harmfulness assessment in yeast, we expressed EGFP and EGFP-Poly_10_X from the strong constitutive *TDH3* promoter (*TDH3_pro_*) and increased plasmid copy numbers using the genetic tug-of-war (gTOW) method ^50,51^. During this process, we measured both growth inhibition and EGFP expression levels (cellular fluorescence). In the gTOW assay, plasmid copy number can be increased by culturing cells with or without leucine supplementation. Under both conditions, we evaluated cell growth and fluorescence to calculate the maximum growth rate (MGR) and maximum fluorescence intensity (MFI). To enable one-dimensional comparison of the harmfulness of each Poly_10_X variant, we defined relative neutrality based on growth and expression data as follows: Relative neutrality = %(MGR_Poly_10_X / MGR_Δ) × %(MFI_Poly_10_X / MFI_Δ). Where, Δ refers to the control EGFP without any Poly_10_X. Thus, the relative neutrality values were normalized such that the control (Δ) equals 10,000 (Figure 1A) .

**Figure 1.**
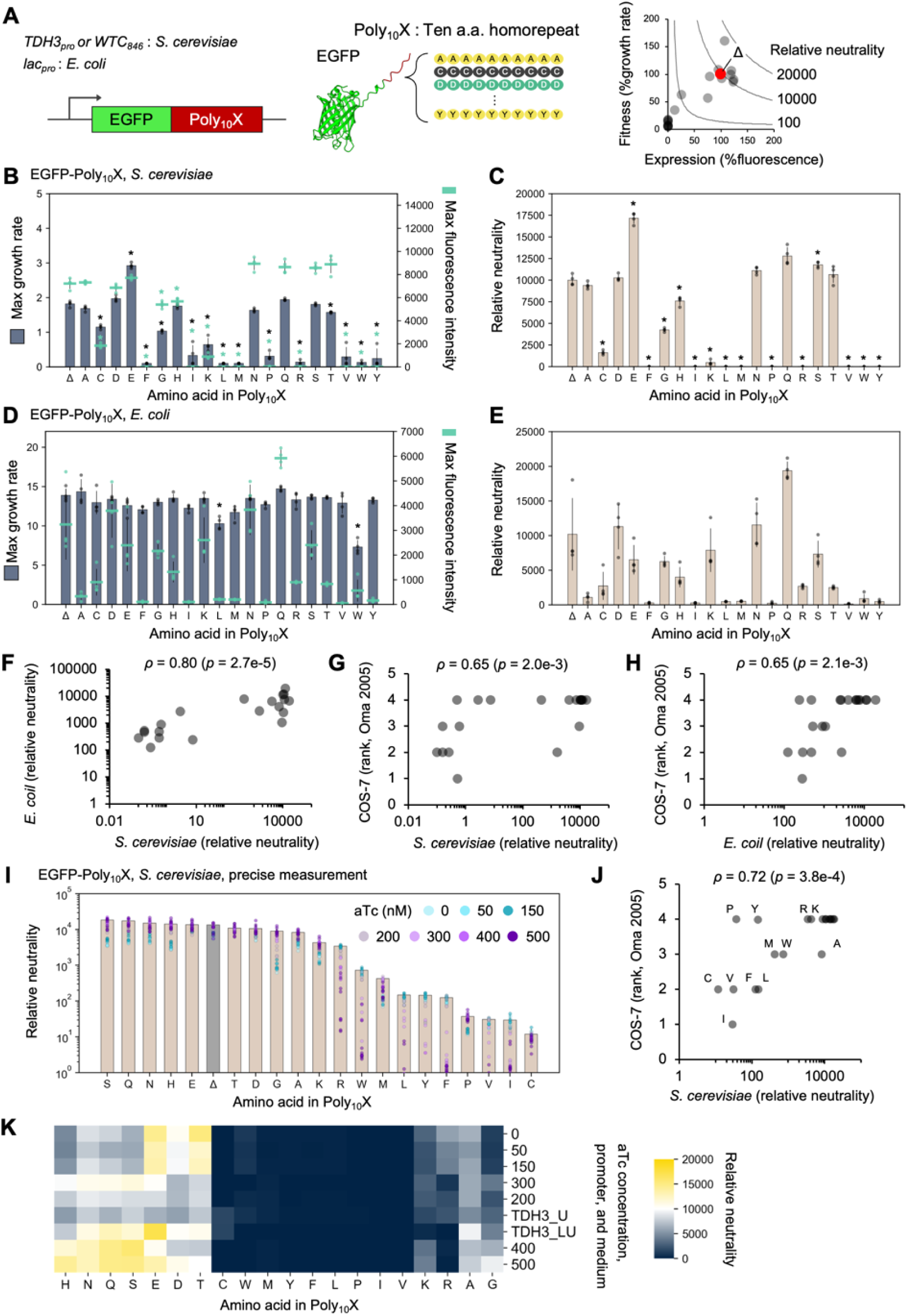
The harmful effects of each Poly_10_X are generally conserved across species. **A)** Experimental setup. Poly_10_X sequences were fused to the C-terminus of EGFP and expressed in *S. cerevisiae* under the *TDH3* or *WTC_846_* promoter, or in *E. coli* under the *lac* promoter. Cellular fitness (growth rate) and expression level (fluorescence intensity) of cells overexpressing each construct were measured. Relative neutrality was calculated by comparison with the corresponding control EGFP construct lacking Poly_10_X (Δ). **B, C)** Maximum growth rate and maximum fluorescence intensity of *S. cerevisiae* cells overexpressing EGFP–Poly_10_X in SC–LU medium at 30 °C, and the calculated relative neutrality (**C**). Growth and fluorescence curves are shown in Figure S2B. **D, E)** Maximum growth rate and maximum fluorescence intensity of *E. coli* expressing EGFP–Poly_10_X in LB + ampicillin medium supplemented with 1 mM IPTG at 37 °C, and the calculated relative neutrality (**E**). Growth and fluorescence curves are shown in Figure S3B. **F–H)** Cross-species comparison of Poly_10_X harmful effects. Average relative neutrality values obtained in *S. cerevisiae* (**C**) and *E. coli* (**E**) were compared with each other and with previously reported Poly_10_X cytotoxicity ranks in COS-7 cells ^26^. Cytotoxicity ranks were calculated as described in *Materials and Methods*. **I)** Precise determination of relative neutrality in *S. cerevisiae*. EGFP–Poly_10_X expression was titrated across seven concentrations of aTc, and the maximum neutrality point was used for final normalization to the EGFP control (Δ). Growth and fluorescence curves are shown in Figure S4B and C. **J)** Correlation between the maximum relative neutrality of Poly_10_X in *S. cerevisiae* (**I**) and the Poly_10_X cytotoxicity ranking in COS-7 cells ^26^. Single-letter codes indicate the amino acid repeated in PolyX. **K)** Hierarchical clustering analysis of relative neutrality values across varying expression levels. On the y-axis, TDH3_U and TDH3_LU indicate experiments using *TDH3_pro_* under SC-U (Figure S1) and SC-LU (**C** and Figure S2) conditions, respectively. The numbers indicate the concentration of aTc in experiments using *WTC_846_* (**I** and Figure S4). Each dataset was normalized to the neutrality of the EGFP control. Euclidean distance and average linkage were used. The heatmap represents relative neutrality levels. In **B**–**E** and **I**, bars, dots, and error bars represent the mean, individual values, and standard deviation from at least three biological replicates. In **B**, **C**, and **D**, asterisks indicate significant differences in maximum growth rate, maximum fluorescence intensity, and relative neutrality, respectively, compared with the control (Δ) (*p* < 0.05, Student’s *t*-test with Bonferroni correction). In **F**–**H** and **J,** Spearman’s rank correlation coefficient (ρ) and its *p*-value are shown.

As a result, the growth rate and fluorescence intensity varied greatly depending on the type of expressed EGFP-Poly_10_X (Figure 1B). Under leucine-depleted conditions, strong growth inhibition was observed for about half of the Poly_10_X variants, and yeast cells showed almost no growth. The harmfulness of each Poly_10_X was quantified as relative neutrality (Figure 1C), revealing that the degree of harmfulness differed substantially among amino acids. The addition of some Poly_10_X sequences markedly reduced the neutrality of EGFP (increasing harmful effects), whereas the addition of other Poly_10_X sequences resulted in neutrality comparable to, or even higher than, that of EGFP.

Next, to examine whether the neutrality of Poly_10_X is conserved across species, we performed equivalent experiments in *Escherichia coli*. Since the gTOW method cannot be applied to *E. coli*, we constructed multicopy plasmids expressing EGFP-Poly_10_X under the control of the *lac* promoter (*lac_pro_*) and induced their expression with isopropyl β-D-1-thiogalactopyranoside (IPTG) (Figure S3). The resulting data were also converted into relative neutrality values (Figures 1D and 1E). In *E. coli*, although the expression levels of EGFP-Poly_10_X varied considerably among the different Poly_10_X variants, the effects on growth were limited (Figure 1D). This is likely due to the weaker selection pressure on plasmid maintenance compared with the yeast gTOW system—plasmids expressing highly harmful Poly_10_X variants are thought to be lost from the population during cultivation (Figure S3E). Previously, the cytotoxicity of PolyX (∼30 residues) in mammalian cells was evaluated by Oma *et al.* ^26^. They assessed PolyX cytotoxicity in African green monkey kidney–derived COS-7 cells by strongly expressing YFP-PolyX under the CMV promoter and measuring cell viability. Because numeric cell viability data were not provided, we relied on reported statistical significance and converted it into four ranks (1–4 corresponding to *p* < 0.001, *p* < 0.01, *p* < 0.05, and *p* > 0.05, respectively). We then compared the Poly_10_X neutrality data obtained in this study for yeast and *E. coli* with those reported by Oma *et al.* for mammalian cells. The results showed a generally high correlation among the three systems (Spearman’s correlation *ρ* > 0.6; Figures 1F-H), indicating that Poly_10_X harmfulness exhibits a conserved trend across species.

To further refine the analysis, we evaluated the neutrality of highly harmful Poly_10_X variants by using the *WTC*_846_ promoter, which allows tunable expression ^52^. By gradually modulating expression levels, we assessed the neutrality of each Poly_10_X variant (Figure S4). The neutrality values (Figure 1I) showed a stronger correlation with mammalian cell cytotoxicity (*ρ* = 0.72, Figure 1J), thereby reinforcing the conclusion that Poly_10_X harmful exhibits a conserved trend across species. Figure 1K compares the neutrality of EGFP-Poly_10_X variants in yeast across a range of expression strengths. Some amino acids were consistently harmful across all expression levels, whereas others were more neutral than EGFP (i.e., beneficial) at specific expression ranges.

### Poly_10_X harmful and beneficial effects are associated with amino acid polarity and hydrophobicity

Next, to examine whether the harmful or beneficial effects of Poly_10_X depend on the physicochemical properties of the fused protein, we fused Poly_10_X to the C-terminus of six different fluorescent proteins and evaluated their neutrality in the same manner (Figure 2A, S5–S9). The obtained neutrality values showed similar trends, with correlations above 0.85 between datasets (Figure 2C). Based on this, we analyzed the relationship between amino acid biochemical and physicochemical parameters and neutrality. Correlation analysis with 28 amino acid indices ^2,53^ (Figure S10A) revealed that biosynthetic energy cost showed the highest correlation (*ρ* = –0.76, *p* = 1.8e-23), followed by polar requirement (*ρ* = 0.66, *p* = 2.2e-16) and hydrophobicity (*ρ* = –0.60, *p* = 4.0e-13) (Figure 2D). Because these parameters mutually showed high intercorrelation (Figure S10B), neutrality may be largely explained by these characteristics.

**Figure 2.**
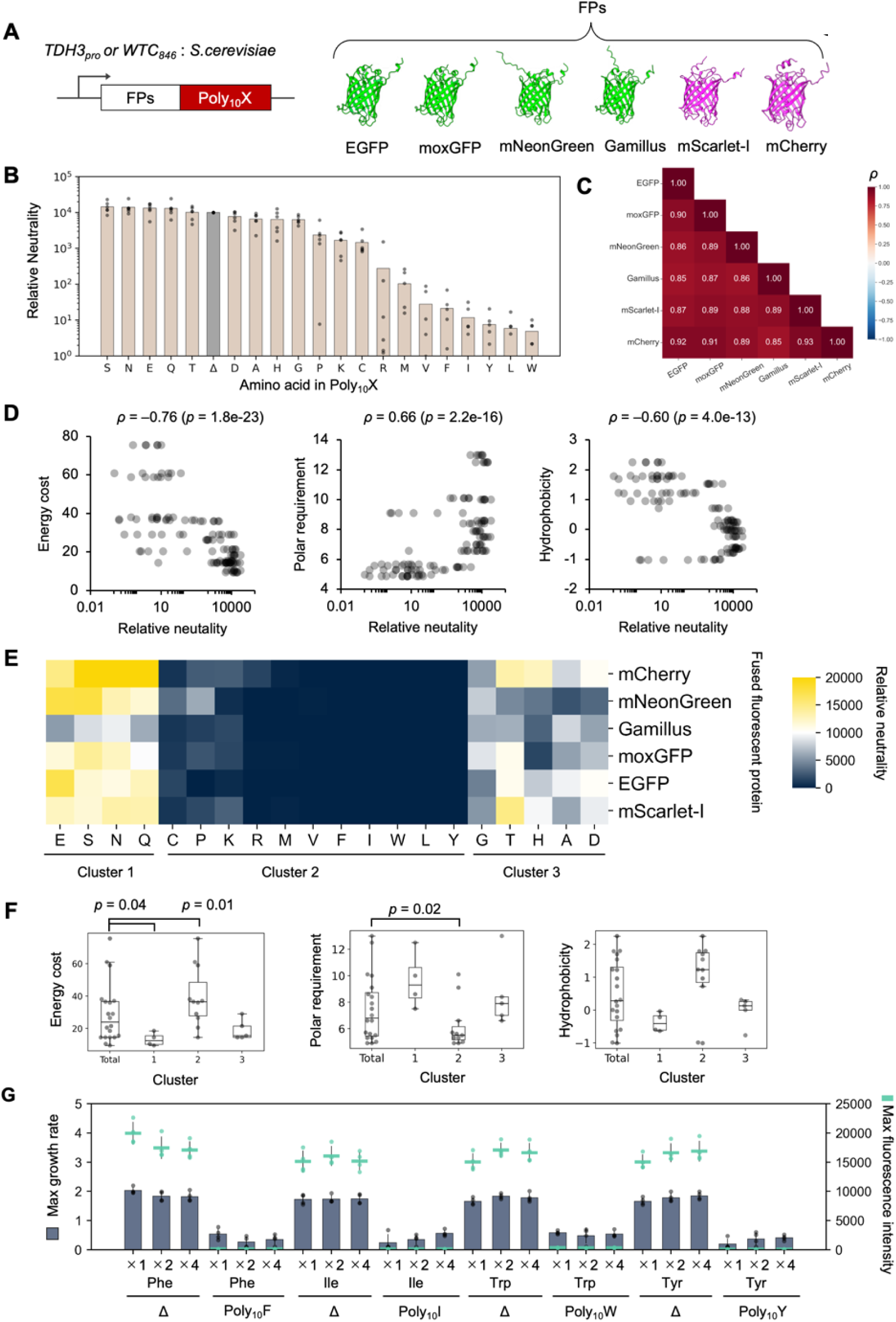
Poly_10_X harmful and beneficial effects are associated with amino acid polarity and hydrophobicity. **A**) Schematic of the constructs used in this study. Each Poly_10_X was attached to the C-terminal of each fluorescent protein (FP), and expressed in *S. cerevisiae* under the control of the *WTC_846_* promoter. EGFP constructs were expressed under the *TDH3* promoter (see Figure 1B, C). **B**) Relative neutrality of Poly_10_X measured using various fluorescent proteins in SC–LU medium at 30 °C. Bars represent the mean relative neutrality for each amino acid, and dots correspond to measurements across different fluorescent proteins. Each dot represents the mean of at least three biological replicates, and the bars indicate the mean relative neutrality obtained across six different fluorescent proteins. Growth and fluorescence curves are shown in Figure S2B, S5B–S9B. **C**) Correlation matrix comparing the trends of Poly_10_X neutrality across different fluorescent proteins. Each value represents the Spearman’s rank correlation coefficient based on relative neutrality values measured for each FPs–Poly_10_X. **D**) Relationships between Poly_10_X neutrality and amino acid physicochemical properties (energy cost, polar requirement and hydrophobicity). Spearman’s rank correlation coefficient (*ρ*) and its *p*-values are shown. **E**) Hierarchical clustering of relative neutrality values obtained from different fluorescent proteins. Clustering was performed using Euclidean distance and average linkage. Single-letter codes indicate the amino acid repeated in Poly_10_X. **F**) Comparison of the three Poly_10_X clusters identified in **E** with amino acid physicochemical properties (energy cost, polar requirement and hydrophobicity). Statistical analyses were performed using the Mann–Whitney *U* test between each cluster and the remaining total samples, followed by Bonferroni correction for multiple comparisons. **G**) Maximum growth rate and maximum fluorescence intensity of *S. cerevisiae* cells overexpressing EGFP without Poly_10_X (Δ) or with C-terminal Poly_10_F, Poly_10_I, Poly_10_W, or Poly_10_Y fusions in SC–LU medium at 30 °C. Indicated amino acid was supplemented to the medium at standard (×1), ×2, or ×4 concentrations. Bars, dots, and error bars represent the mean, individual values, and standard deviation from four biological replicates. Growth and fluorescence curves are shown in Figure S11B.

To further characterize the nature of Poly_10_X, we performed cluster analysis and classified amino acids into three groups (Figure 2E). Cluster 1 included amino acids that increased the relative neutrality above 10,000 for most fluorescent proteins when added to the C-terminus, indicating beneficial effects that reduced protein cytotoxicity. Cluster 2 consisted of amino acids that exhibited harmful effects regardless of the fluorescent protein to which they were attached. Cluster 3 contained amino acids whose effects varied depending on the fused fluorescent protein. Notably, we did not detect any beneficial amino acids in Gamillus, probably because the expression level was insufficient to cause overexpression-dependent growth inhibition, even at maximum induction (Figure S7). Figure 2F shows the representative biochemical and physicochemical properties of amino acids in each cluster. Cluster 1 included E, S, N, and Q, which are low-cost, have high polar requirements, and low hydrophobicity. Cluster 2 mainly contains amino acids with high cost, low polarity, and high hydrophobicity (C, P, M, V, F, I, W, L, Y), but also includes positively charged ones (K, R). Cluster 3 contains G, T, H, and A, which exhibit intermediate cost, polar requirement, and hydrophobicity, while D is somewhat exceptional.

To examine whether the strong harmfulness of the four Poly_10_X variants (F, I, W, and Y) was caused by the high amino acid synthesis cost, we re-evaluated the harmfulness of EGFP (Δ) and four EGFP–Poly_10_X constructs after supplementing the medium with excess corresponding amino acids. As a result, in all cases, the harmfulness was not alleviated (Figure 2G and S11), suggesting that amino acid depletion due to high biosynthetic cost was not the cause of harmfulness. As mentioned above, since the biosynthetic cost correlates well with polarity and hydrophobicity, the apparent correlation with synthesis cost likely reflects a pseudo-correlation arising from these physicochemical properties.

### Structural context modulates the effect of Poly_10_X, while its overall neutrality trend is conserved

So far, we have examined the effects of C-terminally fused Poly_10_X. Next, to investigate the effects of Poly_10_X in different structural contexts, we constructed three types of variants (Figures 3A and S12–14): Poly_10_X was inserted between two fluorescent proteins (EGFP and mCherry); Poly_10_X was inserted into an internal loop of EGFP; and the C-terminally fused Poly_10_X was detached from the fluorescent protein by a P2A sequence ^54^. These designs allowed us to test whether the effects of Poly_10_X depend on the presence of a downstream protein domain, its position within a protein, or its physical coupling to the fluorescent protein. As a result, all constructs showed distinct growth rates, fluorescence levels, and neutrality indices depending on the Poly_10_X sequence (Figures 3B–G and S12–14). It should be noted that in the detached construct, the fluorescence intensity reflects the amount of free EGFP released by the P2A cleavage, but not the amount of the Poly_10_X peptide itself. In contrast, internal Poly_10_X insertions may influence not only protein neutrality but also the fluorescence properties of EGFP, meaning that fluorescence does not necessarily reflect expression levels directly. The neutrality indices among the three structural contexts and the C-terminal EGFP–Poly_10_X construct showed rank correlations above 0.74 (*p* = 2.2e-4) (Figure 3H), indicating that Poly_10_X exhibits a generally conserved trend in neutrality regardless of structural context. Among them, K and P showed markedly reduced harmfulness when placed between two fluorescent proteins compared with when fused to the C-terminus (Figure 3I, upper left). I, V, and W maintained strong harmfulness even in the detached Poly_10_X construct (Figure 3I, lower right), suggesting that the Poly_10_X peptides themselves can exert harmfulness independently. Conversely, the significant harmfulness of C and H (Figure 1C) were lost in the detached Poly_10_X construct (Figure 3G), indicating that these Poly_10_X sequences exhibit harmfulness only when fused to EGFP. No beneficial effect of Poly_10_X was observed when the Poly_10_X were detached (Figure 3G). Therefore, the beneficial effects might arise from mitigating the overexpression-induced harmfulness of fluorescent proteins.

**Figure 3.**
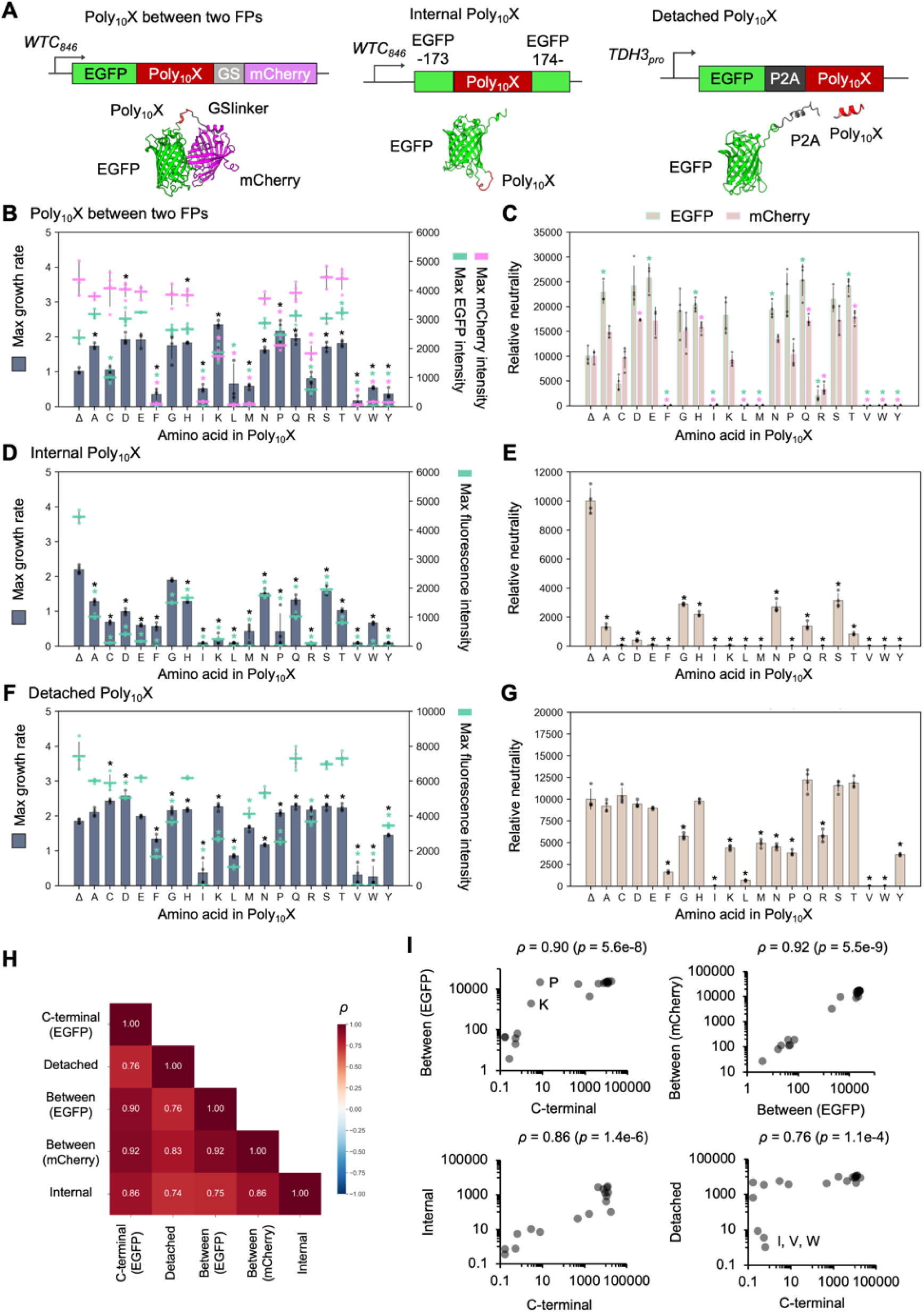
Structural context modulates the effect of Poly_10_X, while its overall neutrality trend is conserved. **A**) Schematic representation of the constructs used to examine the effect of Poly_10_X in different structural contexts. In the “Poly_10_X between two FPs” construct, a Poly_10_X–GS linker sequence was inserted between EGFP and mCherry. In the “internal Poly_10_X” construct, Poly_10_X was inserted between residues 173 and 174 of EGFP. In the “detached Poly_10_X” construct, a P2A self-cleaving sequence was placed between EGFP and Poly_10_X. **B, C**) Maximum growth rate and maximum fluorescence intensity (**B**), and relative neutrality (**C**) of *S. cerevisiae* cells overexpressing the Poly_10_X between two FPs construct in SC–LU medium at 30 °C. Growth and fluorescence curves are shown in Figure S12B. **D**, **E**) Maximum growth rate and maximum fluorescence intensity (**D**), and relative neutrality (**E**) of cells expressing the internal Poly_10_X construct in SC–LU medium at 30 °C. Growth and fluorescence curves are shown in Figure S13B. **F**, **G**) Maximum growth rate and maximum fluorescence intensity (**F**), and relative neutrality (**G**) of cells overexpressing the detached Poly_10_X construct in SC–LU medium at 30 °C. Growth and fluorescence curves are shown in Figure S14B. **H**) Cross-comparison of Poly_10_X neutrality profiles among different structural contexts. Each value represents the Spearman’s rank correlation coefficient calculated from the relative neutrality indices obtained in each construct. **I**) Pairwise comparison of Poly_10_X neutrality trends between different structural contexts. Scatter plots show correlations of average relative neutrality values across constructs. Spearman’s rank correlation coefficient (*ρ*) and its *p*-values are shown. Single-letter codes indicate the amino acid repeated in Poly_10_X. In **B**–**G** bars, dots, and error bars represent the mean, individual values, and standard deviation from at least three biological replicates.

### Poly_10_X induces protein relocalization and aggregate formation

Next, to investigate the effects of Poly_10_X on cellular functions, we observed the intracellular localization of EGFP–Poly_10_X using fluorescence microscopy (Figures 4A and S15–S16). As a result, highly harmful and hydrophobic amino acids—W, I, Y, V, L, and F, except for M—were observed as intracellular aggregates. The appearance of these aggregates varied among amino acids, suggesting that they form distinct types of intracellular assemblies. D and C were enriched in the nucleus, whereas R formed a single bright punctum. Other amino acids showed diffuse cytoplasmic localization similar to the control. These results suggest that highly harmful Poly_10_X sequences exert their harmfulness through aggregation propensity and interactions with cellular structures. Notably, these localization patterns were largely consistent with the subcellular localization of YFP–PolyX observed in COS-7 cells by Oma *et al.* ^27^, indicating that the intracellular behavior of PolyX is conserved among eukaryotic cells.

**Figure 4.**
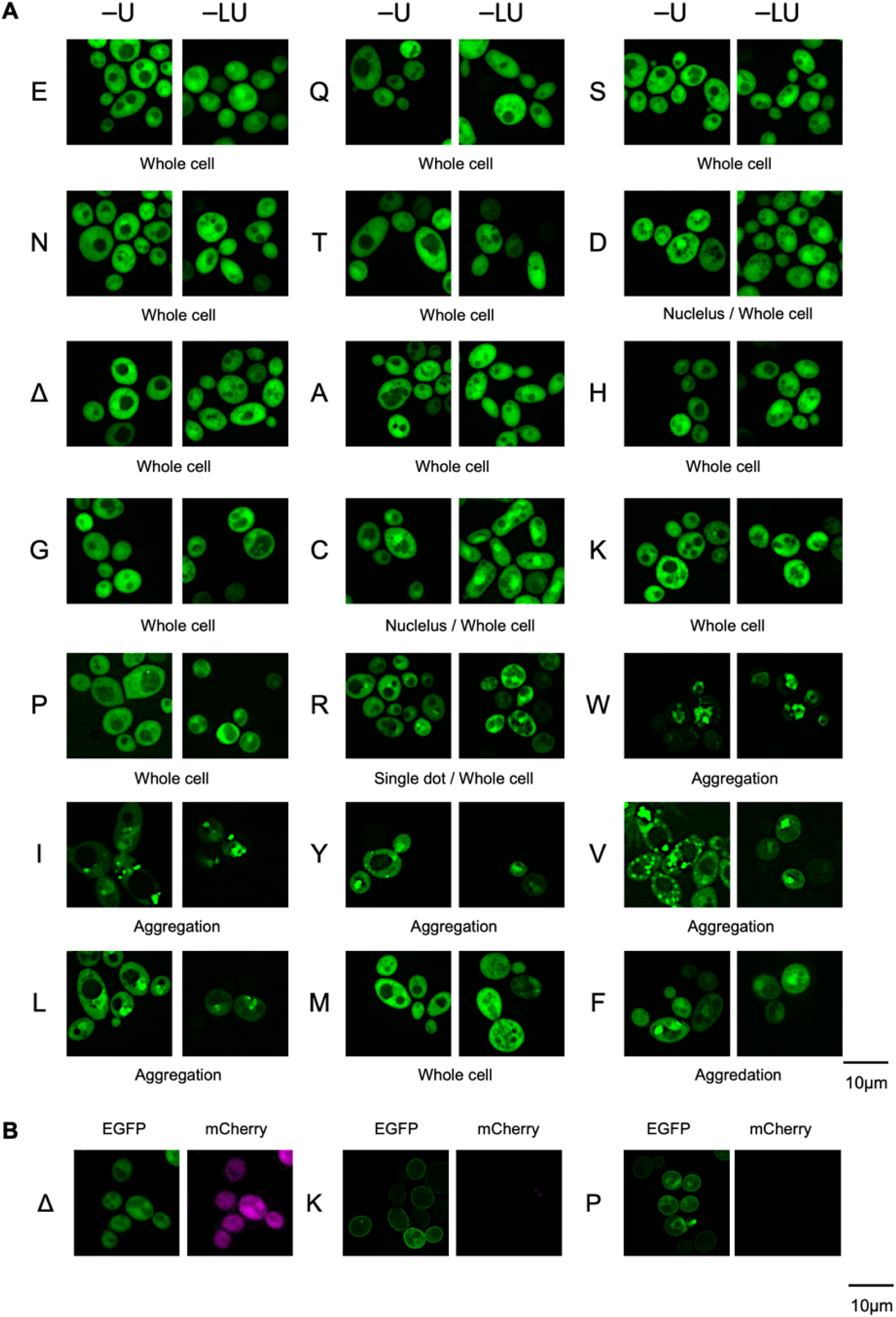
Poly_10_X induces protein relocalization and aggregate formation **A**) Fluorescence microscopy images of *S. cerevisiae* cells expressing EGFP–Poly_10_X under the control of *TDH3_pro_*. Cells were pre-cultured in SC–U medium and then cultured overnight in SC–U or SC–LU medium before imaging shown as –U and –LU. Brightness and contrast were adjusted to allow clear visualization of cell morphology. Indicated subcellular localization and aggregation were categorized by visual inspection. **B**) Fluorescence microscopy images of *S. cerevisiae* cells expressing EGFP–mCherry, EGFP–Poly_10_K–mCherry, and EGFP–Poly_10_P–mCherry under the control of the *WTC_846_* promoter, observed 6 hours after aTc induction. No mCherry fluorescence was detected in cells expressing EGFP–Poly_10_K–mCherry, and EGFP–Poly_10_P–mCherry. Single-letter codes indicate the amino acid repeated in Poly_10_X. Δ represents the protein without a Poly_10_X fusion.

We also examined the localization of constructs in which Poly_10_X was inserted between EGFP and mCherry (Figures S17–S19). For most amino acids, EGFP and mCherry showed identical localization patterns, consistent with those observed for EGFP–Poly_10_X. In contrast, in the Poly_10_K and Poly_10_P constructs, EGFP localized to the cell surface, while mCherry fluorescence was barely detectable (Figure 4B). Because such behavior was not observed in EGFP–Poly_10_K or EGFP–Poly_10_P (Figure 4A), some Poly_10_X can exhibit distinct behaviors depending on the structural context.

### Poly_10_E reduces the harmful effects of protein overexpression through aggregation suppression

As described above, beneficial Poly_10_X sequences are thought to alleviate the harmful effects caused by the overexpression of fluorescent proteins. Overexpressed fluorescent proteins can exert toxicity through multiple mechanisms ^55^. In the case of EGFP, overexpression leads to aggregate formation due to misfolding and imposes a burden on proteostasis ^56^. Since these adverse effects become more pronounced at high temperatures, we evaluated the neutrality of EGFP-Poly_10_X at 38 °C. As a result, the growth of most Poly_10_X-overexpressing strains, including EGFP (Δ), which had been able to grow at 30 °C, was markedly reduced (Figure S20). In contrast, appending Poly_10_E, Poly_10_G, Poly_10_Q, or Poly_10_S maintained growth even under this condition, resulting in a substantial increase in relative neutrality (Figure 5A and Figure S20), with Poly_10_E showing the most pronounced increase in relative neutrality (Figure 5B).

**Figure 5.**
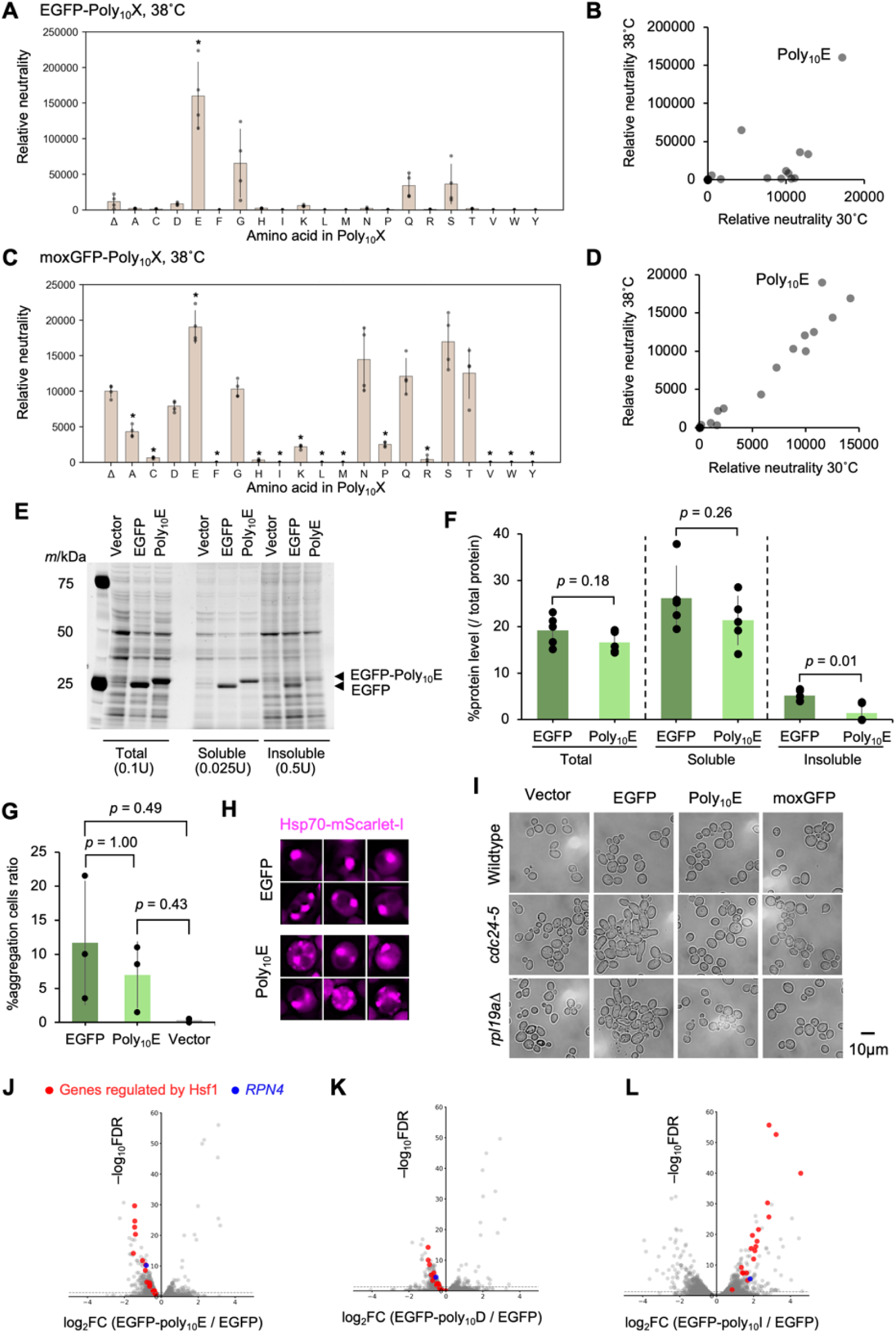
Poly_10_E reduces the harmful effects of protein overexpression through aggregation suppression. **A**) Relative neutrality of EGFP-Poly_10_X under heat stress. Cells were cultured in SC–LU medium at 38 °C. Growth and fluorescence data are shown in Figure S20. **B**) Comparison of the relative neutrality of EGFP–Poly_10_X at 30 °C (from Figure 1C) and 38 °C (from **A**). Average values are used, and the plot of EGFP-Poly_10_E is shown as Poly_10_E. **C**) Relative neutrality of moxGFP-Poly_10_X under heat stress. Cells were cultured in SC–LU medium at 38 °C. Growth and fluorescence data are shown in Figure S21. **D**) Comparison of the relative neutrality of moxGFP–Poly_10_X at 30 °C (from Figure S5D) and 38 °C (from **C**). Average values are used, and the plot of moxGFP-Poly_10_E is shown as Poly_10_E. **E**) Representative image of SDS–PAGE of total, soluble, and insoluble protein fractions from cells overexpressing vector, EGFP, and EGFP–Poly_10_E cultured in SC–LU medium. Proteins were visualized by fluorescent dye staining. The arrowhead indicates the band corresponding to EGFP or EGFP–Poly_10_E. **F**) Quantification of target protein levels (% of total protein) based on the band intensity in **E** and Figure S22. Only monomeric bands were quantified; therefore, polymerized EGFP or EGFP–Poly_10_E may not be accurately represented. Bars, dots, and error bars represent the mean, individual values, and standard deviation from five biological replicates. Statistical comparisons were performed by Welch’s t-test. The raw gel images used for the quantification are shown in Figure S22. **G**) Percentage of cells forming Hsp70 aggregates among those overexpressing vector, EGFP, or EGFP–Poly_10_E. The *p*-values were calculated by Welch’s t-test with Bonferroni correction. For each of three biological replicates, 200 cells per sample were randomly selected, and aggregate formation was visually assessed under blinded conditions. **H**) Fluorescent microscopic images of Hsp70 foci in cells overexpressing EGFP or EGFP–Poly_10_E. Hsp70/Ssa1–mScarlet-I was genomically integrated to monitor Hsp70 aggregate formation. Image brightness and contrast were adjusted to enhance visibility of aggregates. Enlarged images are shown in Figure S23. **I**) Microscopic observation of morphological phenotypes in various yeast strains overexpressing vector, EGFP, EGFP–Poly_10_E, or moxGFP. Wild-type strain BY4741 and mutant strains (*cdc24-5*, *rpl19aΔ*) were cultured in SC–LU medium for 18 h before imaging. Representative images, the morphological quantification method, and the quantitative results for all seven mutants are shown in Figures S24 and S25. **J**–**L**) Transcriptomic analysis of *S. cerevisiae* cells overexpressing EGFP or EGFP–Poly_10_X constructs by RNA-seq. Volcano plots show differential gene expression between cells expressing EGFP and EGFP–Poly_10_E (**J**), EGFP–Poly_10_D (**K**), and EGFP–Poly_10_I (**L**). Genes regulated by the heat-shock transcription factor Hsf1 and the proteasome stress-response gene *RPN4* are highlighted. Cells expressing EGFP–Poly_10_E and EGFP–Poly_10_D were cultured in SC–LU medium, while those expressing EGFP–Poly_10_I were cultured in SC–U together with the EGFP control.

The adverse effects on proteostasis caused by overexpression are limited in the case of moxGFP, whose folding properties have been improved compared to EGFP ^56^. Indeed, all moxGFP-Poly_10_X overexpression strains that were able to grow at 30 °C (Figure S5) also grew at 38 °C, and their relative neutrality did not change significantly (Figure 5C, 5D, and S21). In contrast, although Poly_10_E did not show any beneficial effect in the 30 °C experiments using moxGFP (Figure S6), it became beneficial at 38 °C (Figure 5C and 5D). It is likely that moxGFP, which has stable folding at 30 °C, does not benefit from Poly_10_E under that condition, but as the structure becomes less stable at 38 °C, the beneficial effect of Poly_10_E emerges.

Next, we examined whether the addition of Poly_10_E mitigates the formation of protein aggregates, Hsp70 aggregates, morphological abnormalities in mutant strains, and the induction of the heat shock response that occur upon EGFP overexpression ^56^. The amount of insoluble EGFP-Poly_10_E in overexpressing cells was significantly lower than that of insoluble EGFP (Figure 5E, 5F, and S22). The number of Hsp70 aggregates in EGFP-Poly_10_E–overexpressing cells showed a decreasing trend, though not statistically significant, compared to EGFP-overexpressing cells (Figure 5G), and images showing more dispersed aggregates were frequently observed (Figure 5H and S23). Furthermore, the morphological abnormalities of the *cdc24-5* and *rpl19aΔ* strains induced by EGFP overexpression were not observed in EGFP-Poly_10_E–overexpressing cells (Figure 5I, S24, and S25). Finally, transcriptome analysis by RNA-seq revealed that the heat shock response and the induction of the proteasome-related transcription factor *RPN4*, both triggered by EGFP overexpression, were markedly reduced in EGFP-Poly_10_E (Figure 5J and S26A). In contrast, EGFP-Poly_10_D, which does not show a growth advantage over EGFP (Figure 1C), exhibited only limited suppression of the heat shock response and *RPN4* induction (Figure 5K and S26A). Collectively, all these results support the conclusion that the addition of Poly_10_E to EGFP alleviates the proteostatic burden caused by misfolding during EGFP overexpression.

Notably, when one of the most harmful Poly_10_X variants, Poly_10_I, was appended to EGFP, it induced a strong heat shock response and *RPN4* activation, opposite to the effect observed with Poly_10_E (Figure 5L and S26B). Therefore, Poly_10_I is suggested to either promote misfolding of EGFP more strongly or aggregate on its own, thereby disrupting intracellular proteostasis. These contrasting effects suggest that the physicochemical properties of the appended poly-amino acid strongly influence protein folding and proteostasis.

### The neutrality of Poly_10_X mirrors its evolutionary usage in proteomes

To explore whether the experimentally observed neutrality of Poly_10_X is reflected in natural proteomes, we first analyzed the maximum number of consecutive residues for each of the 20 amino acids (PolyX_max_) and the number of sequences containing ten or more consecutive identical residues (Num-Poly_10_X) in the *S. cerevisiae* S288C reference ORFs. We found that both PolyX_max_ and Num-Poly_10_X showed clear biases (Figure 6A). About half of the amino acids never appeared as stretches longer than ten residues. To test whether the observed PolyX_max_ values are maintained by natural selection, we compared them with those obtained from a simulation using randomized ORF sequences (Shuffled S288C ORFs). This analysis revealed that D, Q, N, S, E, R, H, P, K, A, and V displayed significantly longer consecutive runs than expected (*q* < 0.001, Monte Carlo method with FDR correction). The other amino acids fell within the expected range, and no clear bias toward increasing or decreasing repeat length was detected. Next, using ORFs from 1,392 *S. cerevisiae* isolates (pan *Sc* ORFs) ^57^, we calculated PolyX_max_ and Num-Poly_10_X with higher resolution. In the *S. cerevisiae* S288C reference strain, the amino acids A, V, T, G, L, F, I, Y, and M, which showed zero Num-Poly_10_X values, appeared as Poly_10_X sequences at very low frequencies (≤ 1); Poly_10_X of C and W were never observed (Figure 6A).

**Figure 6.**
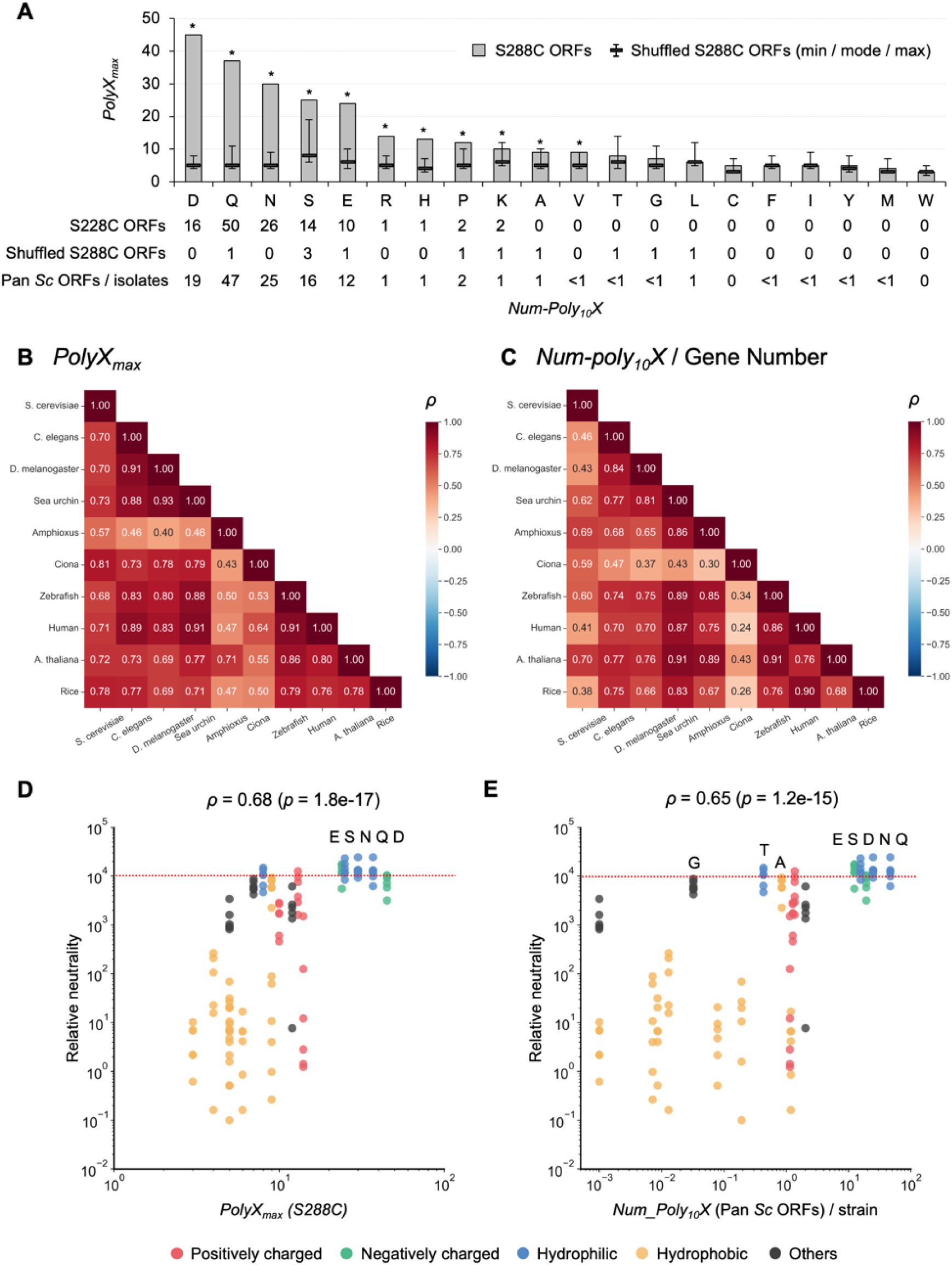
The neutrality of Poly_10_X mirrors its evolutionary usage in proteomes. **A** Analysis of PolyX occurrence patterns (PolyX_max_ and Num–Poly_10_X) in *S. cerevisiae*. The bar graph shows the maximum number of consecutive identical residues (PolyX_max_) for each amino acid in S288C ORFs. Shuffled S288C ORFs represent the minimum, mode, and maximum values obtained from 10,000 random simulations. Asterisks indicate amino acids that exhibit significantly longer homorepeat lengths than expected from simulation (*q* < 0.001, Monte Carlo test with FDR correction). Numbers below the graph indicate the counts of proteins containing ≥10 consecutive identical residues (Num–Poly_10_X) for S288C ORFs, shuffled S288C ORFs, and pan *Sc* ORFs/isolates (from top to bottom). Values for pan-*Sc* ORFs/isolates are rounded to one decimal place. **B** Spearman’s rank correlation coefficient (*ρ*) of PolyX_max_ values among different species. The underlying numerical data used for the calculation are provided in Figure S27. **C** Spearman’s rank correlation coefficients (*ρ*) of Num–Poly_10_X/Gene_number across species. The underlying numerical data used for the calculation are provided in Figure S27. **D** Relationship between PolyX_max_ in S288C ORFs and experimentally measured relative neutrality. Colors indicate the chemical properties of each amino acid. **E** Relationship between Num–Poly_10_X in Pan *Sc* ORFs/isolates and experimentally measured relative neutrality. For cysteine (C) and tryptophan (W), values of 0 were plotted as 0.001 for visualization purposes. In **D** and **E**, Spearman’s rank correlation coefficient (*ρ*) and its *p*-value are shown. Colors indicate the physicochemical properties of each amino acid.

To examine whether the PolyX occurrence patterns observed in *S. cerevisiae* are shared across other organisms, we analyzed nine species (*C. elegans*, *D. melanogaster*, sea urchin, amphioxus, *Ciona*, zebrafish, human, *A. thaliana*, and rice) for PolyX_max_ and Num-Poly_10_X normalized by gene number, and compared their frequencies among species. A comparison of PolyX_max_ between *S. cerevisiae* and the other species showed a high correlation coefficient of 0.57 or greater, suggesting that the overall occurrence pattern of PolyX_max_ is largely conserved across species (Figure 6B). In contrast, a similar comparison using Num-Poly_10_X yielded a lower correlation of 0.38 or higher (Figure 6C). Whether this lower correlation reflects species-specific characteristics or results from annotation errors in ORF datasets remains to be determined.

Finally, we compared the occurrence patterns of PolyX observed in ORFs with the neutrality of Poly_10_X determined in our experiments, in which each Poly_10_X was fused to the C-terminus of a fluorescent protein (Figure 2). As shown in Figure 6D and 6E, both PolyX_max_ and Num-Poly_10_X in the pan-ORF dataset showed strong correlations with neutrality. Five amino acids—E, S, N, Q, and D—were clearly separated from the others, exhibiting longer PolyX_max_ values and higher occurrence numbers in Poly_10_X. These amino acids also displayed beneficial effects when fused to the C-terminus of certain fluorescent proteins (neutrality >10,000). In contrast, hydrophobic amino acids with low occurrence frequencies generally showed low neutrality and high harmfulness. Exceptions were A, G, and T, which exhibited high neutrality despite having an average Num-Poly_10_X of less than one in the pan *Sc* ORF dataset. The probability that these amino acids (A, G, and T) would randomly appear as ten consecutive residues is low (Figure 6A). Therefore, their behavior may represent that of amino acids that are neutral and lack inherent beneficial effects. Taken together, these findings indicate that Poly_10_X neutrality, as determined experimentally, closely explains the occurrence patterns of amino acid homorepeats in natural proteomes.

## Discussion

In this study, we systematically evaluated the neutrality (harmful and beneficial properties) of simple amino acid sequences in which a single residue is repeated ten times (Poly_10_X). We found that neutrality varied markedly among amino acids, with hydrophobic residues exhibiting pronounced harmfulness (Figures 1 and 2). In contrast, several hydrophilic or negatively charged amino acids (e.g., E, S, N, Q) showed beneficial effects on cellular tolerance to protein overexpression, which was particularly intriguing. Notably, the harmfulness of Poly_10_I, Poly_10_V, and Poly_10_W persisted even when detached from EGFP (Figure 3), indicating that these repeats possess intrinsic harmfulness. Conversely, the beneficial effects of Poly_10_E, Poly_10_Q, Poly_10_N, and Poly_10_S disappeared upon detachment from EGFP (Figure 3), indicating that these repeats reduce cellular burden by mitigating the toxicity of the host protein. Mechanistically, the harmful effect of Poly_10_I appears to stem from its intrinsic aggregation propensity, whereas the beneficial effect of Poly_10_E is likely due to its ability to prevent aggregation (Figures 4 and 5).

Furthermore, when we compared the occurrence frequencies of PolyX across the proteomes of diverse organisms, we found that their cross-species occurrence profiles were remarkably similar across species and showed strong correlation with the experimentally determined neutrality values (Figure 6). Notably, hydrophobic residues, which rarely form homorepeats in natural proteins, exhibited high harmfulness in our assays, suggesting that such sequences are likely eliminated immediately upon emergence during evolution. In contrast, amino acids such as E, N, Q, and S—whose homorepeats are widely observed in natural proteins—showed low harmfulness and, in some cases, even beneficial effects, implying that their appearance may confer an advantage to cells from the outset. PolyQ, for example, can contribute to normal protein function by modulating protein–protein interactions when maintained at moderate lengths, but become pathogenic upon excessive expansion ^39^. Thus, the evolutionarily permissible range of PolyQ length is likely determined by a balance between harm and benefit.

Taken together, the occurrence frequencies of amino acid sequences in natural proteomes are well explained by experimentally determined neutrality, reflecting a balance between harmful and beneficial effects. Our assays were performed by placing simple homorepeat sequences at the C-terminus or within internal loops of fluorescent proteins and expressing them in the cytosol. Because homorepeats found in natural proteins predominantly occur in intrinsically disordered regions (Figure S29), our experimental design provides an appropriate framework for evaluating their harmful and beneficial effects. In contrast, whether similar principles apply to more complex sequences buried within membranes or folded protein cores remains an open question. For structured or interaction-dependent motifs, intrinsic harmfulness might be masked—or even neutralized—by specific binding partners. This idea raises the possibility that “eliminating harmfulness” itself acts as an evolutionary pressure driving the formation of higher-order structures or interaction modules. Interestingly, hydrophobic homorepeats—among the most harmful sequences in our assays—are also utilized as secretion signals ^58^. It is tempting to speculate that what was originally a “dangerous” sequence requiring removal from the cytosol may have been evolutionarily co-opted into the secretory pathway.

The design of artificial proteins is rapidly advancing. However, it remains difficult to determine whether sequences absent in nature are missing because they offer no benefit or because they are eliminated due to harmfulness. The parameter we identified in this study—neutrality, which integrates both harmful and beneficial effects—may represent a fundamental evolutionary design principle that shapes the compatible amino acid sequence space in living organisms. A deeper understanding of this parameter will aid in rational protein design and may also shed light on the molecular mechanisms underlying neurodegenerative diseases and aging, in which mutations elevate protein harmfulness.

## Materials and Methods

### Strains, plasmids, growth conditions

*S. cerevisiae* and *E. coli* strains and plasmids used in this study are listed in the Key Resources Table. Synthetic complete (SC) medium lacking uracil (U) or leucine (L) was used for yeast cultures. LB medium was used for *E. coli* cultures.

### Genetic tug-of-war (gTOW) method

The gTOW method ^50,51^ enables artificial elevation of plasmid copy number for a target protein by exploiting the combination of a multi-copy plasmid replication origin (2μ ORI) and the auxotrophic selection marker *leu2-89*. In SC–U medium, where no copy-number–increasing selection pressure is present, the plasmid remains at a low copy number (approximately 30 copies for an empty vector). In contrast, in SC–LU medium, strong selection pressure increases the plasmid copy number to approximately 150 copies. As plasmid copy number increases, the expression level of the target protein also rises. If expression becomes high enough to inhibit cell growth before the plasmid reaches ∼150 copies, this expression level is defined as the protein’s maximum tolerable expression level. Thus, gTOW allows quantitative determination of how much of a given protein a yeast cell can tolerate before growth inhibition occurs. By combining the measured expression level with the degree of growth inhibition, protein harmfulness can be represented as a one-dimensional parameter, relative neutrality, enabling direct comparison of harmfulness across different proteins ^55^. In this study, modified fluorescent proteins were expressed under either the strong *TDH3* promoter or the *WTC_846_* (*P_7.tet1_*) promoter, a *TDH3*-derived variant whose expression can be induced by anhydrotetracycline (aTc) ^52^.

### Measurement of growth rate, fluorescence intensity, and calculation of the relative neutrality in yeast

Cells of the BY4741 or BYW2 strain carrying the expression plasmids were pre-cultured in SC–U and then transferred to either SC–LU (for constructs expressed under *TDH3_pro_* in BY4741) or SC–LU supplemented with aTc (for constructs expressed under the *WTC_846_* promoter in BYW2). Cells were cultured under these conditions while OD_595_ and fluorescence signals were monitored every 10 or 30 minutes using an Infinite F Nano+ microplate reader. Fluorescence of EGFP, moxGFP, mNeonGreen, and Gamillus was measured using an excitation/emission filter set of 485/535 nm, whereas fluorescence of mScarlet-I and mCherry was measured using a 535/590 nm filter set. From these time-course measurements, the maximum growth rate (MGR) and maximum fluorescence intensity (MFI) were determined using a custom Python script. The relative neutrality index was calculated by multiplying the percentage of MGR relative to the control vector (Δ) by the percentage of MFI relative to the control fluorescent protein as follows: Relative neutrality = %(MGR_Poly_10_X / MGR_Δ) × %(MFI_Poly_10_X / MFI_Δ).

### Measurement of growth rate, fluorescence intensity, and calculation of the relative neutrality in *E. coli*

Cells of the BW25113 strain carrying the expression plasmids were pre-cultured in LB + ampicillin medium and then transferred to fresh LB + ampicillin medium. Cells were cultured under these conditions while OD_595_ and fluorescence (485/535 nm) signals were monitored every 5 minutes using an Infinite F Nano+ microplate reader. From these time-course measurements, the maximum growth rate (MGR) and maximum fluorescence intensity (MFI) were determined using a custom Python script. The relative neutrality index was calculated by multiplying the percentage of MGR relative to the control vector (Δ) by the percentage of MFI relative to the control fluorescent protein as follows: Relative neutrality = %(MGR_Poly_10_X / MGR_Δ) × %(MFI_Poly_10_X / MFI_Δ).

### Clustering analysis

Clustering analysis was performed using a custom Python script (pandas, seaborn, and matplotlib) after organizing the relative neutrality data in an Excel file. Hierarchical clustering and heatmap visualization were conducted using the clustermap function in seaborn. Euclidean distance (metric = “euclidean”) was used as the distance metric, and average linkage (UPGMA; method = “average”) was applied for clustering. The resulting clustering patterns were visualized as row- and column-wise heatmaps, with dendrograms shown or hidden depending on the purpose of the analysis. A custom continuous colormap defined by three reference points was used, and the display range was fixed from 0 to 20,000. This scaling was chosen because relative neutrality values were normalized such that the control (Δ) corresponded to 10,000, representing the non-toxic state; accordingly, the color scale was centered at 10,000.

### Amino acid parameters

For the correlation analysis in Figure S10, a total of 28 amino acid parameters were used, consisting of 18 nonredundant indices from the AAindex database (https://www.genome.jp/aaindex/; see https://doi.org/10.1002/pro.5239) and 10 additional parameters: hydrophobicity ^59^, hydropathy index ^60^, isoelectric point, side-chain molecular weight, amino acid usage (%) (this study), metabolic cost ^61^, biosynthetic cost (ATPs) ^62^, biosynthetic steps (enzymes) ^62^, and total energy costs of amino acids and nucleotide precursors under fermentative or respiratory conditions ^63^.

### Microscopic observation

Yeast cells overexpressing the target protein were cultured in either SC–U or SC–LU medium, and imaged using a DMI6000B microscope (Leica Microsystems), and images were processed using the Leica Application Suite X software. GFP and RFP fluorescence signals were acquired using a GFP filter cube and RFP filter cube, respectively.

For the morphology analysis in Figure S24 and S25, microscopic images were processed using Cellpose ^64^ for cell segmentation. The segmented images were subsequently analyzed using the *MeasureObjectSizeShape* module in CellProfiler ^65^ to quantify the major and minor axes of individual cells. Elongation ratio values were calculated for each cell using the major and minor axis lengths obtained from the *MeasureObjectSizeShape* module, either with a custom Python script or in Excel, according to the following formula: Elongation Ratio = MajorAxisLength / MinorAxisLength. Cells with an elongation ratio ≥ 1.5 were defined as morphologically abnormal.

### Protein analysis

BY4741 strains expressing the target proteins were cultured overnight in 6 mL or 25 mL of SC–LU medium. Total protein was extracted from cells in the logarithmic growth phase (OD_660_ = 0.9–1.0) using 0.2 mol/L NaOH, followed by solubilization in NuPAGE sample buffer (Thermo Fisher Scientific). For each analysis, total protein was extracted from a cell mass corresponding to 1 OD unit at OD_660_.

For the cell lysate analysis, cells were washed with PBST [10 mM phosphate-buffered saline (pH 7.4), 0.001% Tween 20] supplemented with Halt protease inhibitor cocktail (Thermo Fisher Scientific), then disrupted with glass beads using a bead-beating homogenizer (Micro Smash MS-100, TOMY) at 5000 rpm for 30 s, repeated five times. Cell lysates were centrifuged at 15,000 rpm for 10 min to separate soluble and insoluble fractions. The insoluble fraction was washed with 1 mL PBST and centrifuged again at 15,000 rpm for 10 min.

Extracted proteins were fluorescently labeled using EZLabel Fluoro NEO (ATTO) according to the manufacturer’s instructions and separated by SDS–PAGE on 4–12% gradient gels. Fluorescent signals were detected using the SYBR Green fluorescence mode of a LAS-4000 image analyzer (GE Healthcare). Proteins separated by SDS–PAGE were transferred onto a PVDF membrane (Thermo Fisher Scientific). GFP signals were detected using an anti-GFP primary antibody (Roche), an peroxidase-conjugated secondary antibody (Nichirei Bioscience), and chemiluminescent substrate (Thermo Fisher Scientific). Chemiluminescent images were acquired with the chemiluminescence mode of the LAS-4000 image analyzer.

### RNA sequencing analysis

*S cerevisiae* strain BY4741 with the control vector or overexpressing EGFP, EGFP–Poly_10_D, or EGFP–Poly_10_E was cultured in SC–LU medium, and strain BY4743 with the vector or overexpressing EGFP, or EGFP–Poly_10_I was cultured in SC–U medium. Cells were collected at the logarithmic growth phase (OD_660_ ≈ 1.0). Total RNA was extracted according to a previously described protocol ^66^. The concentration of purified RNA was initially measured using a Qubit fluorometer (Thermo Fisher Scientific), and samples were stored at −80 °C until further use. Before library preparation, RNA concentrations were re-measured using the Quant-iT RiboGreen RNA Assay Kit (Invitrogen) on an ARVO Multimode Microplate Reader (PerkinElmer), and RNA quality was assessed with a 2100 Bioanalyzer (Agilent Technologies). cDNA libraries were prepared using the TruSeq Stranded mRNA Library Prep Kit (Illumina) and sequenced on a NovaSeq X Plus platform (Illumina) with 100-bp paired-end reads. Raw sequencing data were quality-checked using FastP ^67^ and aligned to the reference genome with Hisat2 ^68^ . Aligned data were converted to BAM format with Samtools ^69^ and quantified using StringTie ^70^. Differential gene expression analysis was performed using EdgeR ^71^. Gene Ontology (GO) enrichment analysis was conducted with the clusterProfiler ^72^ and org.Sc.sgd.db packages (DOI: 10.18129/B9.bioc.org.Sc.sgd.db) in R to identify significantly enriched GO terms. All analyses were performed using three biological replicates for each strain. The raw data were deposited into DDBJ (accession number: PRJDB39951).

### PolyX analysis in the *S. cerevisiae* proteome and other species

Proteome-wide PolyX analysis was performed using all verified ORFs registered in the *Saccharomyces* Genome Database (SGD), covering all 20 amino acids. As of October 2024, 6,585 ORFs are annotated in the *S. cerevisiae* (S288C) genome. From this set, 5,911 verified ORFs (Sc ORFs) were retained for analysis after removing those annotated as “Dubious.” A custom Python script first loaded the ORF list and excluded all entries whose Qualifier field was labeled “Dubious.” Regular expressions were then used to identify, for each ORF, the longest homorepeat for each amino acid. The resulting maximum repeat lengths were compiled into a dataframe, and two metrics were calculated as follows: PolyX_max_ – the maximum length of consecutive identical amino acids for each amino acid across all ORFs, and Num_Poly_10_X – the number of ORFs containing a homorepeat of length ≥ 10 for each amino acid. For pan *Sc* ORFs, the same analysis was conducted using FASTA assemblies downloaded from Zenodo (https://zenodo.org/records/3407352), which include 1,392 *S. cerevisiae* genome assemblies ^57^.

Simulated (shuffled) S288C ORFs were generated as follows using a custom Python script. Each sequence of above 5,911 ORFs was randomly shuffled while preserving amino acid composition. This produced randomized sequences with identical amino acid compositions but altered residue order. PolyX_max_ and Num_Poly_10_X values were computed for each shuffled ORF set using the same regular expression pipeline described above. This simulation was repeated 10,000 times, using seed values from 0 to 9,999. For each iteration, PolyX_max_ and Num_Poly_10_X results were stored in a dataframe. For every amino acid, the minimum, mode, mean, maximum, and standard deviation across the 10,000 trials were calculated and summarized.

For cross-species comparisons, proteomes of nine eukaryotic species (*Caenorhabditis elegans, Drosophila melanogaster, Strongylocentrotus purpuratus, Branchiostoma lanceolatum, Ciona intestinalis, Danio rerio, Homo sapiens, Arabidopsis thaliana*, and *Oryza sativa*) were downloaded from Ensembl databases (https://asia.ensembl.org). When multiple isoforms were annotated for a given gene, the isoform with the longest amino acid sequence was selected as the representative sequence for analysis. For each amino acid sequence, PolyX of identical amino acids were detected based on regular-expression matching. Two metrics were calculated for each of the 20 amino acids: (i) PolyXmax, defined as the maximum length of consecutive identical residues observed across all proteins, and (ii) Num–Poly_10_X, defined as the number of proteins containing a homorepeat of length ≥10 residues. For interspecies comparisons, Num–Poly_10_X values were normalized by the total number of genes analyzed for each species. Correlations between PolyX occurrence metrics (PolyXmax and Num–Poly_10_X) and experimentally determined Poly_10_X neutrality values were evaluated using Spearman’s rank correlation coefficients.

## Data availability statement

All source data used for figure generation, and all analysis scripts have been deposited on GitHub (https://github.com/hisaomlab/Murase_PolyX). RNA-seq data have been deposited in a public repository (DDBJ accession number: PRJDB39951). All plasmids used in this study are available from the National BioResource Project (NBRP)-yeast (https://yeast.nig.ac.jp/yeast/).

## Acknowledgements

We thank the members of the Moriya laboratory (Okayama University) for helpful discussions, Yuhei Chadani (Okayama University) for providing the *E. coli* strain, and Christian Landry (Université Laval) for valuable comments and suggestions on the manuscript.

## Supplementary Figures

**Figure S1.**
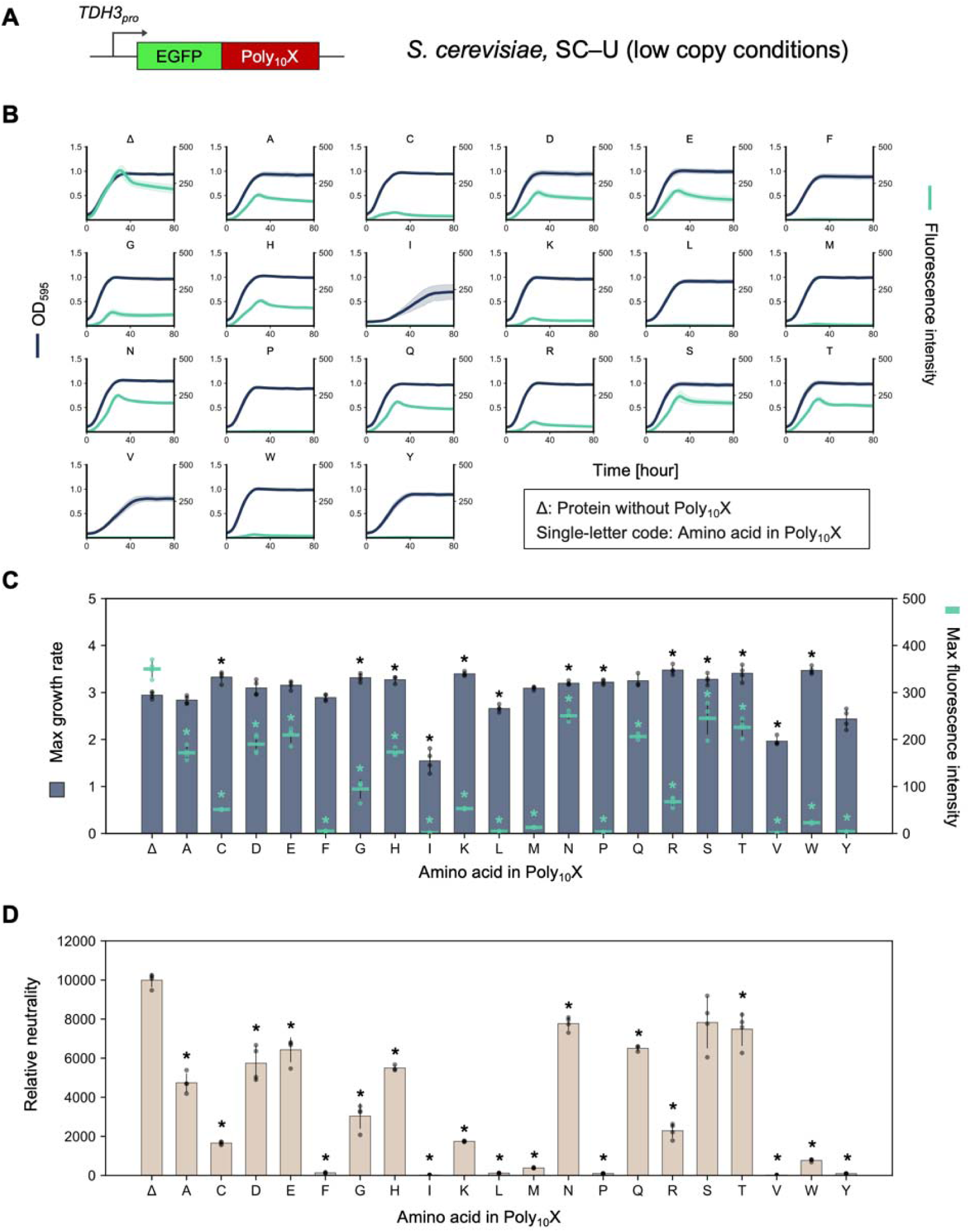
Neutrality of C-terminal Poly10X fusions to EGFP in yeast (low-copy conditions). **A)** Schematic representation of the expression constructs. Poly_10_X was fused to the C-terminus of EGFP and expressed under the *TDH3* promoter in *S. cerevisiae*. **B)** Growth and fluorescence curves of *S. cerevisiae* cells expressing EGFP–Poly_10_X, measured using the gTOW method in SC–U medium at 30 °C. Curves represent the mean values from at least three biological replicates, and shaded regions indicate the standard deviation (SD). **C, D)** Maximum growth rate and maximum fluorescence intensity of *S. cerevisiae* cells low-level overexpressing EGFP–Poly_10_X in SC–U medium at 30 °C, and the calculated relative neutrality (**D**). Bars, dots, and error bars represent the mean, individual data points, and standard deviation from at least three biological replicates. Asterisks indicate significant differences in maximum growth rate, maximum fluorescence intensity, and relative neutrality, respectively, compared with the control (Δ) (*p* < 0.05, Student’s *t*-test with Bonferroni correction).

**Figure S2.**
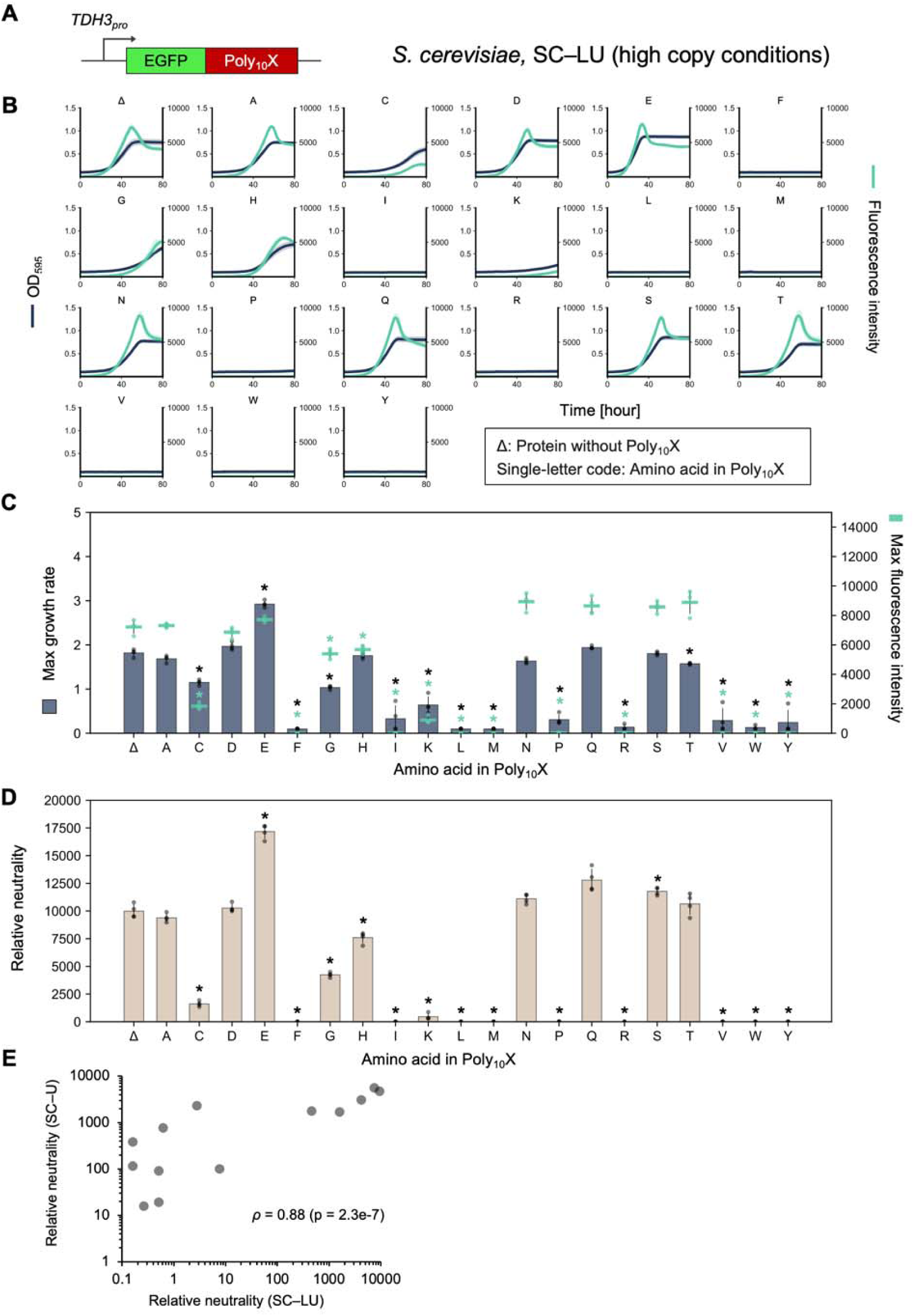
Neutrality of C-terminal Poly_10_X fusions to EGFP in yeast (High-copy conditions). **A)** Schematic representation of the expression constructs. Poly_10_X was fused to the C-terminus of EGFP and expressed under the *TDH3* promoter in *S. cerevisiae*. **B)** Growth and fluorescence curves of *S. cerevisiae* cells expressing EGFP–Poly_10_X, measured using the gTOW method in SC–LU medium at 30 °C. Curves represent the mean values from at least three biological replicates, and shaded regions indicate the standard deviation (SD). **C, D)** Maximum growth rate and maximum fluorescence intensity of *S. cerevisiae* cells overexpressing EGFP–Poly_10_X in SC–LU medium at 30 °C, and the calculated relative neutrality (**D**). Bars, dots, and error bars represent the mean, individual data points, and standard deviation from at least three biological replicates. Asterisks indicate significant differences in maximum growth rate, maximum fluorescence intensity, and relative neutrality, respectively, compared with the control (Δ) (*p* < 0.05, Student’s *t*-test with Bonferroni correction). **E** Comparison of Poly_10_X harmfulness trends under low-level (SC–U) and high-level (SC–LU) overexpression conditions. Spearman’s rank correlation coefficient (*ρ*) is shown.

**Figure S3.**
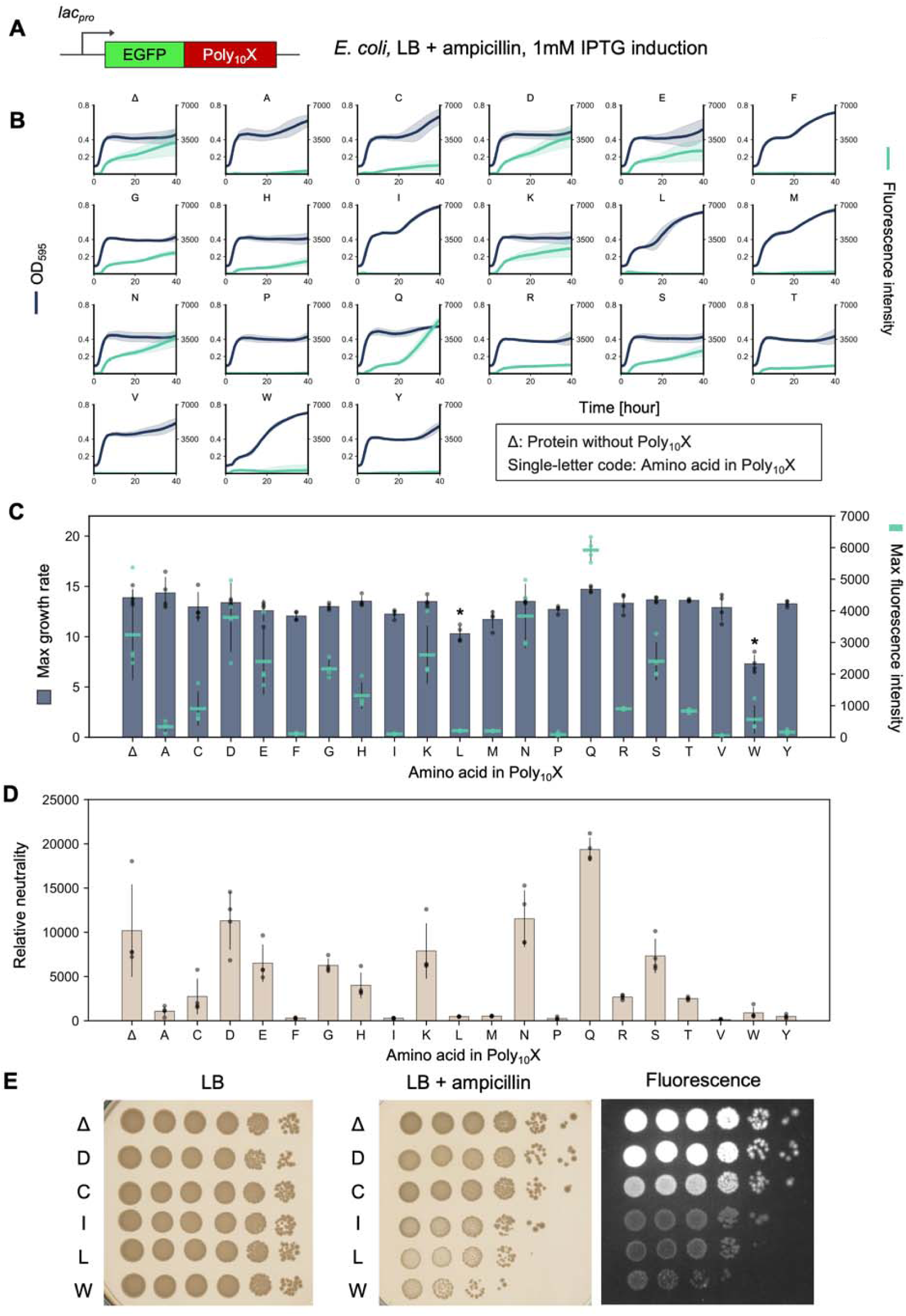
Neutrality of C-terminal Poly_10_X fusions to EGFP in *E. coli*. **A)** Schematic representation of the expression constructs. Poly_10_X was fused to the C-terminus of EGFP and expressed under the *lac* promoter in *E. coli*. **B)** Growth and fluorescence curves of *E. coli* cells expressing EGFP–Poly_10_X, measured in LB + ampicillin medium at 37 °C. Curves represent the mean values from four biological replicates, and shaded regions indicate the standard deviation (SD). **C, D)** Maximum growth rate and maximum fluorescence intensity of *E. coli* cells expressing EGFP–Poly_10_X in LB + ampicillin medium at 37 °C, and the calculated relative neutrality (**D**). Bars, dots, and error bars represent the mean, individual data points, and standard deviation from four biological replicates. Asterisks indicate significant differences in maximum growth rate, maximum fluorescence intensity, and relative neutrality, respectively, compared with the control (Δ) (*p* < 0.05, Student’s *t*-test with Bonferroni correction). **E** Several *E. coli* strains expressing EGFP–Poly_10_X were collected after cultivation in **B**, serially diluted 10-fold, and spotted (5 µl each) onto LB or LB + ampicillin agar plates. Plates were incubated overnight at 37 °C and photographed the following day. These data suggest plasmid loss after cultivation of cells expressing harmful Poly10X variants. Single-letter codes indicate the amino acid repeated in Poly_10_X. Δ represents the protein without a Poly_10_X fusion.

**Figure S4.**
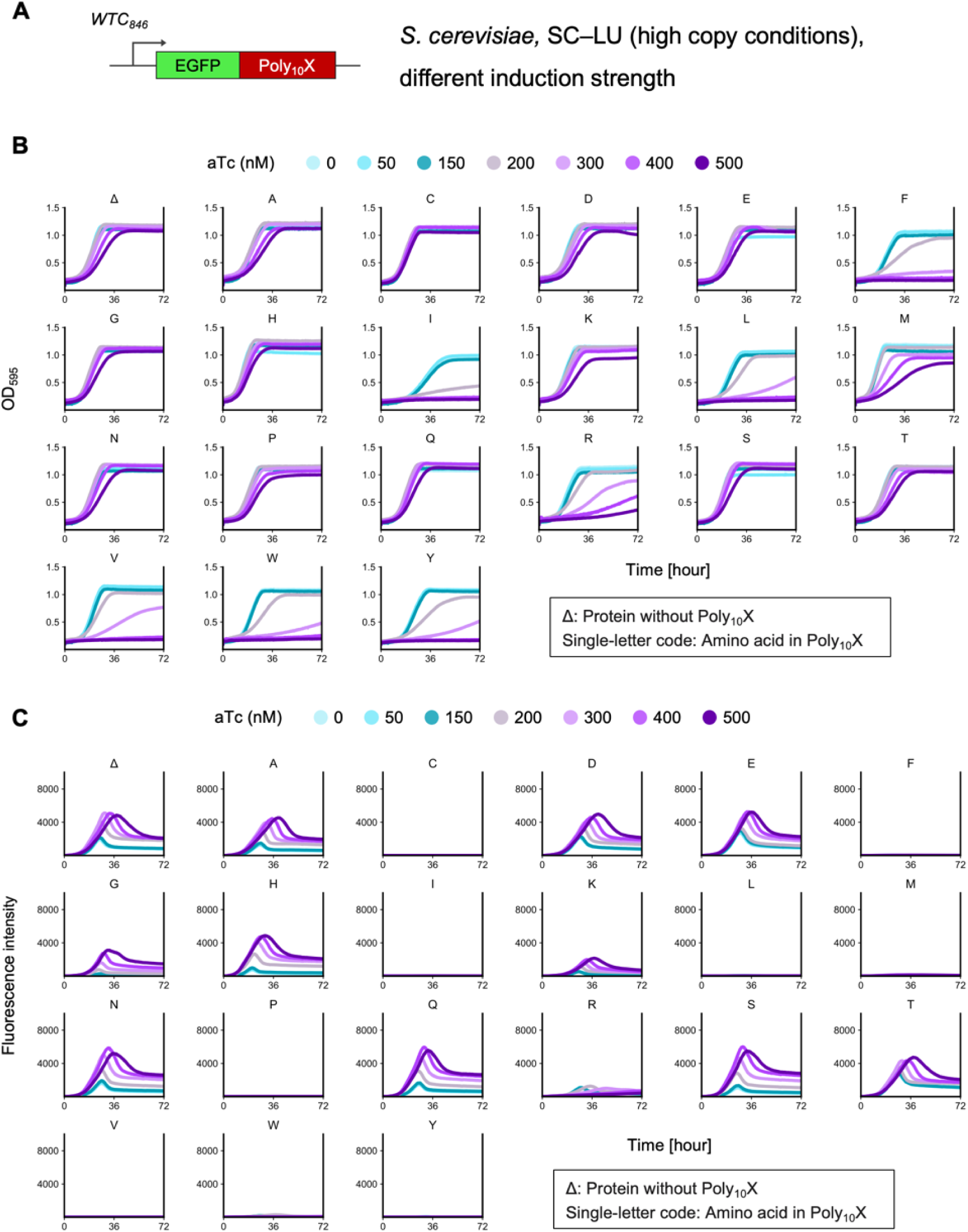
Neutrality of C-terminal Poly_10_X fusions to EGFP under different induction levels in yeast. **A)** Schematic representation of the expression constructs. Poly_10_X was fused to the C-terminus of EGFP and expressed under the control of the *WTC_846_* promoter in *S. cerevisiae*. **B, C)** Stepwise growth (**B**) and fluorescence (**C**) curves of *S. cerevisiae* cells expressing EGFP–Poly_10_X measured using the gTOW method in SC–LU medium at 30 °C. Gradual induction of expression was achieved under the control of the *WTC_846_* promoter by stepwise adjustment of aTc concentration. Curves represent the mean values from four biological replicates.

**Figure S5.**
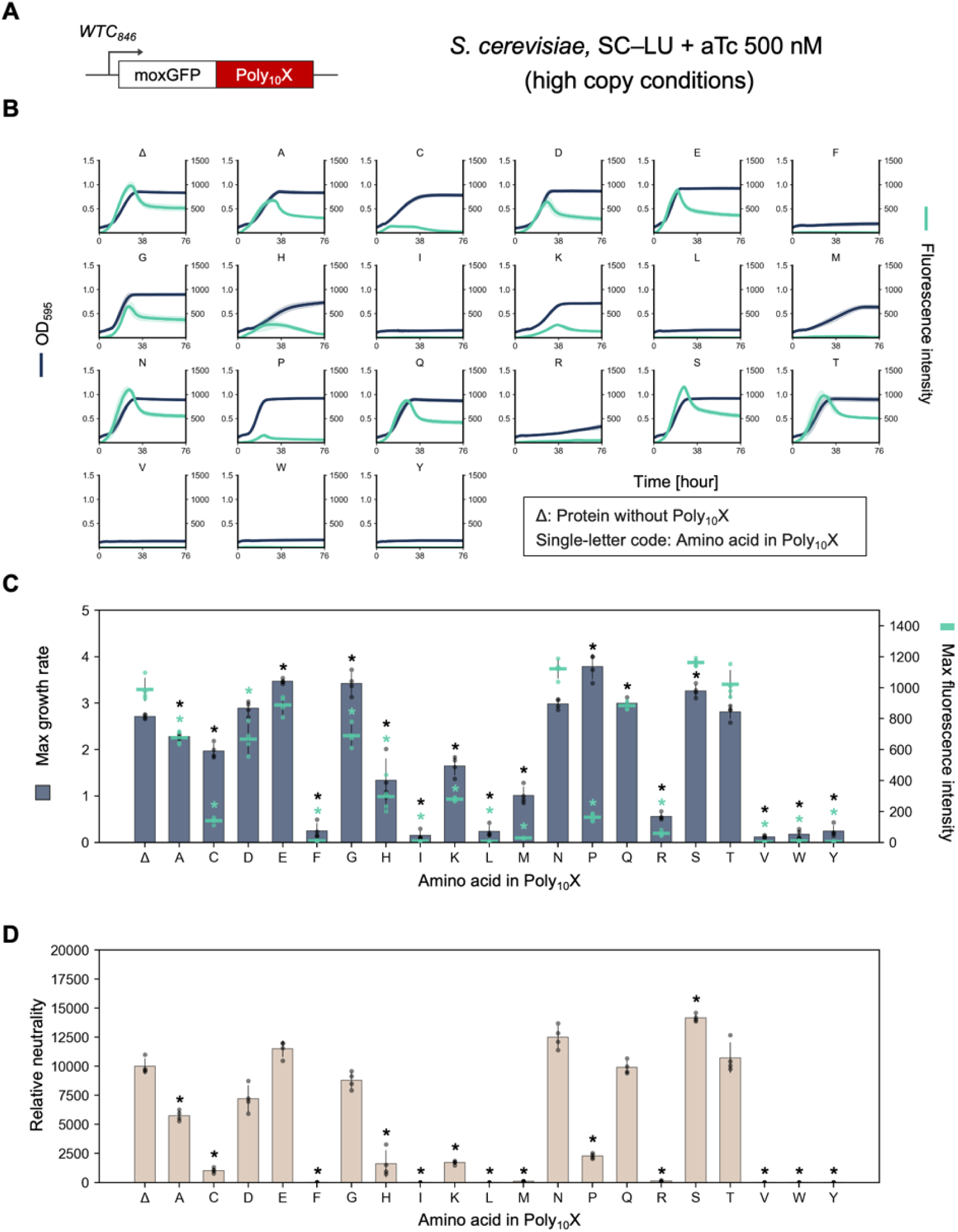
Neutrality of C-terminal Poly_10_X fusions to moxGFP in yeast. **A)** Schematic representation of the expression constructs. Poly_10_X was fused to the C-terminus of moxGFP and expressed under the control of the *WTC_846_* promoter in *S. cerevisiae*. **B)** Growth and fluorescence curves of *S. cerevisiae* cells expressing moxGFP–Poly_10_X, measured using the gTOW method in SC–LU medium at 30 °C. Curves represent the mean values from at least three biological replicates, and shaded regions indicate the standard deviation (SD). **C, D)** Maximum growth rate and maximum fluorescence intensity of *S. cerevisiae* cells overexpressing moxGFP–Poly_10_X in SC–LU medium at 30 °C, and the calculated relative neutrality (**D**). Bars, dots, and error bars represent the mean, individual data points, and standard deviation from at least three biological replicates. Asterisks indicate significant differences in maximum growth rate, maximum fluorescence intensity, and relative neutrality, respectively, compared with the control (Δ) (*p* < 0.05, Student’s *t*-test with Bonferroni correction).

**Figure S6.**
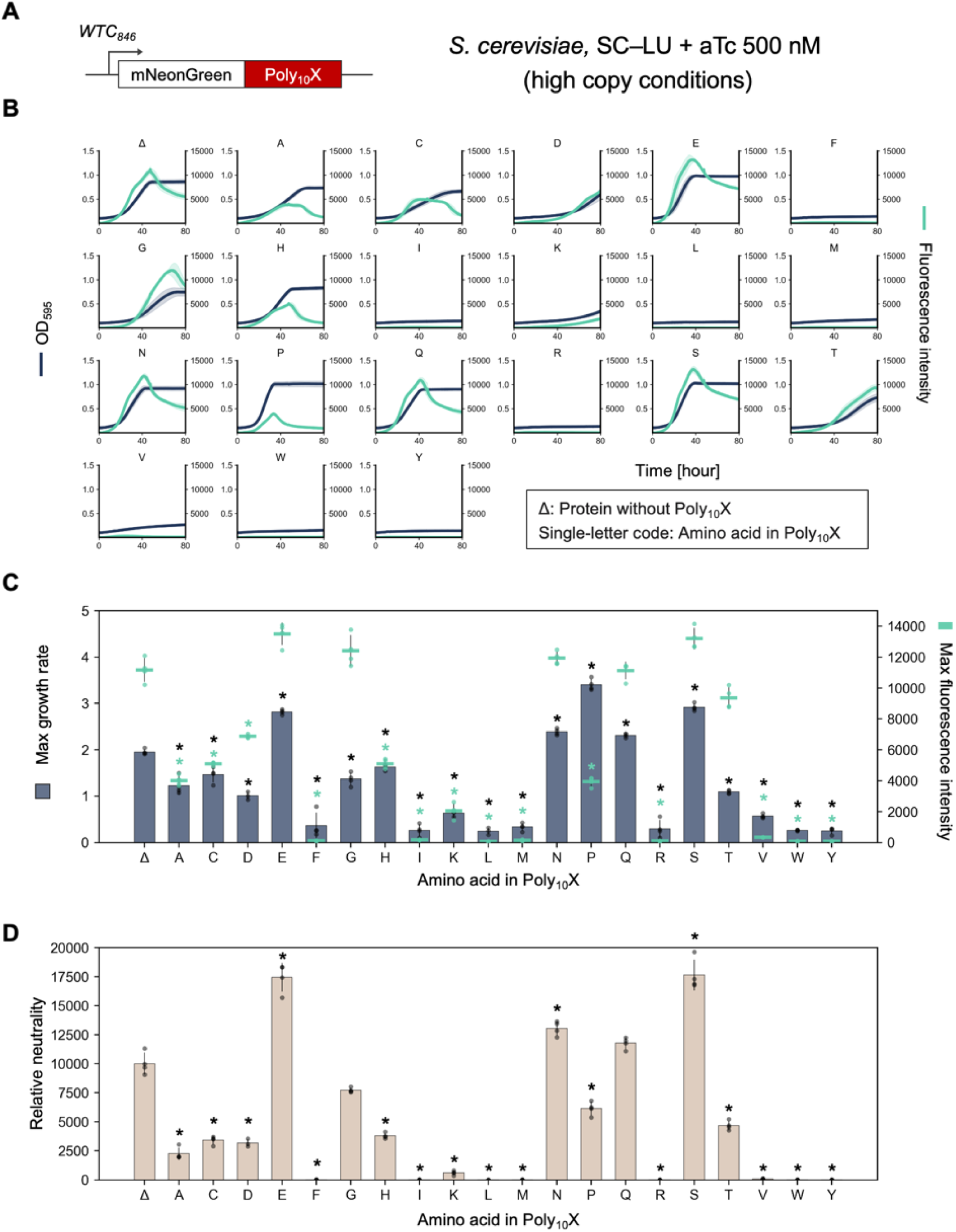
Neutrality of C-terminal Poly_10_X fusions to mNeonGreen in yeast. **A)** Schematic representation of the expression constructs. Poly_10_X was fused to the C-terminus of mNeonGreen and expressed under the control of the *WTC_846_* promoter in *S. cerevisiae*. **B)** Growth and fluorescence curves of *S. cerevisiae* cells expressing mNeonGreen–Poly_10_X, measured using the gTOW method in SC–LU medium at 30 °C. Curves represent the mean values from at least three biological replicates, and shaded regions indicate the standard deviation (SD). **C, D)** Maximum growth rate and maximum fluorescence intensity of *S. cerevisiae* cells overexpressing mNeonGreen–Poly_10_X in SC–LU medium at 30 °C, and the calculated relative neutrality (**D**). Bars, dots, and error bars represent the mean, individual data points, and standard deviation from at least three biological replicates. Asterisks indicate significant differences in maximum growth rate, maximum fluorescence intensity, and relative neutrality, respectively, compared with the control (Δ) (*p* < 0.05, Student’s *t*-test with Bonferroni correction).

**Figure S7.**
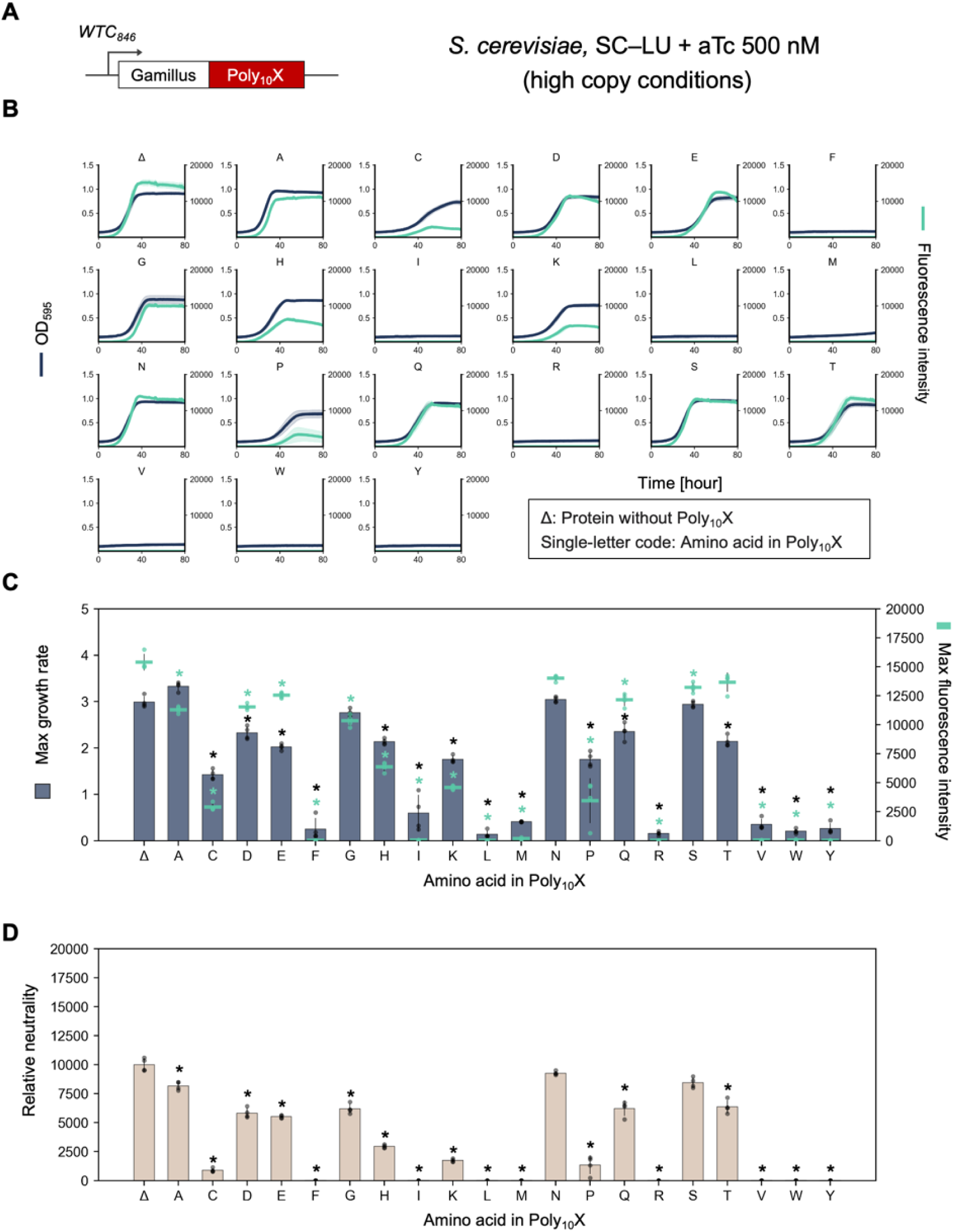
Neutrality of C-terminal Poly_10_X fusions to Gamillus in yeast. **A)** Schematic representation of the expression constructs. Poly_10_X were fused to the C-terminus of Gamillus and expressed under the control of the *WTC_846_* promoter in *S. cerevisiae*. **B)** Growth and fluorescence curves of *S. cerevisiae* cells expressing Gamillus–Poly_10_X, measured using the gTOW method in SC–LU medium at 30 °C. Curves represent the mean values from at least three biological replicates, and shaded regions indicate the standard deviation (SD). **C, D)** Maximum growth rate and maximum fluorescence intensity of *S. cerevisiae* cells overexpressing Gamillus–Poly_10_X in SC–LU medium at 30 °C, and the calculated relative neutrality (**D**). Bars, dots, and error bars represent the mean, individual data points, and standard deviation from at least three biological replicates. Asterisks indicate significant differences in maximum growth rate, maximum fluorescence intensity, and relative neutrality, respectively, compared with the control (Δ) (*p* < 0.05, Student’s *t*-test with Bonferroni correction).

**Figure S8.**
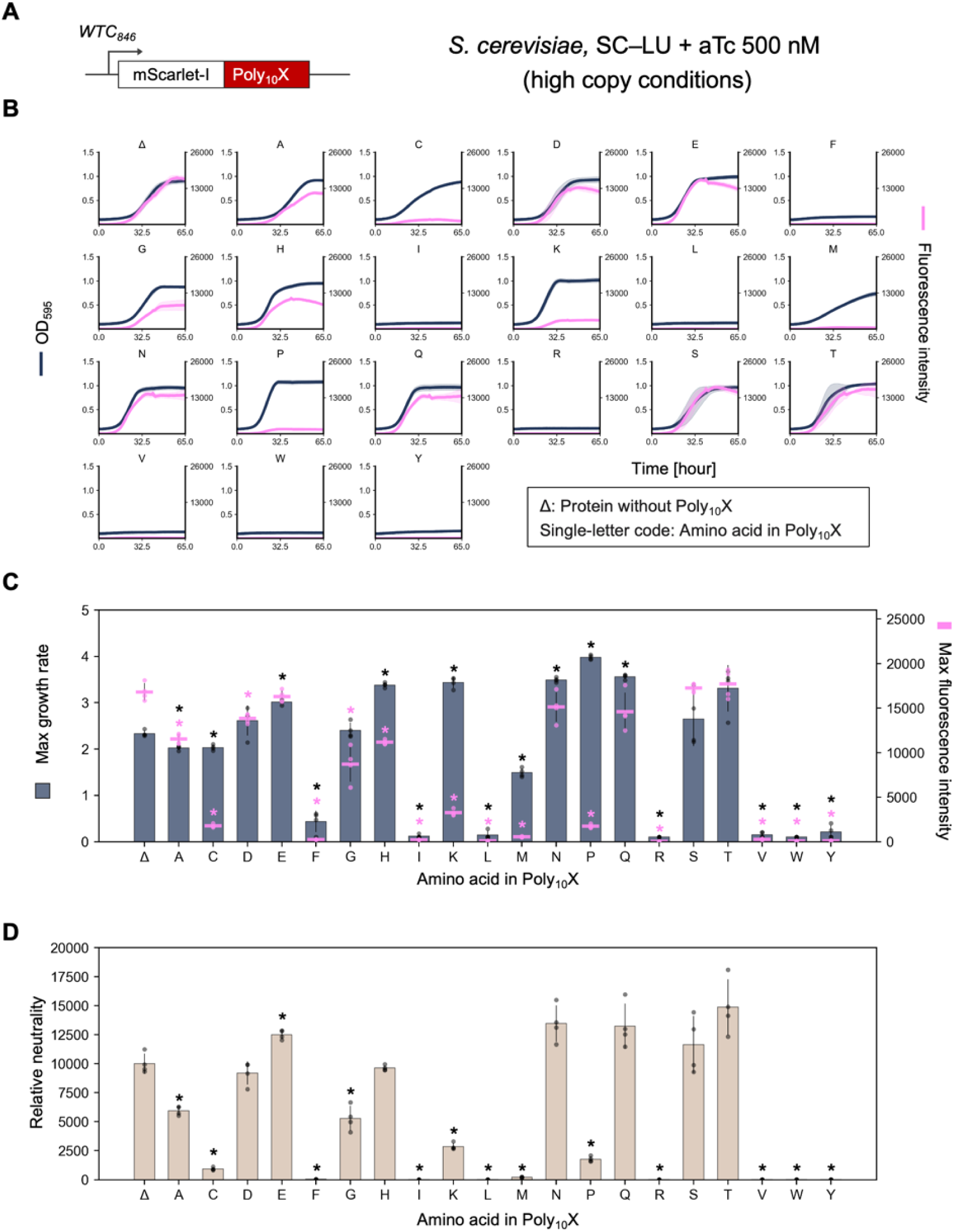
Neutrality of C-terminal Poly_10_X fusions to mScarlet-I in yeast. **A)** Schematic representation of the expression constructs. Poly_10_X was fused to the C-terminus of mScarlet-I and expressed under the control of the *WTC_846_* promoter in *S. cerevisiae*. **B)** Growth and fluorescence curves of *S. cerevisiae* cells expressing mScarlet-I–Poly_10_X, measured using the gTOW method in SC–LU medium at 30 °C. Curves represent the mean values from at least three biological replicates, and shaded regions indicate the standard deviation (SD). **C, D)** Maximum growth rate and maximum fluorescence intensity of *S. cerevisiae* cells overexpressing mScarlet-I–Poly_10_X in SC–LU medium at 30 °C, and the calculated relative neutrality (**D**). Bars, dots, and error bars represent the mean, individual data points, and standard deviation from at least three biological replicates. Asterisks indicate significant differences in maximum growth rate, maximum fluorescence intensity, and relative neutrality, respectively, compared with the control (Δ) (*p* < 0.05, Student’s *t*-test with Bonferroni correction).

**Figure S9.**
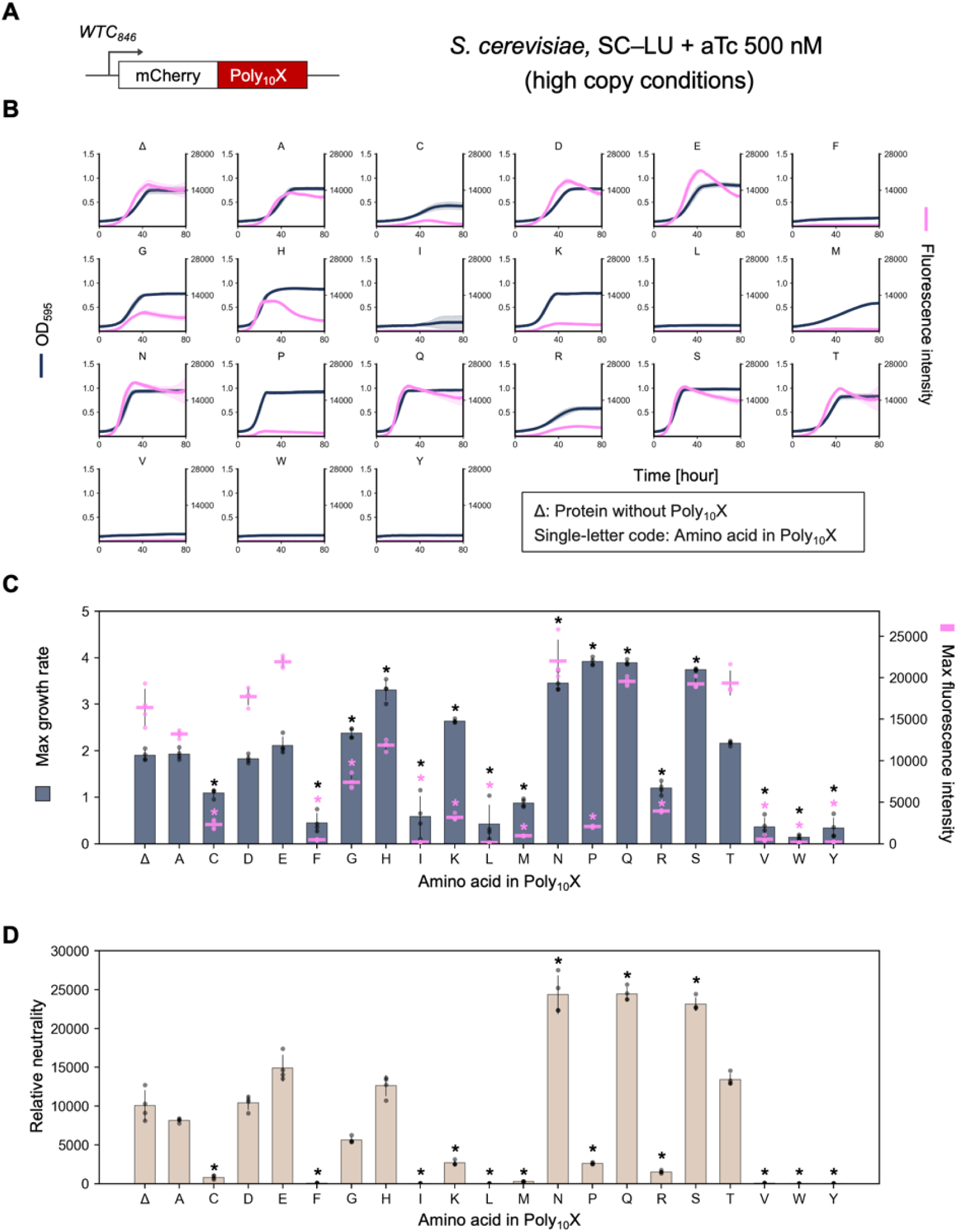
Neutrality of C-terminal Poly_10_X fusions to mCherry in yeast. **A)** Schematic representation of the expression constructs. Poly_10_X was fused to the C-terminus of mCherry and expressed under the control of the *WTC_846_* promoter in *S. cerevisiae*. **B)** Growth and fluorescence curves of *S. cerevisiae* cells expressing mCherry–Poly_10_X, measured using the gTOW method in SC–LU medium at 30 °C. Curves represent the mean values from at least three biological replicates, and shaded regions indicate the standard deviation (SD). **C, D)** Maximum growth rate and maximum fluorescence intensity of *S. cerevisiae* cells overexpressing mCherry–Poly_10_X in SC–LU medium at 30 °C, and the calculated relative neutrality (**D**). Bars, dots, and error bars represent the mean, individual data points, and standard deviation from at least three biological replicates. Asterisks indicate significant differences in maximum growth rate, maximum fluorescence intensity, and relative neutrality, respectively, compared with the control (Δ) (*p* < 0.05, Student’s *t*-test with Bonferroni correction).

**Figure S10.**
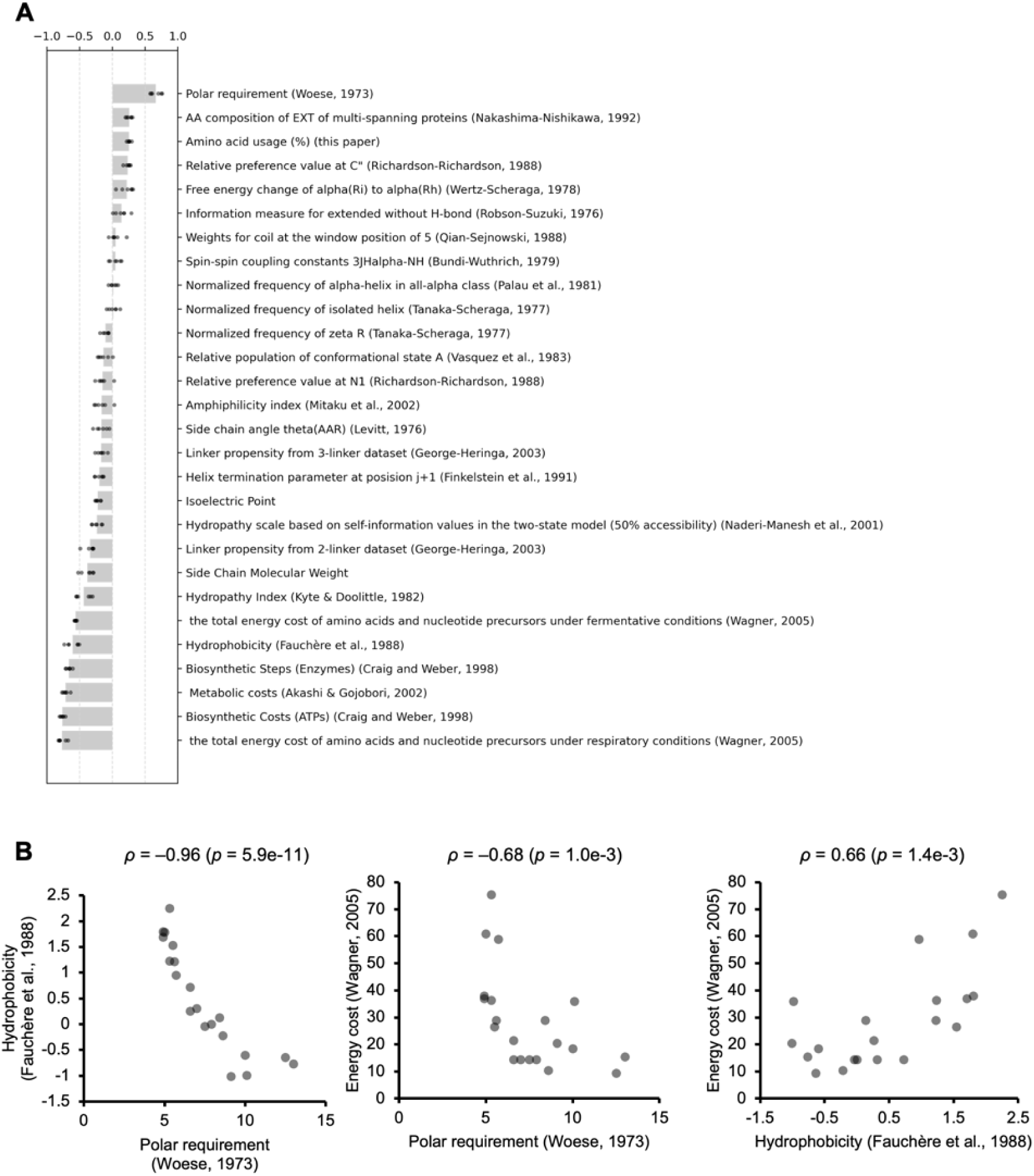
Correlation analysis between Poly_10_X neutrality and amino acid properties. **A)** Correlation between Poly_10_X neutrality and various amino acid indices (physicochemical and usage-related properties). Each dot and bar represents individual Spearman’s rank correlation coefficients and their mean values calculated across six fluorescent proteins. **B)** Relationships among amino acid indices that showed strong correlations with Poly_10_X relative neutrality. Spearman’s rank correlation coefficient (*ρ*) and its *p*-value are shown.

**Figure S11.**
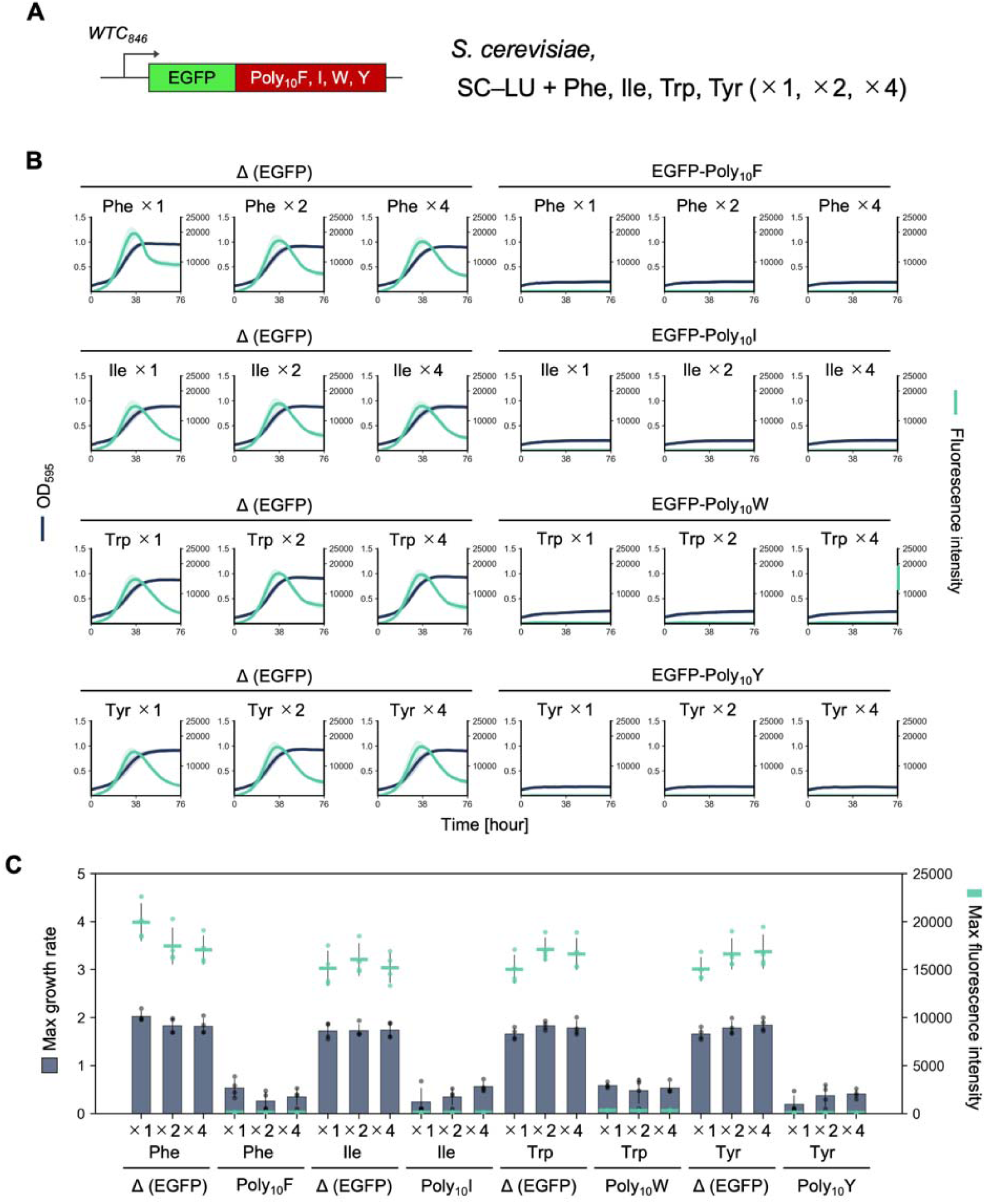
Effects of supplemented amino acids on the harmfulness of high-biosynthetic-cost Poly_10_X repeats. **A)** Schematic representation of the expression constructs. Poly_10_F, Poly_10_I, Poly_10_W, and Poly_10_Y were fused to the C-terminus of EGFP and expressed under the control of the *WTC_846_* promoter in *S. cerevisiae*. **B)** Growth and fluorescence curves of *S. cerevisiae* cells expressing EGFP–Poly_10_X, measured using the gTOW method in SC–LU medium supplemented with additional amino acids (×1, ×2, ×4) at 30 °C. Curves represent the mean values from four biological replicates, and shaded regions indicate the standard deviation (SD). **C)** Maximum growth rate and maximum fluorescence intensity of *S. cerevisiae* cells overexpressing EGFP–Poly_10_X in SC–LU medium supplemented with additional amino acids (×1, ×2, ×4) at 30 °C. Bars, dots, and error bars represent the mean, individual data points, and standard deviation from four biological replicates. Maximum growth rate and maximum fluorescence intensity of *S. cerevisiae* cells overexpressing EGFP without Poly_10_X (Δ) or with C-terminal Poly_10_F, Poly_10_I, Poly_10_W, or Poly_10_Y fusions in SC–LU medium at 30 °C. Indicated amino acid was supplemented to the medium at standard (×1), ×2, or ×4 concentrations. Bars, dots, and error bars represent the mean, individual values, and standard deviation from four biological replicates.

**Figure S12.**
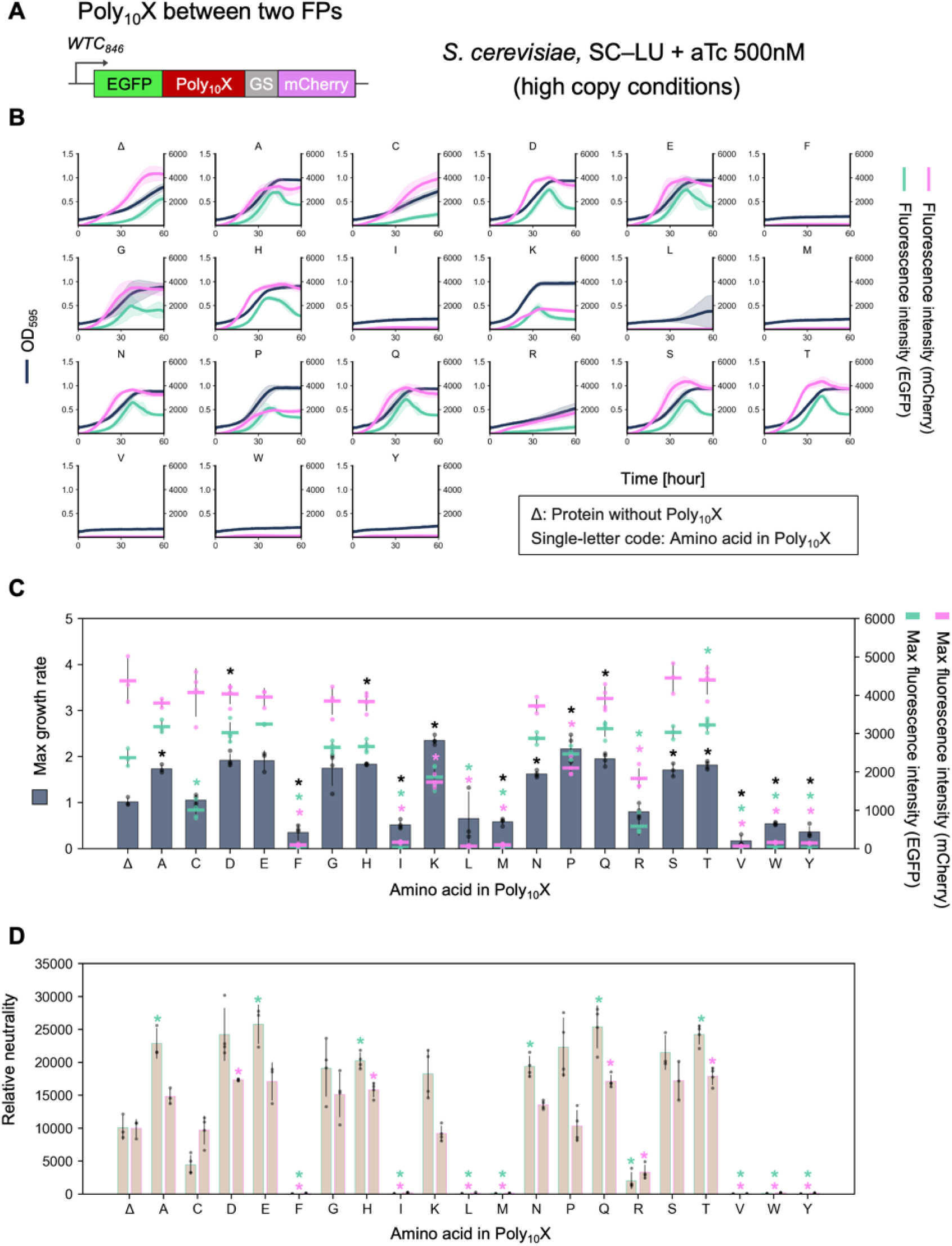
Neutrality of Poly_10_X Insertions between two FPs in yeast. **A)** Schematic representation of the expression constructs. Poly_10_X were inserted between EGFP and mCherry and expressed under the control of the *WTC_846_* promoter in *S. cerevisiae*. **B)** Growth and fluorescence curves of *S. cerevisiae* cells expressing EGFP–Poly_10_X–GSlinker–mCherry, measured using the gTOW method in SC–LU medium at 30 °C. Curves represent the mean values from at least three biological replicates, and shaded regions indicate the standard deviation (SD). **C, D)** Maximum growth rate and maximum fluorescence intensity of *S. cerevisiae* cells overexpressing EGFP–Poly_10_X–GSlinker–mCherry in SC–LU medium at 30 °C, and the calculated relative neutrality (**D**). Bars, dots, and error bars represent the mean, individual data points, and standard deviation from at least three biological replicates. Asterisks indicate significant differences in maximum growth rate, maximum fluorescence intensity, and relative neutrality, respectively, compared with the control (Δ) (*p* < 0.05, Student’s *t*-test with Bonferroni correction).

**Figure S13.**
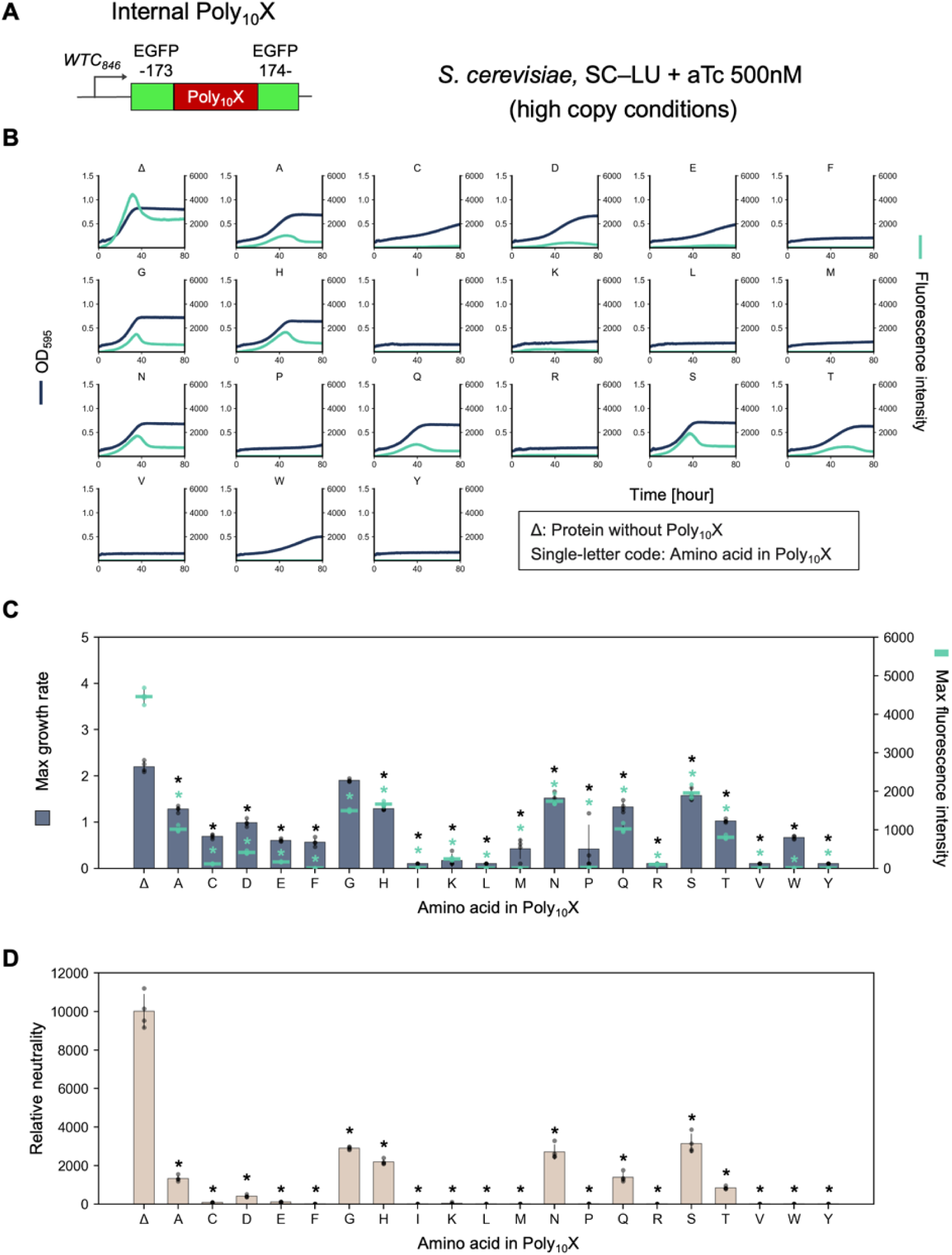
Neutrality of Poly_10_X Insertions within EGFP in yeast. **A)** Schematic representation of the expression constructs. Poly_10_X was inserted into an internal loop of EGFP between residues 173 and 174, and expressed under the control of the *WTC_846_* promoter in *S. cerevisiae*. **B)** Growth and fluorescence curves of *S. cerevisiae* cells expressing EGFP_173_–Poly_10_X–_174_EGFP, measured using the gTOW method in SC–LU medium at 30 °C. Curves represent the mean values from at least three biological replicates, and shaded regions indicate the standard deviation (SD). **C, D)** Maximum growth rate and maximum fluorescence intensity of *S. cerevisiae* cells overexpressing EGFP_173_–Poly_10_X–_174_EGFP in SC–LU medium at 30 °C, and the calculated relative neutrality (**D**). Bars, dots, and error bars represent the mean, individual data points, and standard deviation from at least three biological replicates. Asterisks indicate significant differences in maximum growth rate, maximum fluorescence intensity, and relative neutrality, respectively, compared with the control (Δ) (*p* < 0.05, Student’s *t*-test with Bonferroni correction).

**Figure S14.**
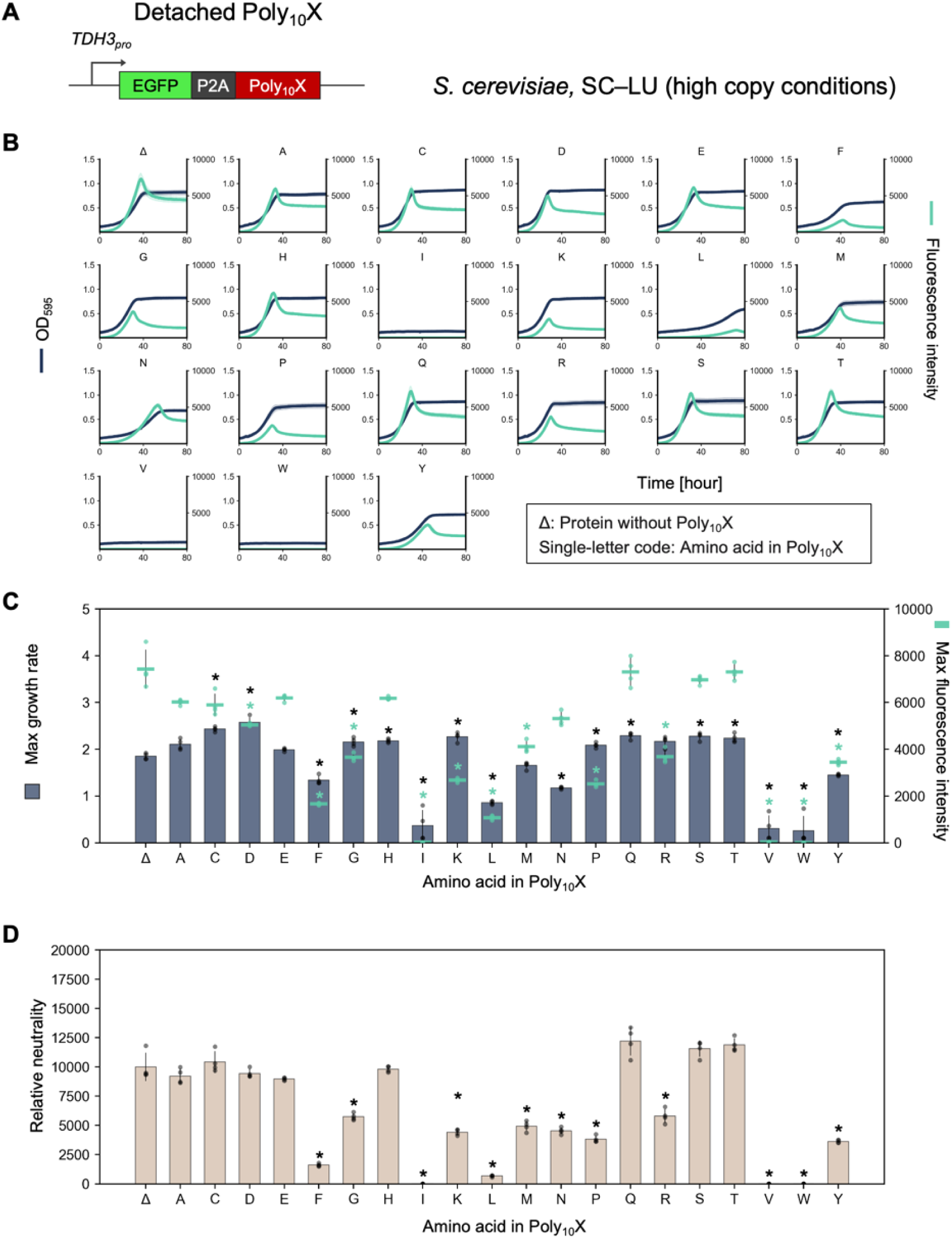
Neutrality of Poly_10_X detached from EGFP via P2A in Yeast. **A)** Schematic representation of the expression constructs. A self-cleaving P2A sequence was inserted between EGFP and Poly_10_X, allowing Poly_10_X to be detached from EGFP during translation. The construct was expressed under the control of the *TDH3* promoter in *S. cerevisiae*. **B)** Growth and fluorescence curves of *S. cerevisiae* cells expressing EGFP–P2A–Poly_10_X, measured using the gTOW method in SC–LU medium at 30 °C. Curves represent the mean values from four biological replicates, and shaded regions indicate the standard deviation (SD). **C, D)** Maximum growth rate and maximum fluorescence intensity of *S. cerevisiae* cells overexpressing EGFP–P2A–Poly_10_X in SC–LU medium at 30 °C, and the calculated relative neutrality (**D**). Bars, dots, and error bars represent the mean, individual data points, and standard deviation from four biological replicates. Asterisks indicate significant differences in maximum growth rate, maximum fluorescence intensity, and relative neutrality, respectively, compared with the control (Δ) (*p* < 0.05, Student’s *t*-test with Bonferroni correction).

**Figure S15.**
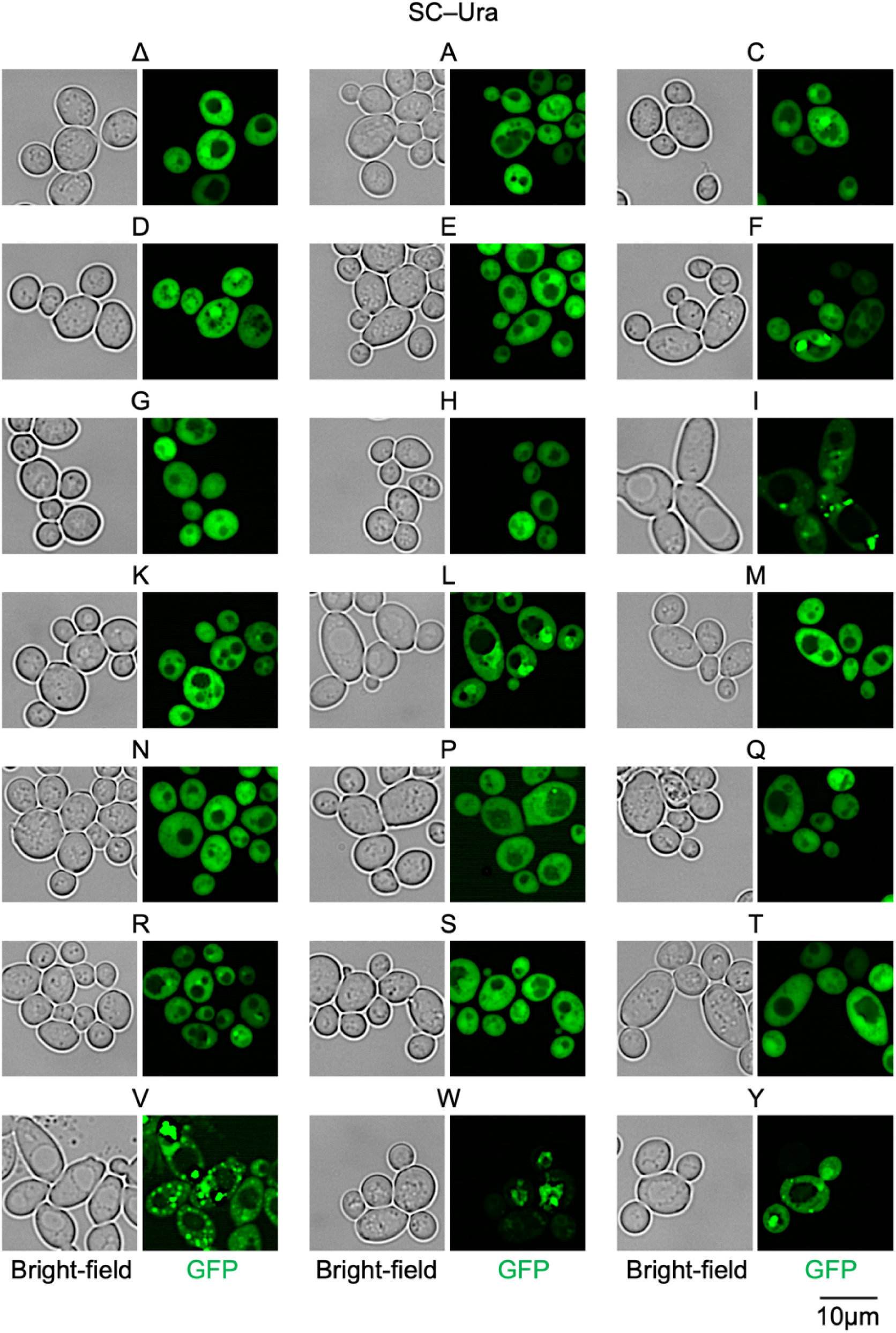
Fluorescence microscopy of yeast cells expressing EGFP–Poly_10_X (low-copy conditions). Fluorescence microscopy images of *S. cerevisiae* cells expressing EGFP–Poly_10_X under the control of *TDH3_pro_*. Cells were pre-cultured in SC–U medium and subsequently cultured overnight in the same medium before imaging. Brightness and contrast were adjusted to clearly visualize cell morphology. For each sample, the bright-field image (left) and the corresponding GFP fluorescence image (right) are shown. Single-letter codes indicate the amino acid repeated in Poly_10_X. Δ represents the protein without a Poly_10_X fusion.

**Figure S16.**
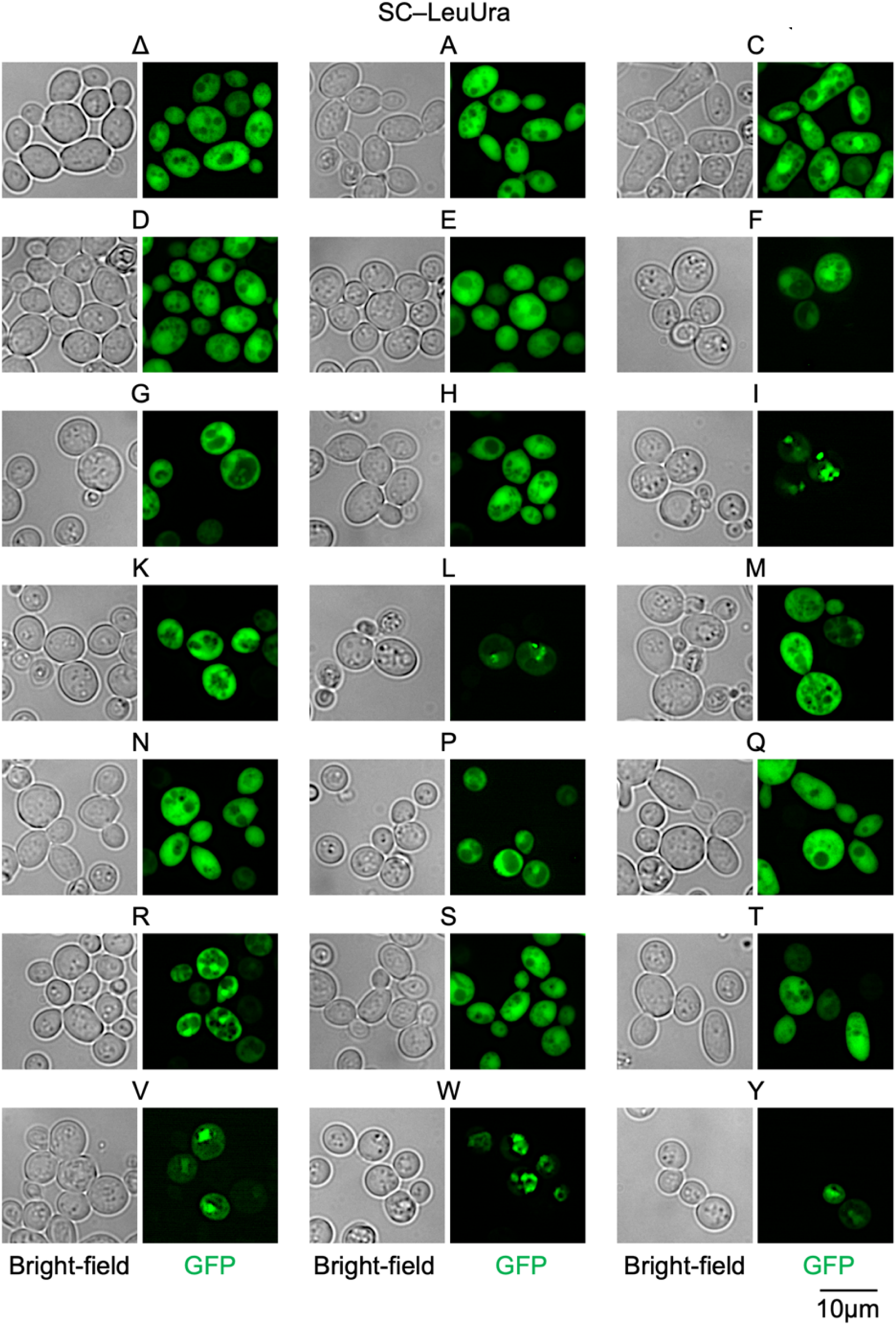
Fluorescence microscopy of yeast cells expressing EGFP–Poly_10_X (High-copy expression). Fluorescence microscopy images of *S. cerevisiae* cells expressing EGFP–Poly_10_X under the control of *TDH3_pro_*. Cells were pre-cultured in SC–U medium and subsequently cultured overnight in SC–LU medium before imaging. Brightness and contrast were adjusted to clearly visualize cell morphology. For each sample, the bright-field image (left) and the corresponding GFP fluorescence image (right) are shown. Single-letter codes indicate the amino acid repeated in Poly_10_X. Δ represents the protein without a Poly_10_X fusion.

**Figure S17.**
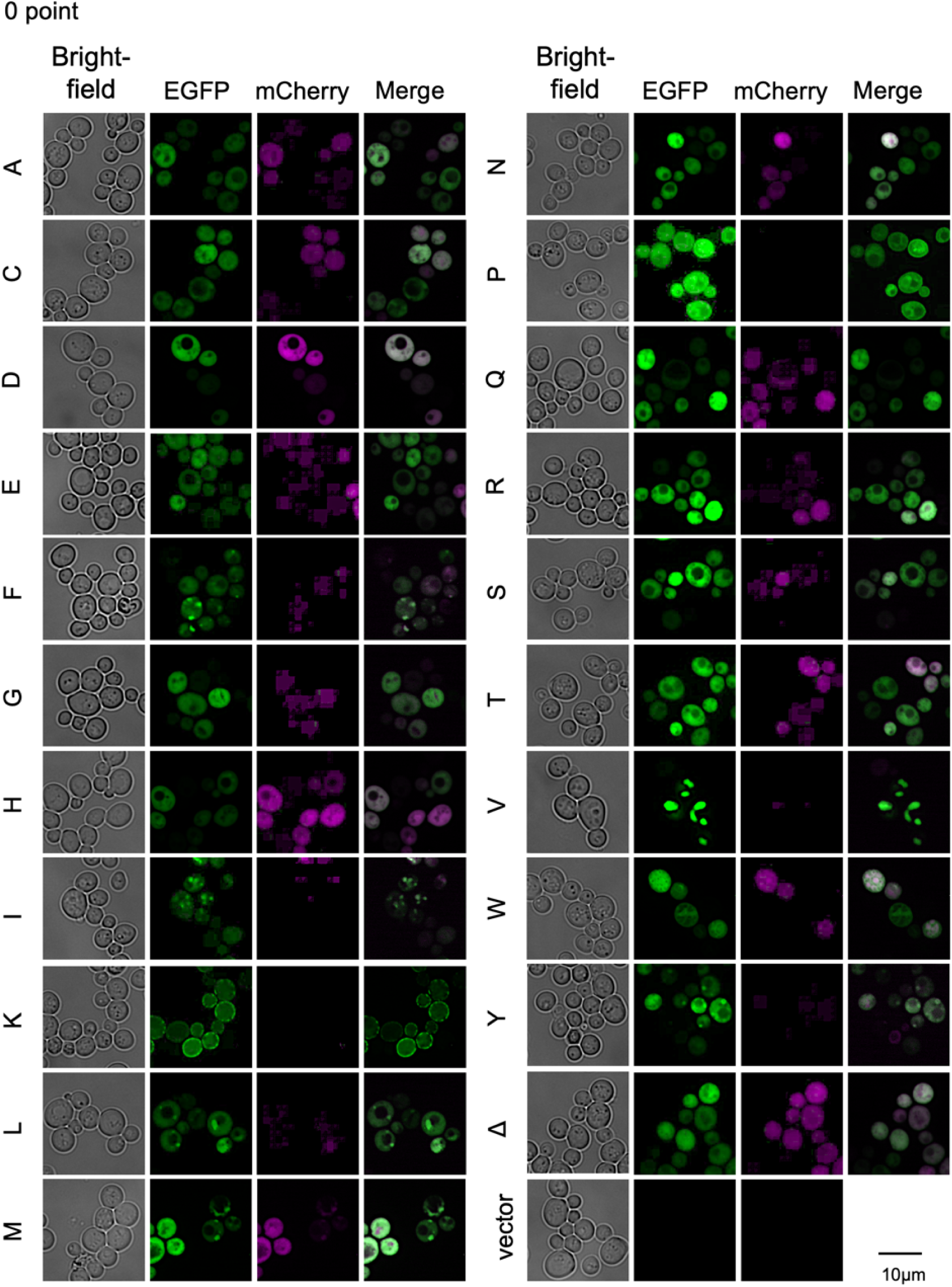
Fluorescence microscopy of EGFP–Poly_10_X–mCherry expression immediately after induction in yeast. Fluorescence microscopy images of *S. cerevisiae* cells expressing EGFP–Poly_10_X–GSlinker–mCherry under the control of the *WTC_84_*_6_ promoter immediately after aTc induction. Cells were pre-cultured in SC–U medium and subsequently cultured overnight in SC–LU medium before imaging. Brightness and contrast were adjusted to clearly visualize cell morphology. For each sample, the bright-field image (left), GFP fluorescence image (center left), RFP fluorescence image (center right), and the merged GFP/RFP image (right) are shown. Single-letter codes indicate the amino acid repeated in Poly_10_X. Δ represents the protein without a Poly_10_X fusion.

**Figure S18.**
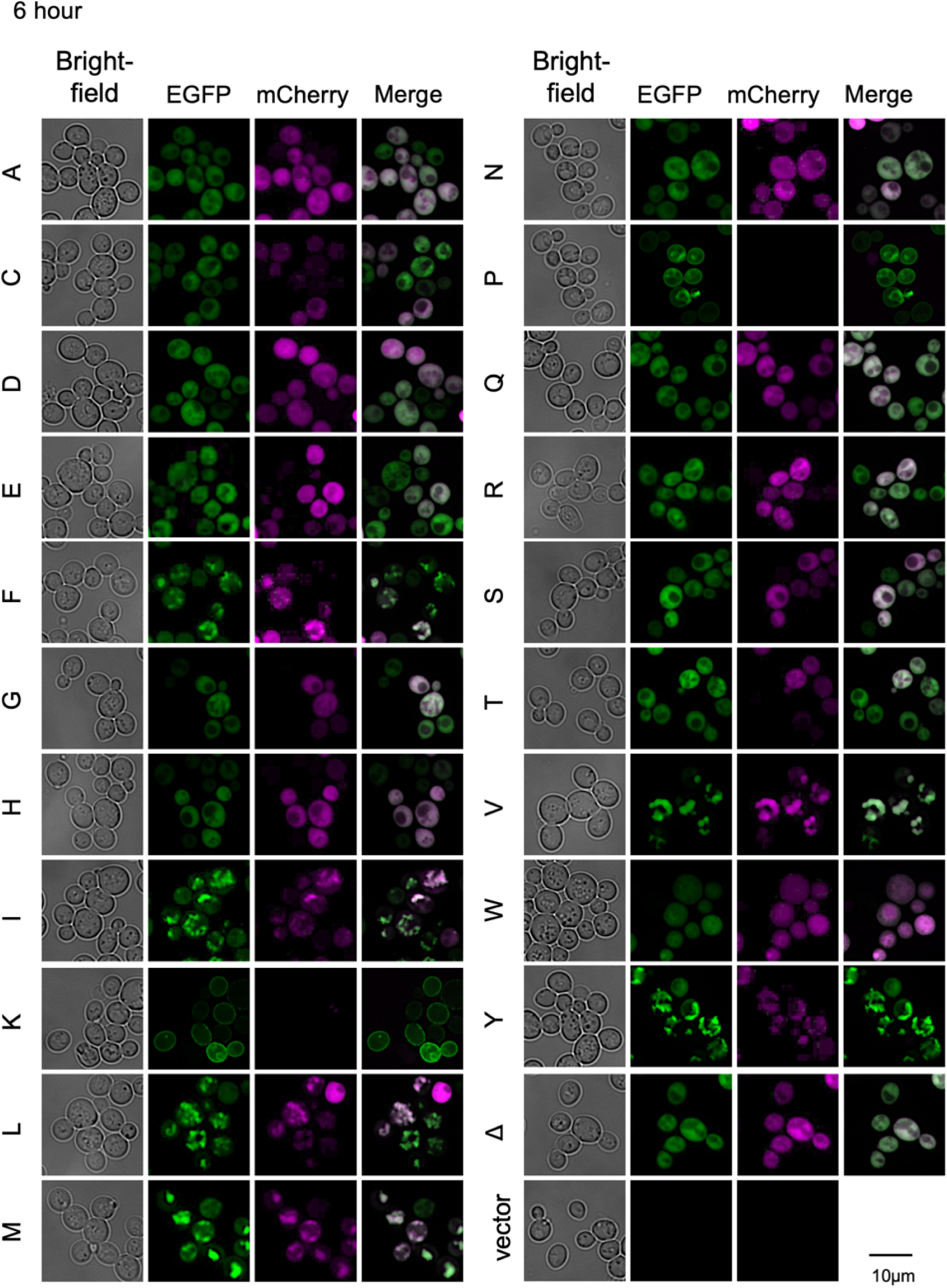
Fluorescence microscopy of EGFP–Poly_10_X–mCherry expression six hours after aTc induction in yeast Fluorescence microscopy images of *S. cerevisiae* cells expressing EGFP–Poly_10_X–GSlinker–mCherry under the control of the *WTC_84_*_6_ promoter six hours after aTc induction. Cells were pre-cultured in SC–U medium and subsequently cultured overnight in SC–LU medium before imaging. Brightness and contrast were adjusted to clearly visualize cell morphology. For each sample, the bright-field image (left), GFP fluorescence image (center left), RFP fluorescence image (center right), and the merged GFP/RFP image (right) are shown. Single-letter codes indicate the amino acid repeated in Poly_10_X. Δ represents the protein without a Poly_10_X fusion.

**Figure S19.**
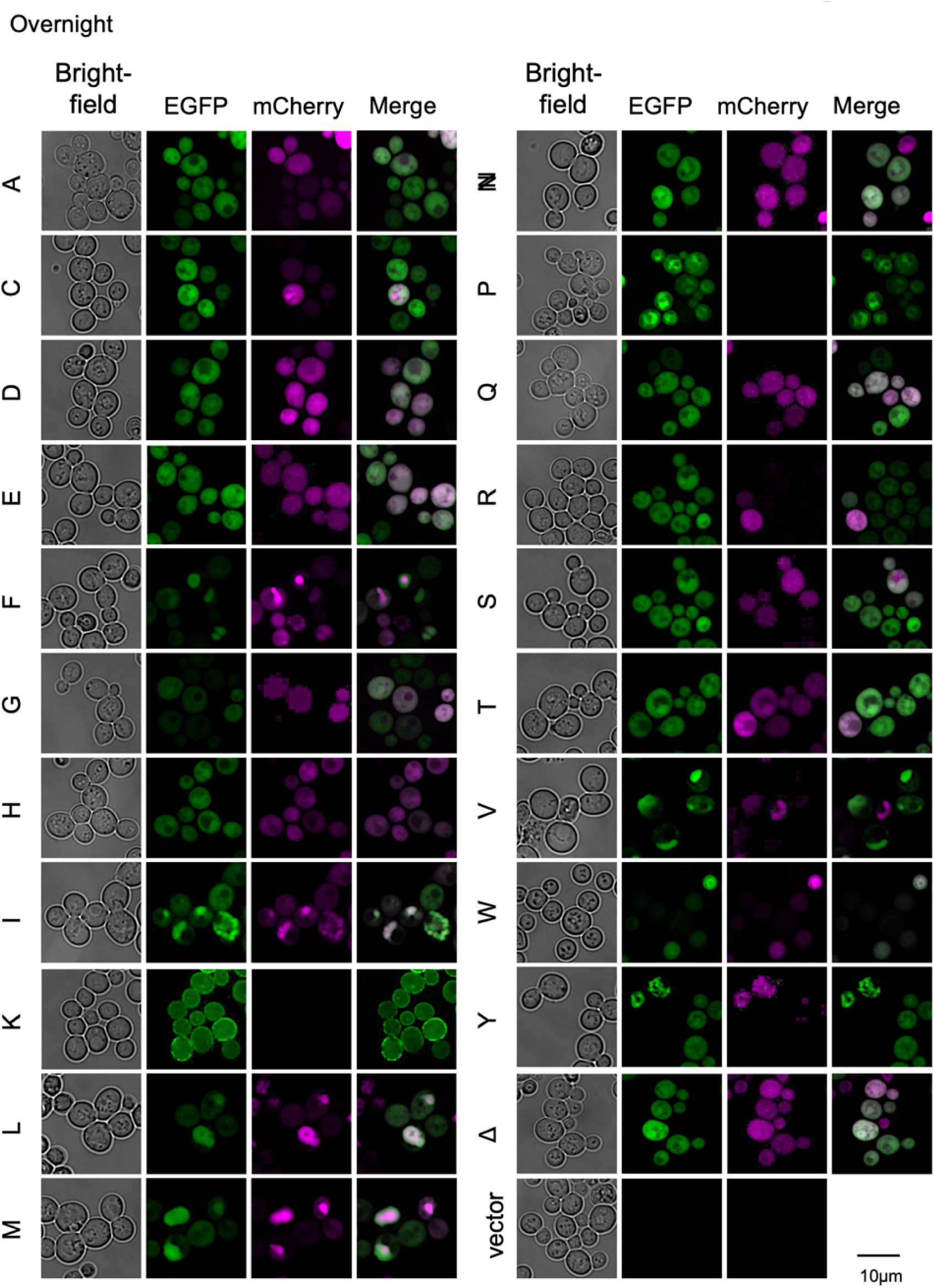
Fluorescence microscopy of EGFP–Poly_10_X–mCherry expression overnight after aTc induction in yeast. Fluorescence microscopy images of *S. cerevisiae* cells expressing EGFP–Poly_10_X–GSlinker–mCherry under the control of the *WTC_84_*_6_ promoter overnight after aTc induction. Cells were pre-cultured in SC–U medium and subsequently cultured overnight in SC–LU medium before imaging. Brightness and contrast were adjusted to clearly visualize cell morphology. For each sample, the bright-field image (left), GFP fluorescence image (center left), RFP fluorescence image (center right), and the merged GFP/RFP image (right) are shown. Single-letter codes indicate the amino acid repeated in Poly_10_X. Δ represents the protein without a Poly_10_X fusion.

**Figure S20.**
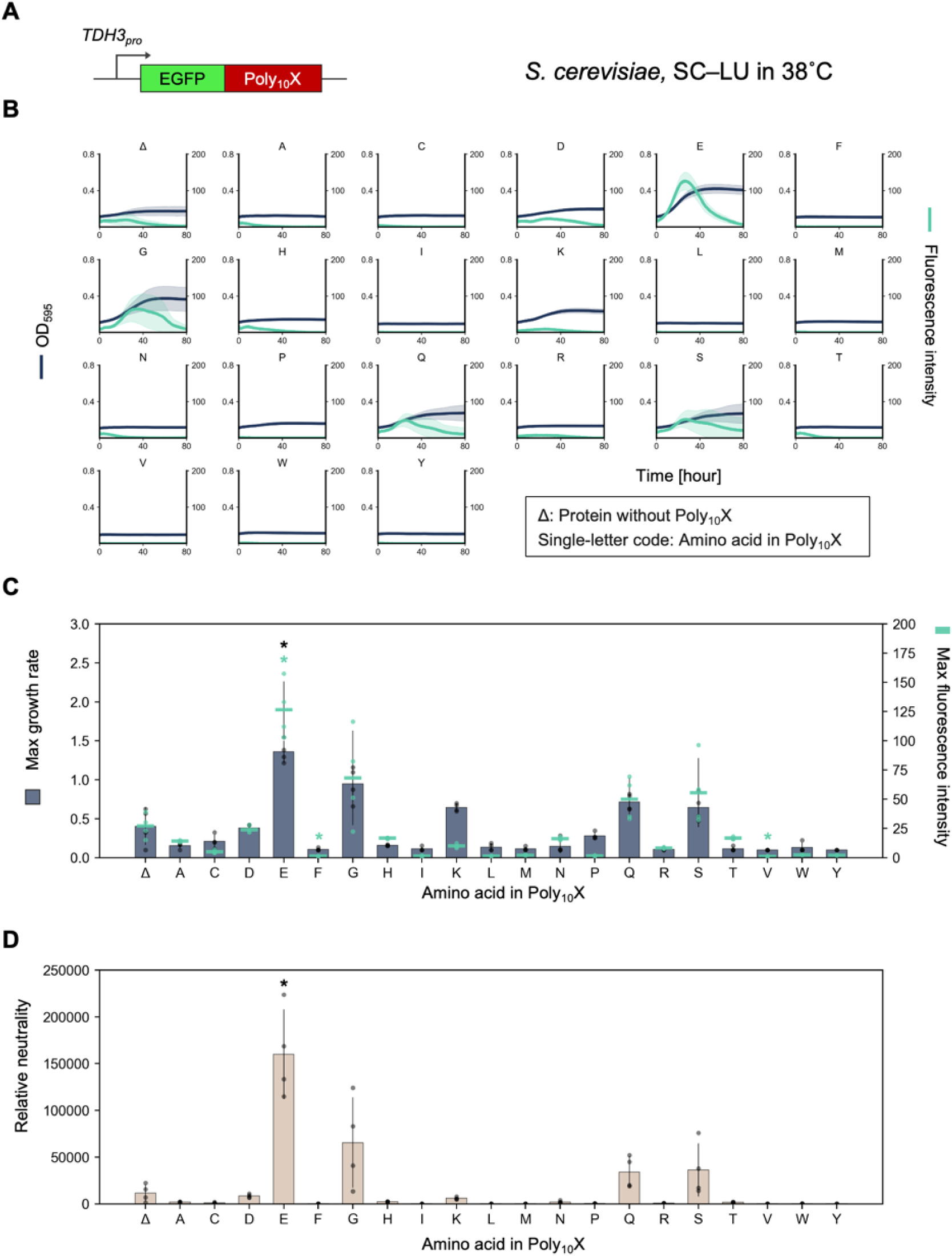
Neutrality of C-terminal Poly_10_X fusions to EGFP in yeast under heat-stress conditions. **A)** Schematic representation of the expression constructs. Poly_10_X was fused to the C-terminus of EGFP and expressed under the *TDH3* promoter in *S. cerevisiae*. **B)** Growth and fluorescence curves of *S. cerevisiae* cells expressing EGFP–Poly_10_X, measured using the gTOW method in SC–LU medium at 38 °C. Curves represent the mean values from at least three biological replicates, and shaded regions indicate the standard deviation (SD). **C, D)** Maximum growth rate and maximum fluorescence intensity of *S. cerevisiae* cells overexpressing EGFP–Poly_10_X in SC–LU medium at 38 °C, and the calculated relative neutrality (**D**). Bars, dots, and error bars represent the mean, individual data points, and standard deviation from at least three biological replicates. Asterisks indicate significant differences in maximum growth rate, maximum fluorescence intensity, and relative neutrality, respectively, compared with the control (Δ) (*p* < 0.05, Student’s *t*-test with Bonferroni correction).

**Figure S21.**
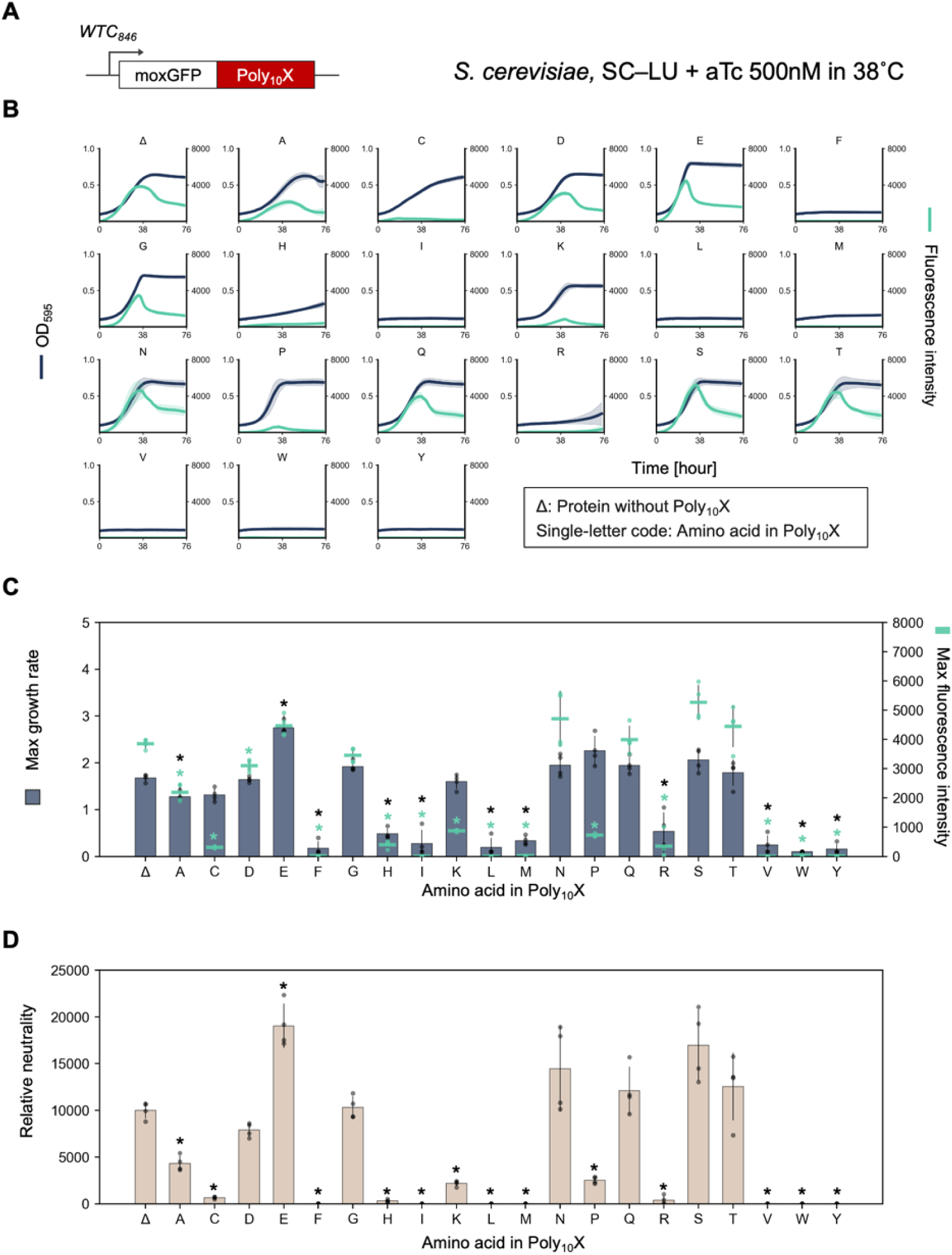
Neutrality of C-terminal Poly_10_X fusions to moxGFP in yeast under heat-stress conditions. **A)** Schematic representation of the expression constructs. Poly_10_X were fused to the C-terminus of moxGFP and expressed under the *WTC_846_* promoter in *S. cerevisiae*. **B)** Growth and fluorescence curves of *S. cerevisiae* cells expressing moxGFP–Poly_10_X, measured using the gTOW method in SC–LU medium at 38 °C. Curves represent the mean values from at least three biological replicates, and shaded regions indicate the standard deviation (SD). **C, D)** Maximum growth rate and maximum fluorescence intensity of *S. cerevisiae* cells overexpressing moxGFP–Poly_10_X in SC–LU medium at 38 °C, and the calculated relative neutrality (**D**). Bars, dots, and error bars represent the mean, individual data points, and standard deviation from at least three biological replicates. Asterisks indicate significant differences in maximum growth rate, maximum fluorescence intensity, and relative neutrality, respectively, compared with the control (Δ) (*p* < 0.05, Student’s *t*-test with Bonferroni correction).

**Figure S22.**
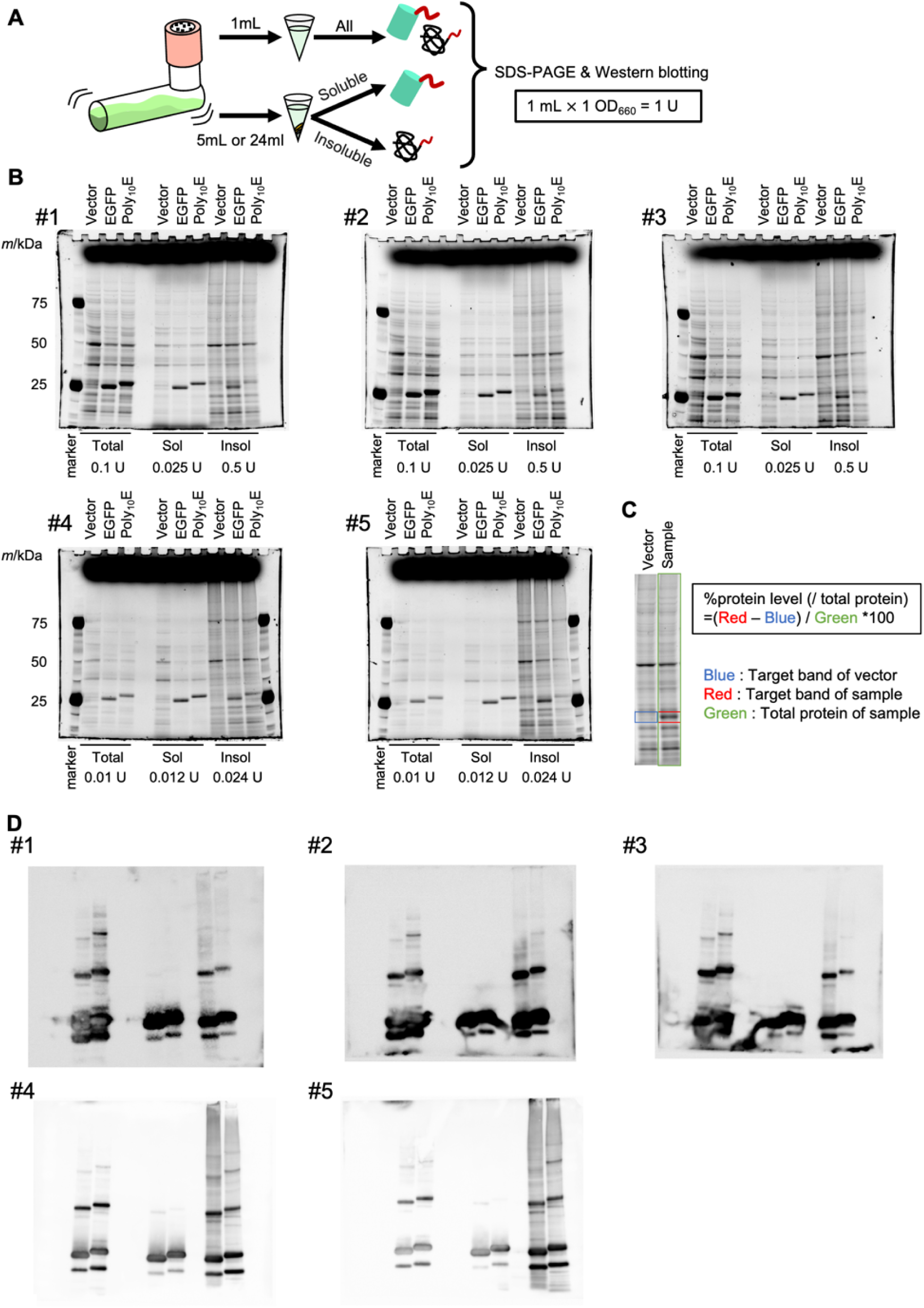
SDS–PAGE and Western blot analysis of EGFP and EGFP–Poly_10_E proteins in yeast **A)** Schematic overview of the protein extraction procedure. A 25 mL culture was collected at approximately OD_660_ = 1. One milliliter was used for total protein extraction, while the remaining 5 or 24 mL was fractionated into soluble and insoluble fractions. Each fraction was analyzed by SDS–PAGE, followed by western blotting. Protein concentrations were normalized based on the total protein amount extracted from cells in 1 mL of culture at OD_660_ = 1, which was defined as 1 unit (1 U). **B)** SDS–PAGE images of proteins extracted from cells with the control vector (Vector) or overexpressing EGFP or EGFP–Poly_10_E (Poly_10_E). Cells were cultured in SC–LU medium and collected at approximately OD_660_ = 1. Lanes correspond to total, soluble (Sol), and insoluble (Insol) fractions (from left to right). Protein bands were visualized using a chemiluminescent detection reagent. Five biological replicates were analyzed. Protein loading concentrations are indicated in the figure. **C)** Quantification method for %protein level from SDS–PAGE images shown in **B**. Signal intensity corresponding to the EGFP band (Red) was corrected by subtracting the background intensity from the same region (Blue), and the resulting value was normalized to the total protein amount in the sample (green) to calculate the percentage. **D)** Western blot images of proteins extracted from cells with the control vector (Vector) or overexpressing EGFP or EGFP–Poly_10_E (Poly_10_E). The same gels used in **B** were transferred to membranes and probed with anti-GFP antibodies. Lanes correspond to total, soluble, and insoluble fractions (from left to right).

**Figure S23.**
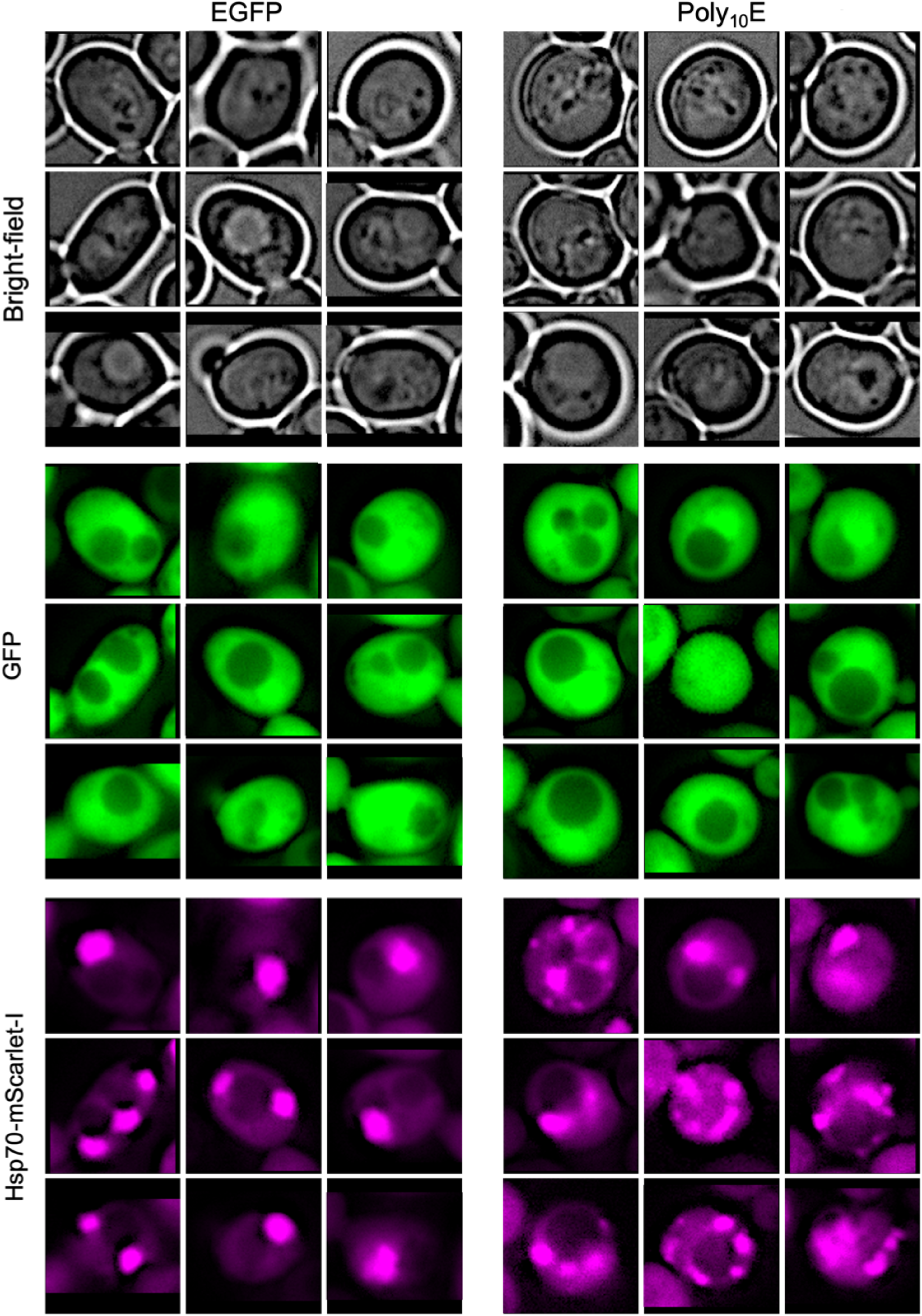
Fluorescence imaging of Hsp70 foci in yeast cells overexpressing EGFP or EGFP–Poly_10_E. Fluorescence microscopy images of Hsp70 foci in cells overexpressing EGFP or EGFP–Poly_10_E under the control of *TDH3_pro_* in SC–LU conditions. Hsp70–mScarlet-I was genomically integrated to visualize Hsp70 aggregate formation. Representative images of nine individual cells are shown, along with the corresponding GFP and RFP fluorescence images. Green indicates EGFP or EGFP–Poly_10_E, and magenta represents the distribution of Hsp70. Image brightness and contrast were adjusted to enhance the visibility of aggregates.

**Figure S24.**
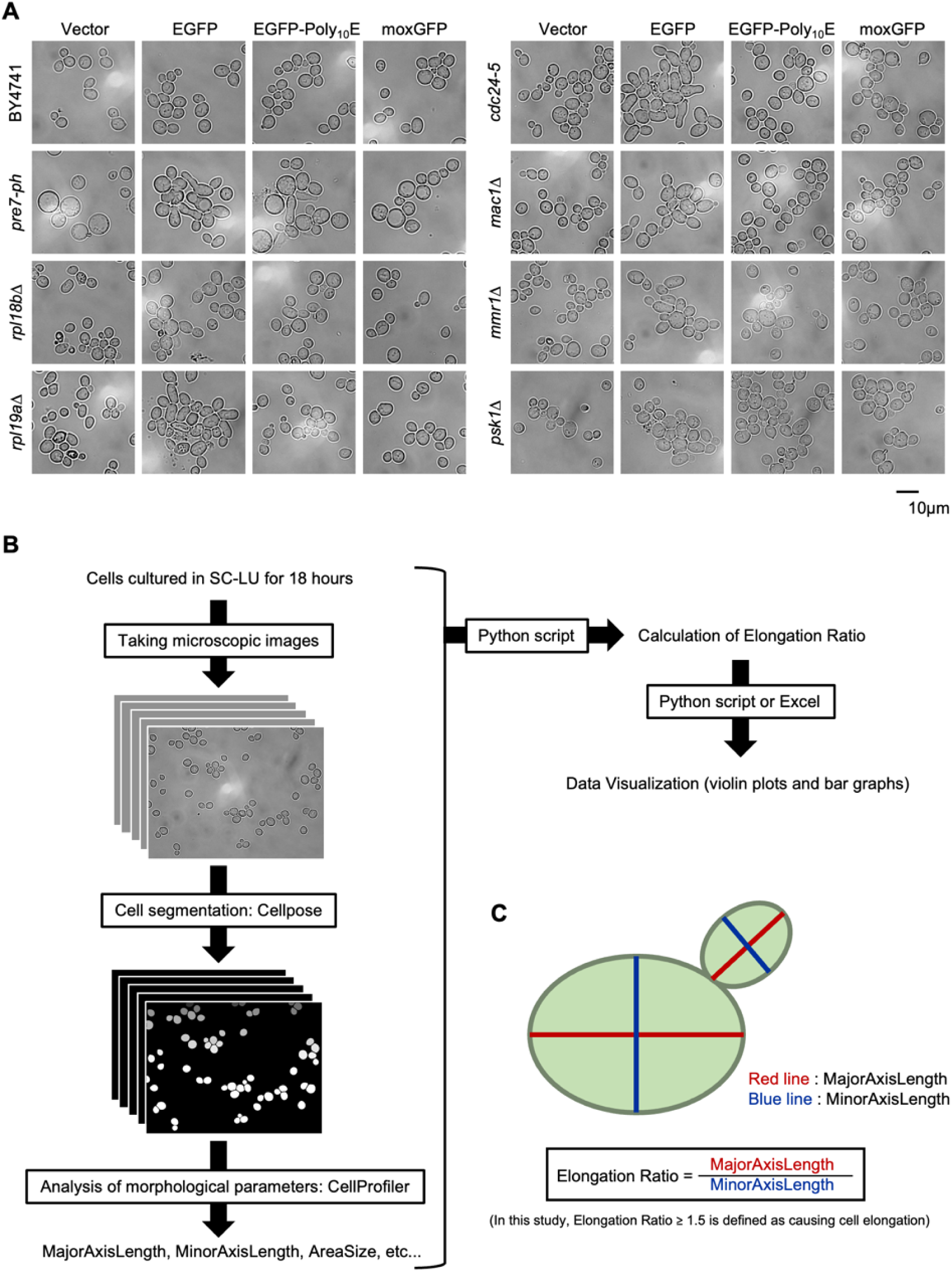
Morphological analysis of yeast strains overexpressing EGFP or EGFP–Poly_10_E **A)** Microscopy images of *S. cerevisiae* strain BY4741 and seven mutant strains (*pre7-ph*, *rpl18bΔ*, *rpl19aΔ*, *cdc24-5*, *mac1Δ*, *mmr1Δ*, and *psk1Δ*) with the control vector (Vector) or overexpressing EGFP, EGFP–Poly_10_E, or moxGFP under the control of *TDH3_pro_*. Cells were pre-cultured in SC–U medium and then cultured for 18 h in SC–LU medium before imaging. Brightness and contrast were adjusted to clearly visualize cell morphology. **B)** Schematic workflow of image analysis. After culturing the cells for 18 h in SC–LU medium, microscopic images were acquired and processed using Cellpose 2.2.3 ^64^ for cell segmentation. The segmented images were then analyzed with the *MeasureObjectSizeShape* module in CellProfiler 4.2.6 ^65^ to measure the major and minor axes of each cell. Elongation ratio (major axis/minor axis) values were calculated using a Python script or Excel and visualized. **C)** Schematic representation of cell elongation analysis. In this study, cells with an elongation ratio ≥ 1.5 were defined as morphologically abnormal.

**Figure S25.**
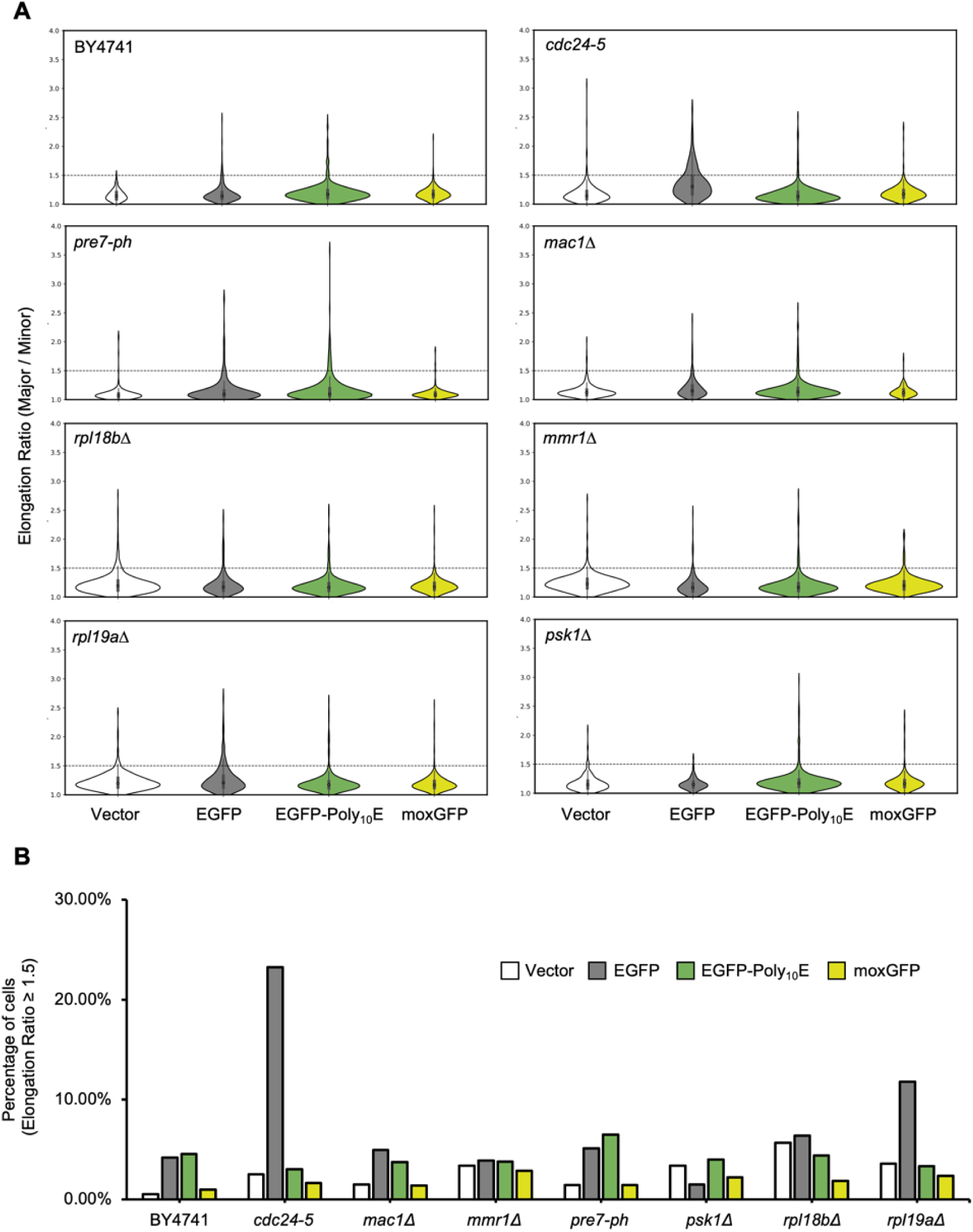
Quantification of cell elongation defects in yeast strains overexpressing EGFP or EGFP–Poly_10_E. **A)** Distribution of elongation ratios in *S. cerevisiae* strain BY4741 and seven mutant strains (*pre7-ph*, *rpl18bΔ*, *rpl19aΔ*, *cdc24-5*, *mac1Δ*, *mmr1Δ*, and *psk1Δ*) with the control vector (Vector) or overexpressing EGFP, EGFP–Poly_10_E, or moxGFP under the control of *TDH3_pro_*. Each violin plot represents the elongation ratio (major axis/minor axis) of individual cells. Wider regions indicate a higher frequency of cells with the corresponding elongation ratio. The dashed horizontal line represents the threshold for morphological abnormality (elongation ratio ≥ 1.5). Morphological parameters were calculated from microscopic images analyzed using Cellpose and CellProfiler as described in Figure S24B and C. **B)** Proportion of morphologically abnormal cells (elongation ratio ≥ 1.5) in each strain. The analysis was performed on a single biological replicate, with at least 197 cells analyzed per condition. Bars indicate the percentage of abnormal cells relative to the total number of segmented cells.

**Figure S26.**
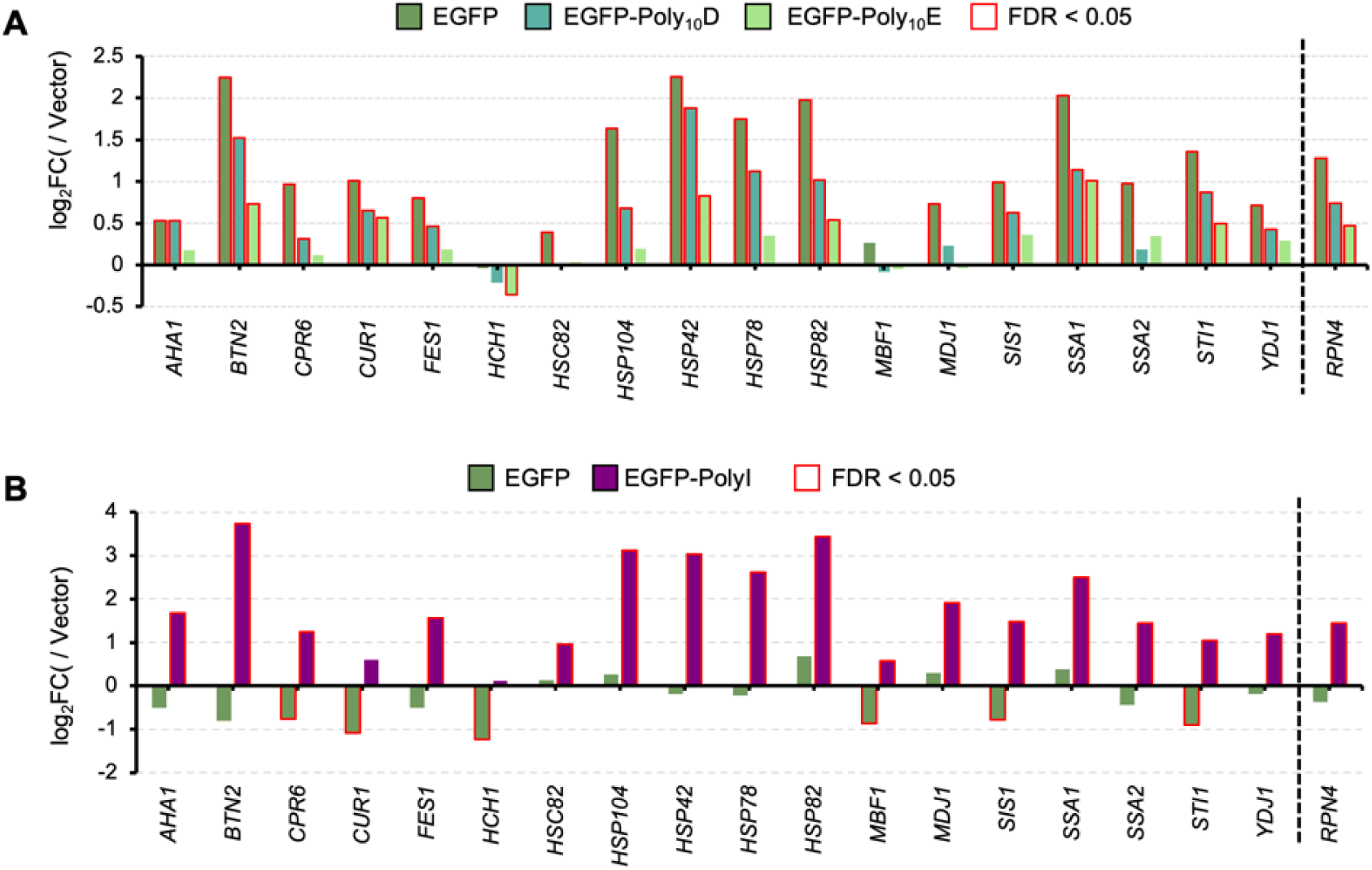
Expression changes of Hsf1-regulated genes and *RPN4* in yeast expressing EGFP-Poly_10_D, EGFP-Poly_10_E, or EGFP-Poly_10_I. **A, B)** Comparison of expression ratios (vs. Vector) for genes regulated by Hsf1 and for *RPN4*, quantified by RNA-seq. In **A**, bars indicate the expression ratios of cells overexpressing EGFP, EGFP–Poly_10_D, and EGFP–Poly_10_E, respectively. In **B**, bars indicate the expression ratios of cells with low-level overexpressing EGFP and EGFP–Poly_10_I, respectively. Outlines denote genes showing significant differential expression compared with the vector control (FDR < 0.05).

**Figure S27.**
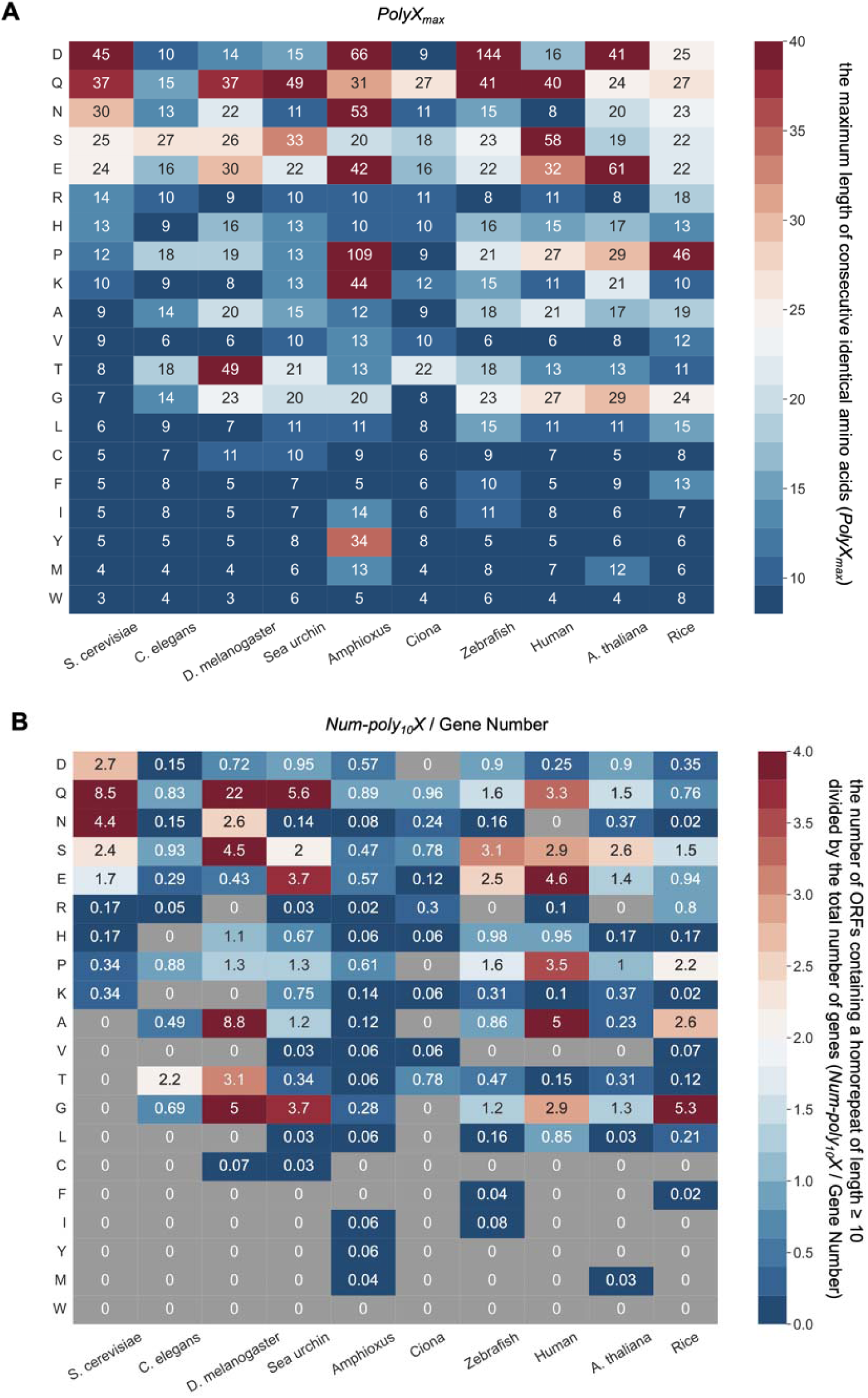
PolyX occurrence across diverse species. **A)** Heatmap of PolyX_max_ values (maximum homorepeat length for each amino acid) across multiple species. **B)** Heatmap of Num-Poly_10_X values across the same species. Each value represents the number of proteins containing ≥10-residue homorepeats per amino acid, normalized by the total number of genes in the species and multiplied by 1000. Gray boxes indicate amino acids for which no homorepeats of length ≥10 were detected.

**Figure S28.**
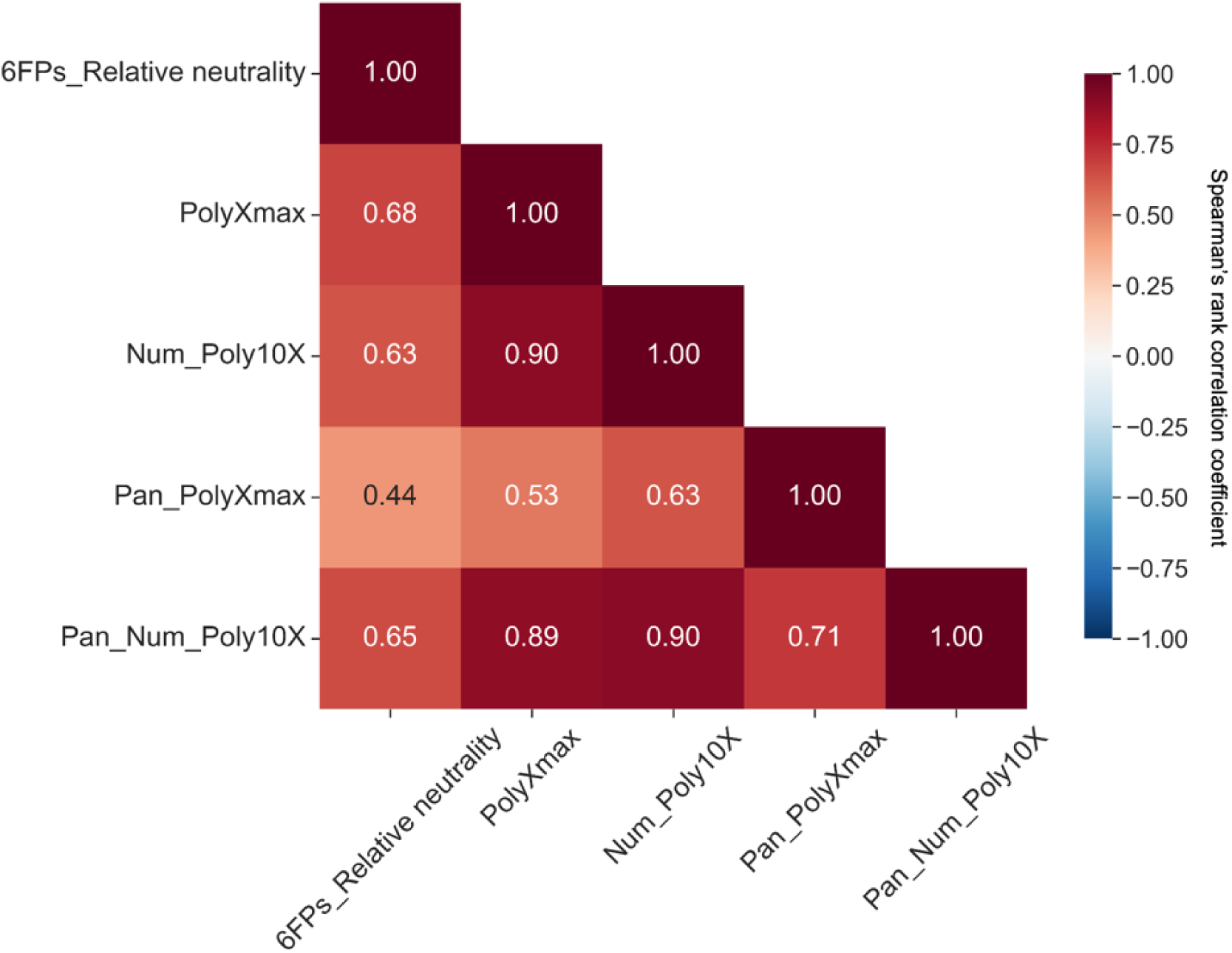
Correlation analysis between experimental Poly_10_X harmfulness patterns and proteome-wide PolyX occurrence trends. Correlation matrix comparing Poly_10_X harmfulness patterns observed across different fluorescent proteins with amino acid–level PolyX_max_ and Num-Poly_10_X values obtained from proteome-wide computational analyses. Each value represents the Spearman’s rank correlation coefficient for the corresponding pairwise comparison.

**Figure S29.**
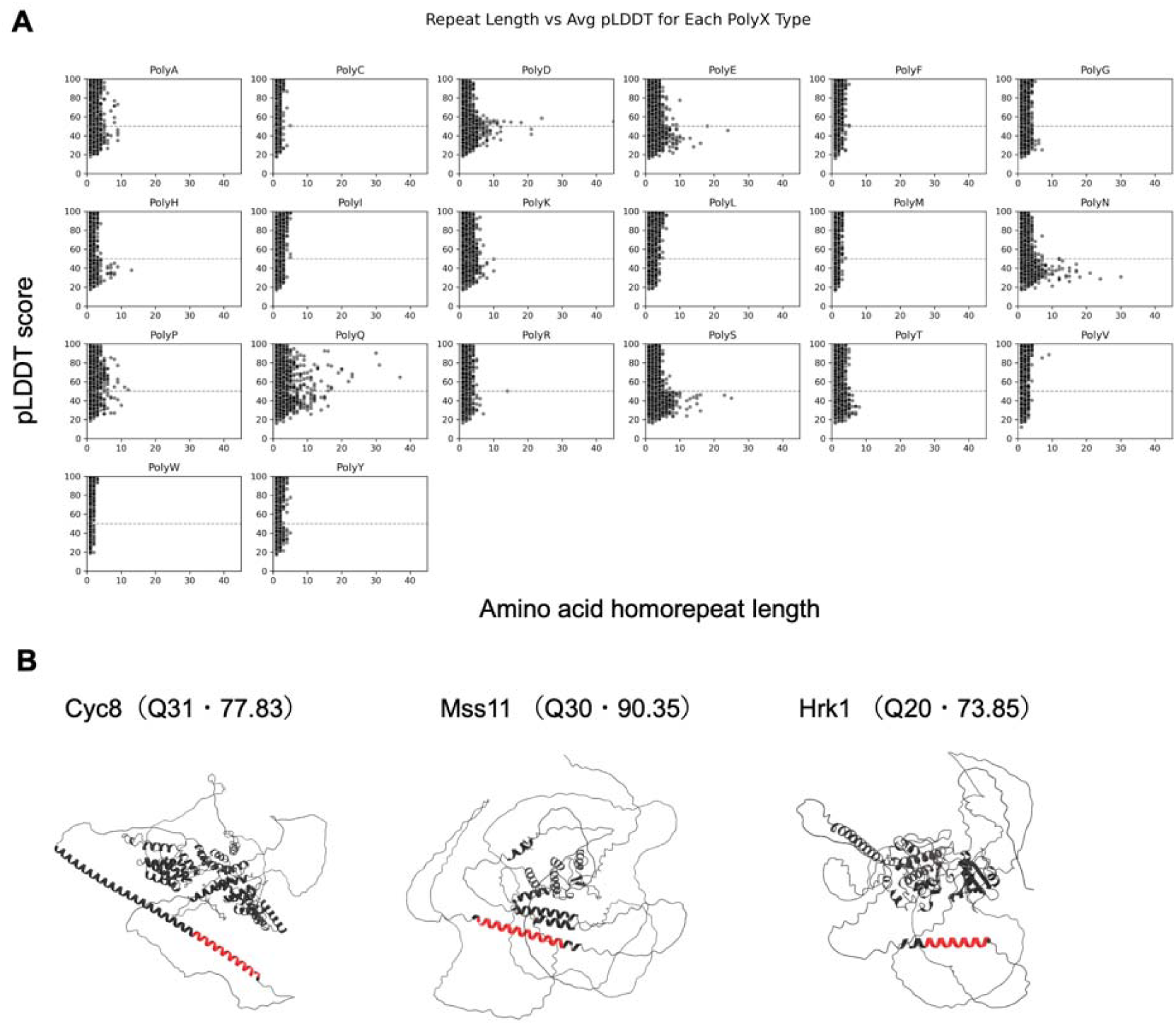
Structural context of PolyX homorepeats in the *S. cerevisiae* proteome based on AlphaFold pLDDT scores. **A)** Distribution plots showing the relationship between amino acid homorepeat length in the *S. cerevisiae* proteome and the corresponding pLDDT scores obtained from the AlphaFold structural prediction dataset (https://alphafold.ebi.ac.uk/). Each dot represents a sequence region within a protein. The pLDDT score indicates prediction confidence (100–90: very high; 90–70: confident; 70–50: low; 50–0: very low). Regions with low pLDDT scores likely correspond not only to low model confidence but also to intrinsically disordered regions (IDRs), which do not adopt a fixed 3D structure. This analysis was performed to examine whether naturally occurring PolyX regions tend to reside within well-structured protein cores or within flexible, disordered regions. For most amino acids, longer homorepeats were associated with lower pLDDT scores, indicating enrichment in unstructured regions. In contrast, PolyQ (glutamine) repeats maintained relatively high pLDDT scores even at longer lengths. **B)** Representative structures of PolyQ-containing proteins, with PolyQ segments that exhibit pLDDT ≥ 70 highlighted in red. Although these PolyQ regions show high prediction confidence, they appear to be located away from the structured protein core. This suggests that PolyX tracts, including PolyQ, are generally enriched in flexible or intrinsically disordered regions rather than in tightly structured core domains.

## References

1. Dryden, D. T. F., Thomson, A. R. & White, J. H. How much of protein sequence space has been explored by life on Earth? J. R. Soc. Interface 5, 953–956 (2008).

2. Barton, M. D., Delneri, D., Oliver, S. G., Rattray, M. & Bergman, C. M. Evolutionary systems biology of amino acid biosynthetic cost in yeast. PLoS One 5, e11935 (2010).

3. Jumper, J. et al. Highly accurate protein structure prediction with AlphaFold. Nature 596, 583–589 (2021).

4. Abramson, J. et al. Accurate structure prediction of biomolecular interactions with AlphaFold 3. Nature 630, 493–500 (2024).

5. Baek, M. et al. Accurate prediction of protein structures and interactions using a three-track neural network. Science 373, 871–876 (2021).

6. Hayes, T. et al. Simulating 500 million years of evolution with a language model. Science 387, 850–858 (2025).

7. Georgakopoulos-Soares, I., Yizhar-Barnea, O., Mouratidis, I., Hemberg, M. & Ahituv, N. Absent from DNA and protein: genomic characterization of nullomers and nullpeptides across functional categories and evolution. Genome Biol. 22, 245 (2021).

8. Koulouras, G. & Frith, M. C. Significant non-existence of sequences in genomes and proteomes. Nucleic Acids Res. 49, 3139–3155 (2021).

9. Keefe, A. D. & Szostak, J. W. Functional proteins from a random-sequence library. Nature 410, 715–718 (2001).

10. Devlin, J. J., Panganiban, L. C. & Devlin, P. E. Random peptide libraries: a source of specific protein binding molecules. Science 249, 404–406 (1990).

11. Romero, P. A. & Arnold, F. H. Exploring protein fitness landscapes by directed evolution. Nat. Rev. Mol. Cell Biol. 10, 866–876 (2009).

12. Fowler, D. M., Stephany, J. J. & Fields, S. Measuring the activity of protein variants on a large scale using deep mutational scanning. Nat. Protoc. 9, 2267–2284 (2014).

13. Diss, G. & Lehner, B. The genetic landscape of a physical interaction. Elife 7, (2018).

14. Vaishnav, E. D. et al. The evolution, evolvability and engineering of gene regulatory DNA. Nature 603, 455–463 (2022).

15. Thompson, M. et al. Massive experimental quantification allows interpretable deep learning of protein aggregation. Sci. Adv. 11, eadt5111 (2025).

16. Castro, J. F. & Tautz, D. The effects of sequence length and composition of random sequence peptides on the growth of E. coli cells. Genes (Basel) 12, 1913 (2021).

17. Neme, R., Amador, C., Yildirim, B., McConnell, E. & Tautz, D. Random sequences are an abundant source of bioactive RNAs or peptides. *Nat*. Ecol. Evol. 1, 0217 (2017).

18. Chavali, S. et al. Constraints and consequences of the emergence of amino acid repeats in eukaryotic proteins. Nat. Struct. Mol. Biol. 24, 765–777 (2017).

19. Chavali, S., Singh, A. K., Santhanam, B. & Babu, M. M. Amino acid homorepeats in proteins. Nat. Rev. Chem. 4, 420–434 (2020).

20. Jorda, J. & Kajava, A. V. Protein homorepeats sequences, structures, evolution, and functions. Adv. Protein Chem. Struct. Biol. 79, 59–88 (2010).

21. Lobanov, M. Y., Sokolovskiy, I. V. & Galzitskaya, O. V. HRaP: database of occurrence of HomoRepeats and patterns in proteomes. Nucleic Acids Res. 42, D273–8 (2014).

22. Lobanov, M. Y. & Galzitskaya, O. V. Occurrence of disordered patterns and homorepeats in eukaryotic and bacterial proteomes. Mol. Biosyst. 8, 327–337 (2012).

23. Mier, P. & Andrade-Navarro, M. A. dAPE: a web server to detect homorepeats and follow their evolution. Bioinformatics 33, 1221–1223 (2017).

24. Mier, P. & Andrade-Navarro, M. A. PolyX2: Fast detection of homorepeats in large protein datasets. Genes (Basel) 13, 758 (2022).

25. Cascarina, S. M. & Ross, E. D. Proteome-scale relationships between local amino acid composition and protein fates and functions. PLoS Comput. Biol. 14, e1006256 (2018).

26. Oma, Y., Kino, Y., Sasagawa, N. & Ishiura, S. Comparative analysis of the cytotoxicity of homopolymeric amino acids. Biochim. Biophys. Acta 1748, 174–179 (2005).

27. Oma, Y., Kino, Y., Sasagawa, N. & Ishiura, S. Intracellular localization of homopolymeric amino acid-containing proteins expressed in mammalian cells. J. Biol. Chem. 279, 21217–21222 (2004).

28. Oma, Y., Kino, Y., Toriumi, K., Sasagawa, N. & Ishiura, S. Interactions between homopolymeric amino acids (HPAAs). Protein Sci. 16, 2195–2204 (2007).

29. Gojobori, J. & Ueda, S. Elevated evolutionary rate in genes with homopolymeric amino acid repeats constituting nondisordered structure. Mol. Biol. Evol. 28, 543–550 (2011).

30. Karlin, S., Brocchieri, L., Bergman, A., Mrazek, J. & Gentles, A. J. Amino acid runs in eukaryotic proteomes and disease associations. Proc. Natl. Acad. Sci. U. S. A. 99, 333–338 (2002).

31. Faux, N. G. et al. Functional insights from the distribution and role of homopeptide repeat-containing proteins. Genome Res. 15, 537–551 (2005).

32. Qiu, Y., Xu, D., Lei, P., Li, S. & Xu, H. Engineering functional homopolymeric amino acids: from biosynthesis to design. Trends Biotechnol. 42, 310–325 (2024).

33. Courtemanche, N. Mechanisms of formin-mediated actin assembly and dynamics. Biophys. Rev. 10, 1553–1569 (2018).

34. Balhorn, R. The protamine family of sperm nuclear proteins. Genome Biol 8, 227 (2007).

35. Guhaniyogi, J. & Brewer, G. Regulation of mRNA stability in mammalian cells. Gene 265, 11–23 (2001).

36. Ito-Harashima, S., Kuroha, K., Tatematsu, T. & Inada, T. Translation of the poly(A) tail plays crucial roles in nonstop mRNA surveillance via translation repression and protein destabilization by proteasome in yeast. Genes Dev. 21, 519–524 (2007).

37. Terpe, K. Overview of tag protein fusions: from molecular and biochemical fundamentals to commercial systems. Appl. Microbiol. Biotechnol. 60, 523–533 (2003).

38. Futaki, S. et al. Arginine-rich peptides. J. Biol. Chem. 276, 5836–5840 (2001).

39. Orr, H. T. & Zoghbi, H. Y. Trinucleotide repeat disorders. Annu. Rev. Neurosci. 30, 575–621 (2007).

40. Dimitrova, L. N., Kuroha, K., Tatematsu, T. & Inada, T. Nascent peptide-dependent translation arrest leads to Not4p-mediated protein degradation by the proteasome. J. Biol. Chem. 284, 10343–10352 (2009).

41. Mizuno, M. et al. The nascent polypeptide in the 60S subunit determines the Rqc2-dependency of ribosomal quality control. Nucleic Acids Res. 49, 2102–2113 (2021).

42. Chadani, Y. et al. Intrinsic ribosome destabilization underlies translation and provides an organism with a strategy of environmental sensing. Mol. Cell 68, 528–539.e5 (2017).

43. Ito, Y. et al. Nascent peptide-induced translation discontinuation in eukaryotes impacts biased amino acid usage in proteomes. Nat. Commun. 13, (2022).

44. Haerty, W. & Golding, G. B. Low-complexity sequences and single amino acid repeats: not just ‘junk’ peptide sequences. Genome 53, 753–762 (2010).

45. Ntountoumi, C. et al. Low complexity regions in the proteins of prokaryotes perform important functional roles and are highly conserved. Nucleic Acids Res. 47, 9998–10009 (2019).

46. Kashi, Y. & King, D. G. Simple sequence repeats as advantageous mutators in evolution. Trends Genet. 22, 253–259 (2006).

47. Pelassa, I. et al. Compound dynamics and combinatorial patterns of amino acid repeats encode a system of evolutionary and developmental markers. Genome Biol. Evol. 11, 3159–3178 (2019).

48. Dorsman, J. C. et al. Strong aggregation and increased toxicity of polyleucine over polyglutamine stretches in mammalian cells. Hum. Mol. Genet. 11, 1487–1496 (2002).

49. Moriya, H. The expression level and cytotoxicity of green fluorescent protein are modulated by an additional N-terminal sequence. AIMS Biophys. 7, 121–132 (2020).

50. Moriya, H., Makanae, K., Watanabe, K., Chino, A. & Shimizu-Yoshida, Y. Robustness analysis of cellular systems using the genetic tug-of-war method. Mol. Biosyst. 8, 2513–2522 (2012).

51. Moriya, H., Shimizu-Yoshida, Y. & Kitano, H. In vivo robustness analysis of cell division cycle genes in Saccharomyces cerevisiae. PLoS Genet. 2, e111 (2006).

52. Azizoglu, A., Brent, R. & Rudolf, F. A precisely adjustable, variation-suppressed eukaryotic transcriptional controller to enable genetic discovery. Elife 10, (2021).

53. Buric, F. et al. Amino acid sequence encodes protein abundance shaped by protein stability at reduced synthesis cost. Protein Sci. 34, e5239 (2025).

54. Liu, Z. et al. Systematic comparison of 2A peptides for cloning multi-genes in a polycistronic vector. Sci. Rep. 7, 2193 (2017).

55. Fujita, Y., Namba, S. & Moriya, H. Impact of maximal Overexpression of a non-toxic protein on yeast cell physiology. (2024) doi:10.7554/elife.99572.

56. Namba, S., Kato, H., Shigenobu, S., Makino, T. & Moriya, H. Massive expression of cysteine-containing proteins causes abnormal elongation of yeast cells by perturbing the proteasome. G3 (2022) doi:10.1093/g3journal/jkac106.

57. Li, G., Ji, B. & Nielsen, J. The pan-genome of Saccharomyces cerevisiae. FEMS Yeast Res. 19, (2019).

58. Hatsuzawa, K., Tagaya, M. & Mizushima, S. The hydrophobic region of signal peptides is a determinant for SRP recognition and protein translocation across the ER membrane. J. Biochem. 121, 270–277 (1997).

59. Fauchère, J. L., Charton, M., Kier, L. B., Verloop, A. & Pliska, V. Amino acid side chain parameters for correlation studies in biology and pharmacology. Int. J. Pept. Protein Res. 32, 269–278 (1988).

60. Kyte, J. & Doolittle, R. F. A simple method for displaying the hydropathic character of a protein. J. Mol. Biol. 157, 105–132 (1982).

61. Akashi, H. & Gojobori, T. Metabolic efficiency and amino acid composition in the proteomes of Escherichia coli and Bacillus subtilis. Proc. Natl. Acad. Sci. U. S. A. 99, 3695–3700 (2002).

62. Craig, C. L. & Weber, R. S. Selection costs of amino acid substitutions in ColE1 and ColIa gene clusters harbored by Escherichia coli. Mol. Biol. Evol. 15, 774–776 (1998).

63. Wagner, A. Energy constraints on the evolution of gene expression. Mol. Biol. Evol. 22, 1365–1374 (2005).

64. Stringer, C., Wang, T., Michaelos, M. & Pachitariu, M. Cellpose: a generalist algorithm for cellular segmentation. Nat. Methods 18, 100–106 (2021).

65. Lamprecht, M. R., Sabatini, D. M. & Carpenter, A. E. CellProfiler: free, versatile software for automated biological image analysis. Biotechniques 42, 71–75 (2007).

66. Köhrer, K. & Domdey, H. [27] Preparation of high molecular weight RNA. in Methods in Enzymology vol. 194 398–405 (Academic Press, 1991).

67. Chen, S., Zhou, Y., Chen, Y. & Gu, J. fastp: an ultra-fast all-in-one FASTQ preprocessor. Bioinformatics 34, i884–i890 (2018).

68. Kim, D., Paggi, J. M., Park, C., Bennett, C. & Salzberg, S. L. Graph-based genome alignment and genotyping with HISAT2 and HISAT-genotype. Nat. Biotechnol. 37, 907–915 (2019).

69. Li, H. et al. The Sequence Alignment/Map format and SAMtools. Bioinformatics 25, 2078–2079 (2009).

70. Pertea, M. et al. StringTie enables improved reconstruction of a transcriptome from RNA-seq reads. Nat. Biotechnol. 33, 290–295 (2015).

71. Robinson, M. D., McCarthy, D. J. & Smyth, G. K. edgeR: a Bioconductor package for differential expression analysis of digital gene expression data. Bioinformatics 26, 139–140 (2010).

72. Xu, S. et al. Using clusterProfiler to characterize multiomics data. Nat. Protoc. 19, 3292–3320 (2024).

